# STIE: Single-cell level deconvolution, convolution, and clustering in spatial transcriptomics by aligning spot level transcriptome to nuclear morphology

**DOI:** 10.1101/2023.12.17.572084

**Authors:** Shijia Zhu, Naoto Kubota, Shidan Wang, Tao Wang, Guanghua Xiao, Yujin Hoshida

## Abstract

In spot-based spatial transcriptomics, spots that are of the same size and printed at the fixed location cannot precisely capture the actual randomly located single cells, therefore failing to profile the transcriptome at the single-cell level. The current studies primarily focused on enhancing the spot resolution in size via computational imputation or technical improvement, however, they largely overlooked that single-cell resolution, i.e., resolution in cellular or even smaller size, does not equal single-cell level. Using both real and simulated spatial transcriptomics data, we demonstrated that even the high-resolution spatial transcriptomics still has a large number of spots partially covering multiple cells simultaneously, revealing the intrinsic non-single-cell level of spot-based spatial transcriptomics regardless of spot size. To this end, we present STIE, an EM algorithm that aligns the spatial transcriptome to its matched histology image-based nuclear morphology and recovers missing cells from up to ∼70% gap area between spots via the nuclear morphological similarity and neighborhood information, thereby achieving the real single-cell level and whole-slide scale deconvolution/convolution and clustering for both low- and high-resolution spots. On both real and simulation spatial transcriptomics data, STIE characterizes the cell-type specific gene expression variation and demonstrates the outperforming concordance with the single-cell RNAseq-derived cell type transcriptomic signatures compared to the other spot- and subspot-level methods. Furthermore, STIE enabled us to gain novel insights that failed to be revealed by the existing methods due to the lack of single-cell level, for instance, lower actual spot resolution than its reported spot size, the additional contribution of cellular morphology to cell typing beyond transcriptome, unbiased evaluation of cell type colocalization, superior power of high-resolution spot in distinguishing nuanced cell types, and spatially resolved cell-cell interactions at the single-cell level other than spot level. The STIE code is publicly available as an R package at https://github.com/zhushijia/STIE.

## Introduction

Recent rapid development in spatially resolved transcriptomics has enabled systematic characterization of cellular heterogeneity while retaining spatial context^1–6^, which is crucial for mapping the structural organization of tissues and facilitates mechanistic studies of cell-environment interactions. The available spatially resolved transcriptomic techniques have three major categories^7^: i) *in situ* hybridization technologies, such as seqFISH^8^ and MERFISH^9^; ii) *in situ* sequencing technologies, such as STARmap^3^ and FISSEQ^10;^ and iii) *in situ* capturing technologies, such as spatial transcriptomics (ST)^4^, SLIDE-seq^2^, ZipSeq^11^, and HDST^12^. The first two categories achieved cellular or subcellular resolution by *in situ* visualization of predefined RNA targets, but the limited multiplex capacity hurdles them with only hundreds to thousands of genes profiled. In contrast, the third category captures transcripts *in situ*, followed by sequencing readout *ex situ*, thereby avoiding the limitations of direct visualization and allowing for an unbiased analysis of the complete transcriptome. The *in situ* capturing/barcoded area is referred to as “spot”. To date, the most predominant commercially available technique is 10X Visium, which is a spot-based spatial transcriptomics further developed from the ST technology in 2018. It is a higher throughput, more sensitive, and less custom protocol than the other alternative approaches, and very importantly, it supports both fresh frozen and formalin-fixed paraffin-embedded (FFPE) tissues.

One of the primary technological limitations of the spot-based spatial transcriptomics resides in the low resolution of the spot, which often covers multiple cells. The current 10X Visium spot size is 55 μm in diameter, with the number of spot-covering cells ranging from 1 to 30, depending on the biological tissue^13^. To enhance the resolution, BayesSpace^14^ proposed using Bayesian statistics to draw the neighborhood structure in spatial transcriptomic data and increase the resolution to the subspot level. Specifically, each ST spot was segmented into nine subspots, and each Visium spot was segmented into six subspots. Another method, called xfuse^15^, used a deep generative model to combine spatial transcriptome with histological image data, which can characterize the transcriptome of anatomical features in the micrometer resolution. Meanwhile, some other methods, such as stLearn^16^ and SpaGCN^17^, have integrated the histology image to spatially smooth gene expression and improve clustering with contiguous spatial patterns. In addition to the computational enhancement, the resolution of spatial transcriptomics technique is improving itself, e.g., the new version of 10X Visium is expected to be released at the end of 2022 under the name of “Visium HD”, with a much higher resolution (as reported in 10X Genomics Xperience 2022).

However, the spot resolution in cellular size does not equal single-cell level. Differing from single-cell sequencing technologies, the spots of the same size and fixed position cannot precisely capture the actual different-sized and randomly located single cells, which can still partially cover multiple cells in close proximity simultaneously, making spot-based spatial gene expression essentially fail to achieve the single-cell level regardless of spot size. This lack of single-cell level cannot be resolved by the current methods via only enhancing spot resolution computationally or technically, hindering the characterization of individual cellular spatial organization and cell-type-specific gene expression variation in spatial transcriptomics. In addition, on the current Visium platform, 4992 spots of 55 μm in diameter are printed on the glass slide of 6500×6500 µm^2^ in area, leaving 100 μm distance between two spot centers and gene expression unmeasured from up to ∼70% area 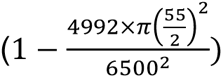. However, most current methods cannot fill this gap, resulting in a considerable loss of information. BayesSpace segmented the spot into subspot but cannot fill the gap area by only using the information from the spot, while xfuse combines spatial transcriptome with histological image to impute super-resolved expression within and between spots, although it is not the single-cell level. Third, compared with the fresh frozen tissue, the staining image from the FFPE tissue better preserves the cellular morphology, therefore facilitating more accurate cell/nucleus segmentation. However, the existing methods do not well utilize the matched image generated along with the expression profile by failing to take into account the cell segmentation, morphology features, and spatial location of every single cell captured by the image.

To this end, we present STIE, which integrates spatial transcriptome data with matched histology image-based nucleus morphological information, to bridge the gap between spot resolution and single-cell level. By assuming that the cell type proportion of one spot inferred from spatial transcriptome and nucleus morphological features is similar, STIE uses the Expectation-Maximization (EM) algorithm to jointly model spatial transcriptome and nucleus morphology to find their consensus. Given cell-type transcriptomic signatures, STIE performed the single-cell level deconvolution and convolution for the low- and high-resolution spots, respectively; when provided no cell-type transcriptomic signatures, STIE performs single-cell level clustering. Moreover, STIE fills the gap between spots by recovering missing cells via neighborhood information. As such, STIE enables the real single-cell level and whole-slide scale analysis for spot-based spatial transcriptomics.

## Results

### Single-cell resolution does not equal single-cell level

On the 10X Visium platform, the tissue sections are placed onto glass slides, where the spots of the same size are printed at fixed coordinates with barcoded Reverse Transcript (RT) primers. During tissue permeabilization, mRNA molecules diffuse vertically down to the solid surface and hybridize locally to the RT primers within the spot *in situ. The* cDNA-mRNA complexes are further extracted for library preparation and the sequencing readout. Different from scRNA-seq, which precisely captures each cell individually and profiles the gene expression of the whole cell, the spot at the predefined position is highly likely to partially cover multiple cells in close proximity simultaneously. However, these single cells cannot be resolved via only reducing spot size into cellular size computationally or technically, i.e., the single-cell resolution is not the single-cell level. We first took two recently proposed computational enhancement methods of single-cell resolution, BayesSpace^14^ and xfuse^15^, as illustrative examples. We run BayesSpace on a 10X Visium mouse brain hippocampus data (Fig. 1a). BayesSpace enhanced the ST resolution by segmenting one Visium spot into six subspots, and computationally impute the gene expression for each subspot. Because of smaller size, the subspot covers the cell in a small fraction, but due to the fixed size and location, it covers multiple cells simultaneously, therefore failing to achieve the single-cell level. Likewise, we investigated xfuse on a human breast cancer spatial transcriptomics data^4^ (Supplementary Fig. 1), which enhanced the spot resolution into even higher resolution, but it is still not the single-cell level due to the fixed size and location of the single enhancement unit.

**Fig. 1.**
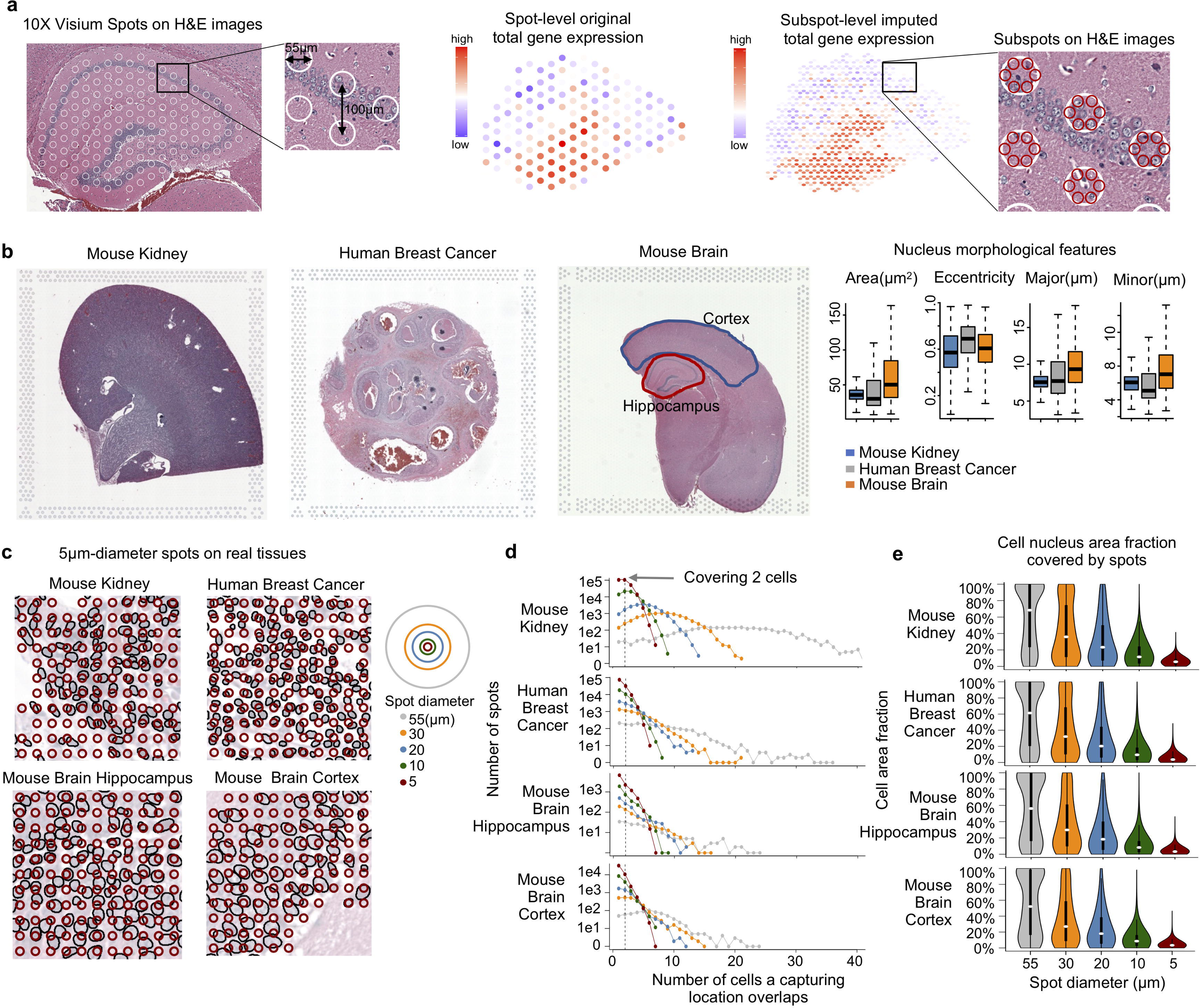
Single-cell resolution is not single-cell level. **a**, Computational resolution enhancement cannot achieve single-cell level. Illustration of the spot layout on the mouse brain hippocampus 10X Visium FFPE spatial transcriptome (left), the original gene expression summary at the spot level (middle), the BayesSpace-imputed gene expression summary at the subspot level along with the enlarged subspots on the H&E image (right). In the enlarged area, the white circle represents the original spot, and the red circle represents the enhanced subspot. **b-f,** Systematic evaluation of spatial relationship between single cells and spots via simulation of high-resolution spots from real 10X Visium spatial transcriptomics FFPE data. **b**, Nuclear morphological feature distribution of mouse kidney, mouse brain and human breast cancer. **c**, Examples of simulated high-resolution spots on the real nuclear segmentation. The brown circle represents the high-resolution spot with 5 μm in diameter, and the black circle represents the nucleus in the real tissue. **d**, The frequency of spots covering different numbers of cells, where the x-axis is the cell count covered by one single spot, and the y-axis represents the spot frequency. Only the spot that covers cells was considered. **e**, The distribution of the cell area fraction covered by the spots in different diameters.

In addition to the computational imputation-based resolution enhancement, the resolution of spatial transcriptomics technique is also increasing itself. For example, the current spot size of 10X Visium is 55 μm in diameter, while the new version with enhanced resolution, called “Visium HD”, is expected to be released at the end of 2022. However, it still holds a high chance to partially cover multiple cells. Next, we performed the cell segmentation on the spatial transcriptomics-matched H&E image and systematically investigated the spatial relationship between single cells and spots of different resolutions. We first explored a real high-resolution breast cancer HDST dataset^12^, which captures RNA from histological tissue sections on a dense, spatially barcoded bead array. However, similar to most of the current spatial transcriptomic techniques, HDST only supports the frozen tissue section, which is still more challenging to segment nuclei compared to FFPE (Supplementary Note 2, Supplementary Fig. 2). Accurate nuclear segmentation is crucial here to understand the spatial relationship between spots and single cells. Therefore, we simulated the high-resolution spots from the real 10X Visium spatial transcriptome FFPE data. Three tissues, mouse brain, mouse kidney, and human breast cancer, were investigated to account for different cell sizes of cell types (Fig. 1b). In addition to 55 μm, we simulated spots with diameters of 30 μm, 20 μm, 10 μm, and 5 μm to cover the whole tissue (Fig. 1c), where the 5 μm diameter is even smaller than the average cell diameter of real tissues. With no loss of generality, we still left the gap area between spots. At 55 μm, each spot covered a median of 5-19 cells (Fig. 1d; mouse kidney: 19, mouse brain cortex: 6, mouse brain hippocampus: 5, human breast cancer: 6), while with the spot size reduced to 5 μm, the cell count largely decreased to 1-2 cells (Fig. 1d; mouse kidney: 2, mouse brain cortex: 1, mouse brain hippocampus: 1, human breast cancer: 1). Likewise, the cell area fraction covered by spots also largely decreases with the reduced spot size. At 55 μm, the cell area demonstrates a bimodal distribution, with the majority of cells fully covered by the spot, while at 5 μm, only ∼5% of the cell area is covered by the spot by the median (Fig. 1e). Of note, the high-resolution spot covered 1-2 cells by the median, but there were still a large number of spots covering multiple cells. Even at 5 μm, many spots covered at least two cells, which was consistent across the three tissues (Fig. 1d, mouse kidney: 62%, mouse brain hippocampus: 38%, mouse brain cortex: 29%, and human breast cancer: 37%). As such, one spot, even at a very small size, is still highly likely to cover multiple cells simultaneously with each cell covered at a very small fraction, making the spot still capture the mixed gene expression of cell types.

Thus, the above gap between spot resolution and single-cell level raises the new computational challenges: “single-cell deconvolution” and “single-cell convolution” for low- and high-resolution spots, respectively. Differing from the traditional cell type deconvolution^18–20^ that estimates cell-type proportion for spots, both single-cell deconvolution and convolution aim to resolve single cells from spatial transcriptomics spots, find their spatial location and identify their cell types, where the difference resides in that single-cell deconvolution refers to splitting the large low-resolution spot into single cells, while single-cell convolution refers to assembling the small pieces covered by adjacent high-resolution spots into one complete single cell.

### Overview of the STIE workflow

To address the above challenges, we proposed an EM algorithm, called STIE, which used two pieces of information (Fig. 2), spot-level gene expression and matched histology image-based nucleus segmentation. STIE took as input (but is not limited to) the nucleus segmentation implemented in DeepImageJ^21^, called the “Multi-Organ Nucleus Segmentation model”. It was proposed by the “Multi-Organ Nucleus Segmentation Challenge^22^” to train the generalized nucleus segmentation model in H&E stained tissue images. Next, the nuclear location along with the nuclear morphological features were extracted for every single nucleus.

**Fig. 2.**
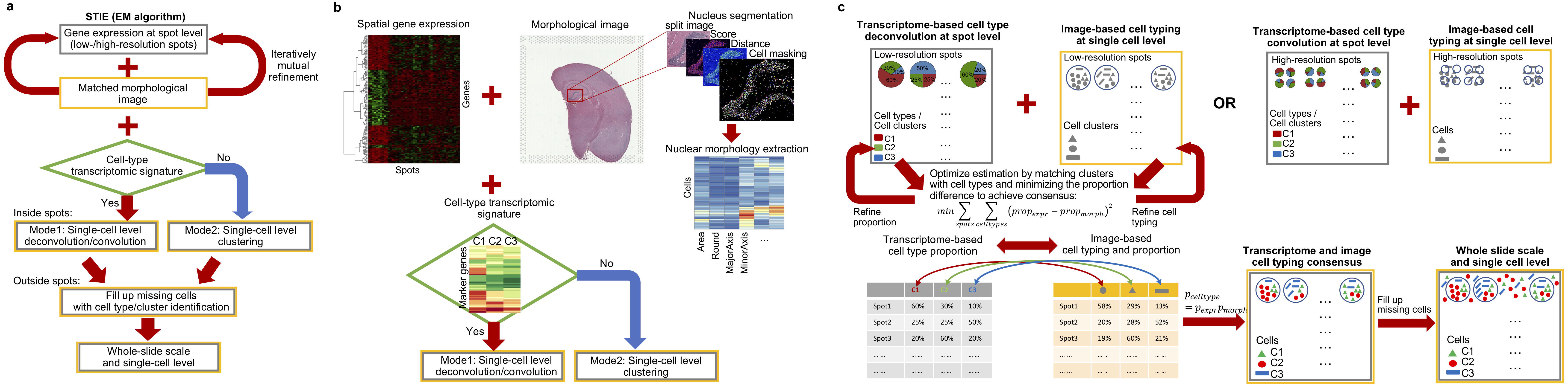
Workflow of STIE by integrating spot-level spatial transcriptome with nuclear morphology. **a**, Overview of STIE workflow. **b**, Two modes of STIE: 1) single-cell deconvolution/convolution for low-/high-resolution spots and 2) single-cell clustering, i.e., signature-free single-cell deconvolution/convolution. **c**, The flowchart of STIE single-cell deconvolution/convolution.

The rationale of the STIE is that, given the cell type transcriptomic signature, the cell type proportion can be estimated from the gene expression of each spot, while a similar proportion can also be calculated from the histology image by segmenting cells/nuclei, extracting cell/nucleus morphological features, clustering single cells into groups, locating every single cell and calculating the cell cluster proportion covered by each spot. The gene expression-based and nuclear morphology-based cell type/cluster proportions (Supplementary Note 1) are theoretically the same, motivating us to model gene expression and nucleus morphology jointly, i.e., borrow information from nuclear morphology-based cell typing to refine gene expression-based cell type proportion estimation, followed by the other way around. Accordingly, we proposed STIE, an EM algorithm to model their joint likelihood as main loss function along with their difference in cell-type proportion estimation as penalty, thereby mutually drawing information and gradually refining each other to achieve their consensus. Information borrowing was implemented in Equation (4) and Equation (8) of the M-step (see Method), which utilized both gene expression and nuclear morphology to refine the morphological model and the gene expression model, respectively.

STIE has two modes: 1) Single-cell deconvolution/convolution: Given the cell type transcriptomic signature, STIE deconvolutes/convolutes the low-/high-resolution spots into single cells. The transcriptomic signature can be defined from the existing single-cell atlas of different tissues^23–25^. 2) Single-cell clustering (Signature-free single cell deconvolution/convolution): Given no cell type transcriptomic signature, STIE can perform clustering at the single-cell level. The clustering algorithm is built on deconvolution/convolution: given the number of clusters and the initial value of the clustering transcriptomic signature, STIE iteratively refines the clustering signature-based single-cell deconvolution/convolution and re-estimates the signature until the iteration converges. STIE has two hyperparameters, *λ* and *γ*, representing the shrinkage penalty for nuclear morphology and the area surrounding the center of the spot, respectively. *λ* penalizes the difference between gene expression- and nuclear morphology-predicted cell type proportion, and accordingly, we chose *λ* for STIE via a heuristic strategy (Supplementary Information & Supplementary Fig. 3) to tweak the weights of contribution between gene expression and nuclear morphology to the final model fitting. Meanwhile, by modulating *γ*, STIE aims to find the group of cells that are covered by the spot and can best fit the gene expression of the spot. Based on the observation on different real datasets, we set γ=2.5x reported spot diameter for the 10X Visium datasets used in this paper (Supplementary Information), which gives the best concordance between the prediction and true spatial gene expression. Given no nuclear morphological features prespecified, STIE also selects the nuclear morphological features along with *λ* based on a grid search over the combinations of morphological features and *λ* (Supplementary Information & Supplementary Fig. 4).

At last, the cells covered by the spot are assigned cell types based on both spot gene expression and cellular morphology, which also include those cells located outside the reported spot size but inside the enlarged spot area, *γ*. Moreover, the cells located outside *γ* are assigned cell types using the cellular morphology and gene expression from their adjacent enlarged spot area. As such, STIE filled up the missing cells from the gap area between spots. We performed the simulation analysis by generating different combinatorial scenarios between nuclear morphology and spatial gene expression with either high or low noise. On the simulated dataset, we confirmed the convergence of STIE deconvolution (Supplementary Fig. 5) and evaluated the running time (Supplementary Fig. 6), which scales well with number of cell types and spots and is not significantly influenced by the number of marker genes. Moreover, we compared STIE with the other deconvolution methods, SPOTlight^18^, DWLS^19, 26^, Stereoscope^27^, RCTD^20^, Tangram^28^ and BayesPrism^29^, which only reply on gene expression (Supplementary Fig. 3). STIE outperformed all the others due to incorporation of nuclear morphology as additional information.

### Integration of gene expression and nuclear morphology enables single-cell level deconvolution/convolution in spatial transcriptomics

We investigated the mouse brain 10X Visium FFPE spatial transcriptome (Fig. 3a-h, see the Methods section), and deconvoluted the hippocampal region based on single-cell RNA-seq (scRNA-seq)-derived mouse brain hippocampus cell-type transcriptomic signatures^30^. Three methods were first compared, SPOTlight^18^, BayesSpace^14^, and STIE, representing three categories of deconvolution methods, direct deconvolution, imputation-based deconvolution and our novel nuclear morphology and gene expression integration-based single-cell level deconvolution. First, SPOTlight^18^ works on each spot as bulk gene expression and deconvoluted it into cell types at different proportions, achieving resolution at the spot level (Fig. 3a). Second, BayesSpace^14^ segmented each spot into subspots and used the information from spatial neighborhoods to impute subspot gene expression. Following up with SPOTlight, enhanced resolution of cell type deconvolution was achieved at the subspot level (Fig. 3b). Third, by integrating the image with gene expression, STIE does not only identify the cell type for each single cell within the spot (Fig. 3c middle) but also recovers the cell outside the spot (Fig. 3c right), making the whole-slide-wide cell type deconvolution at the highest single-cell level (Fig. 3c left). Meanwhile, STIE learned the distribution of morphological features for each cell type, making the result more interpretable (Fig. 3j). STIE primarily considered two categories of nuclear morphological features (but not limited to): size (Area, Major, Minor, Width, Height, Feret, and Perimeter) and shape (Round and Circular). The best features along with *λ* were selected by a grid search via evaluating the fitting of gene expression and nuclear morphology simultaneously (Supplementary Information; Supplementary Fig. 7-10). By comparing with the ground truth^25, 30^ (Fig. 3i), STIE accurately found the position of all four cell types at the single-cell level. The other two methods found the cell types at certain proportions, but SPOTlight found high proportions of CA1 at all spots and failed to see CA2. Moreover, the estimated locations of cell types largely deviate from the truth. With the enhanced subspot resolution by BayesSpace, the follow-up SPOTlight more accurately estimated the positions of cell types, revealing the advantage of high resolution by BayesSpace; however, the CA2 proportion is still underestimated at the true CA2 locations but overestimated at the true CA1 locations. Besides SPOTlight, we compared five other methods, which only take gene expression as input to deconvolute cell types at spot level, including DWLS^19, 26^, Stereoscope^27^, RCTD^20^, Tangram^28^ and BayesPrism^29^ (Fig. 3d-h & Supplementary Fig. 11 for their running time and memory usage). DWLS and RCTD showed better results than SPOTlight, with positions of cell types predicted more accurately. However, one of the shortcomings of the gene expression-based deconvolution method is that it cannot account for the cell count, therefore making misleading the high proportion of cell types within spots. For example, CA1 is primarily located in the pyramidal cell layer (Fig. 3i), and the other layers have much fewer nuclei, but SPOTlight, DWLS and RCTD still suggest the existence of CA1 in those cell-sparse layers at certain proportions (Fig. 3a). Second, the estimated proportion of one cell type within each spot is often accompanied by others (Fig. 3a-b&d-h), but those locations have only one cell type as indicated by ISH (Fig. 3i), such as the spots along the band of CA1, CA2, and CA3. This co-existence may result from the underlying co-linearity of the cell-type transcriptomic signatures (Supplementary Fig. 12). STIE addressed the above shortcomings by incorporating the nuclear segmentation and morphology as orthogonal features, facilitating better distinguishing among cell types.

**Fig. 3.**
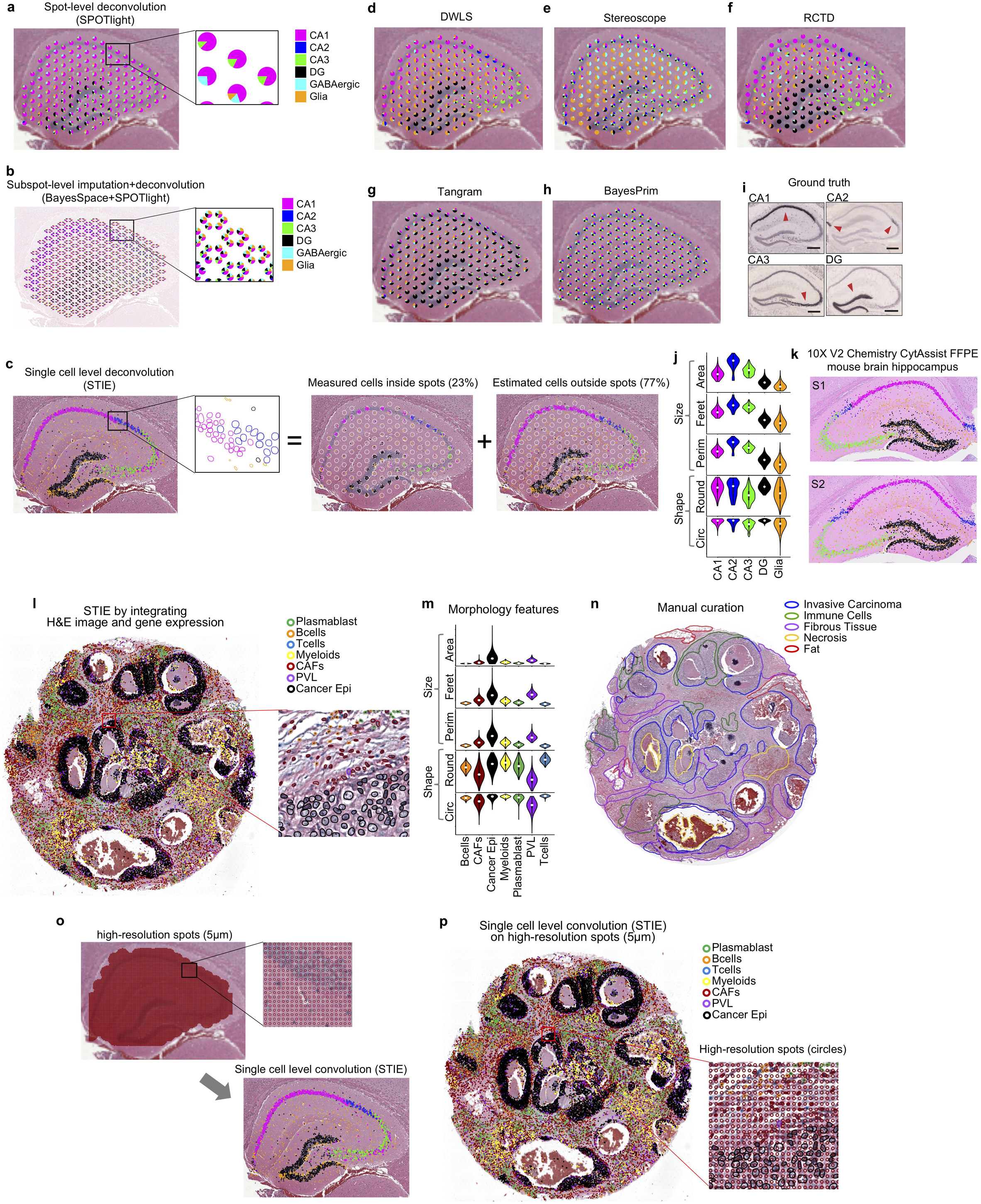
Single-cell level deconvolution in spatial transcriptomics by STIE. **a-j**, Mouse brain hippocampus 10X Visium FFPE spatial transcriptome. **a**, Spot-level cell-type deconvolution using SPOTlight. In the enlarged area, each pie chart represents the proportion of cell types for the corresponding spot. **b**, Subspot-level cell type deconvolution using BayesSpace followed by SPOTlight. In the enlarged area, each pie chart represents the proportion of cell types for the corresponding subspot. **c**, Single-cell level deconvolution by STIE (left panel), which is the aggregation of cells captured by spots (middle panel) and cells missed by spots but recovered by STIE (right panel). In the enlarged area, the circle is the cell contour, with the color representing its cell type. **d-h**, Spot-level cell-type deconvolution using DWLS (**d**), Stereoscope (**e**), RCTD (**f**), Tangram (**g**), and BayesPrism (**h**). **i**, Nuclear morphological feature distribution of each cell type learned by STIE. **j**, Ground truth of mouse brain hippocampus cell types by In Situ Hybridization (ISH): CA1 (*Mpped1*), CA2 (*Map3k15*), CA3 (*Cdh24*) and DG (*Prox1*). The arrowhead indicates high expression. The figure is adapted from ref^30^. **k**, Single-cell level deconvolution by STIE on 10X Visium V2 Chemistry CytAssist FFPE spatial transcriptomics of two consecutive mouse brain hippocampus sections: section 1 (up panel) and section 2 (bottom panel). **l-n**, Human breast cancer 10X Visium FFPE spatial transcriptome. **l**, Single-cell level deconvolution by STIE. **m**, Nuclear morphological feature distribution for each cell type. **n**, Manually annotated human breast cancer pathological regions. **o-p**, Single-cell convolution for the simulated high-resolution spatial transcriptomics data of the mouse brain hippocampus (**o**) and human breast cancer (**p**) using spots with 5 μm in diameter.

To further investigate the accuracy and applicability, we tested STIE deconvolution on the recently released 10X Visium V2 Chemistry CytAssist mouse brain hippocampus FFPE spatial transcriptomics datasets. On both consecutive sections (Fig. 3k), STIE deconvoluted the single cells accurately that are highly consistent with the ground truth. To investigate the influence of selection of marker genes on STIE deconvolution, we used SPOTlight to construct a new transcriptomic signature and use it to rerun STIE deconvolution. Although the new signature has only 39% overlapping genes with the original, STIE still deconvoluted the single cells accurately conferring high concordance with those using original signatures (Supplementary Fig. 13a-c).

Moreover, we randomly sampled the marker genes from the original signature, and rerun STIE deconvolution to investigate its consistency with that using full signatures. As expected, the concordance is reduced with decreasing marker genes. However, with half marker genes, the concordance is still higher than 80%, except the CA2 that has few cells (Supplementary Fig. 13d). In addition, we investigated the influence of image alignment to the STIE deconvolution, we gradually shifted the image to make it misaligned to the spot, rerun STIE, and evaluated its concordance with that under accurate alignment (Supplementary Fig. 14). As expected, the concordance is slowly reduced with the increasing misalignment of cells with the spots. However, STIE still has 88% and 84% concordance with the accurate alignment, even if the cells are misplaced by a half spot and a whole spot, respectively. A slight misalignment, 10% spot size, does not significantly influence the cell typing (98% concordance).

In addition, we investigated the human breast cancer 10X Visium FFPE spatial transcriptome dataset (Fig. 3l-n). Based on the human breast cancer scRNA-seq-derived cell type transcriptomic signature^31^, we used STIE to deconvolute spots into single cells with cell typing, including Cancer and Normal epithelial cells, Cancer-associated fibroblast (CAF), Perivascular-like cells (PVL), Myeloids, B-cell and T-cell (Fig. 3l), along with the morphological features characterized for each cell type (Fig. 3m). The majority of cell types demonstrated distinct morphological features. Among immune cells, Myeloid is relatively easier to distinguish, showing a much larger nucleus size, whereas T-cell and B-cell are well-known to resemble morphologically, but STIE still estimated a slightly larger nucleus area of B-cell than T-cell, which is consistent with the previous observation^32^. We compared our result with the manual annotation by the pathologist (Fig. 3n). Despite different resolutions, our single-cell deconvolution still shows a high resemblance to the annotated regions. On this dataset, we also tested the other deconvolution methods at the spot level, SPOTlight, DWLS, Stereoscope, RCTD, Tangram, and BayesPrism, to deconvolute the spot into cell type proportions (Supplementary Fig. 15). Overall, DWLS, Stereoscope, and RCTD predicted the distinct proportions of CancerEpithelial across spots, and the high-proportion spots tend to enrich for the tumor area, while SPOTlight, Tangram, and BayesPrism predicted more identical CancerEpithelial across all spots, at low, moderate, and high proportions, respectively. We specifically compared their predicted tumor area with the manual annotation (Supplementary Fig. 16a) by evaluating the ARI and R2 (Supplementary Fig. 16b-c). STIE gives the best concordance (ARI=0.64, R2=0.64), which is slightly higher than SPOTlight (ARI=0.54, R2=0.60), RCTD (ARI=0.57, R2=0.6) and DWLS (ARI=0.57, R2=0.59), and much higher than Stereoscope (ARI=0.26, R2=0.13), Tangram (ARI=0.38, R2=0.33), and BayesPrism (ARI=0.4, R2=0.35).

With the technical advancement, the spot resolution is also increasing itself, such as the 10X Visium HD expected to be released at the end of 2022. The high-resolution spot covering multiple cells at a small fraction can be similarly modeled using STIE. To test this hypothesis, we simulated the high-resolution spot mouse brain hippocampus spatial transcriptome data (Fig. 3o; 5 μm in diameter) by using three pieces of information from the real low-resolution spot (Fig. 1a left; 55 μm in diameter), including the nuclear spatial location, STIE-deconvoluted cell types (Fig. 3c) and the associated cell-type transcriptomic signature. Next, we used STIE to convolute the small pieces of cells from adjacent spots into single whole cells (Fig. 3o bottom) and found that the convoluted cell types were highly consistent with the true cell types by deconvolution. Likewise, we also simulated the high-resolution spot spatial transcriptomic data from the real human breast cancer low-resolution spot data (Fig. 3l) and observed similarly high consistency between convoluted cell types (Fig. 3p) and their original truth (Fig. 3l). Despite no real high-resolution spot spatial transcriptomics FFPE data available so far, the above facts support the capability of STIE in convoluting small pieces of cells into single whole cells resulting from the high-resolution spot spatial transcriptomic data.

### STIE enhances spatial transcriptomic clustering to single-cell level

Given cell type transcriptomic signatures, we deconvoluted/convoluted spots into single cells. As opposed, we checked if the cell type transcriptomic signature can be reversely estimated given cell typing (see Methods). We investigated the mouse brain hippocampus and human breast cancer (Fig. 4a-b, Supplementary Fig. 17) and found that all of our learned signatures correlated most with their corresponding scRNA-seq-derived signatures. Moreover, we used the non-negative least square (NNLS) to deconvolute our estimated signatures based on the scRNA-seq-derived signatures. Again, all of our learned signatures showed high proportions of the corresponding cell types. DWLS was used as an independent method and observed similar proportions. Despite small variation between the original scRNA-seq and the learned spatial transcriptomic signatures, the above results largely corroborated that we could estimate the single-cell level transcriptomic signature from the spot-level spatial transcriptome.

**Fig. 4.**
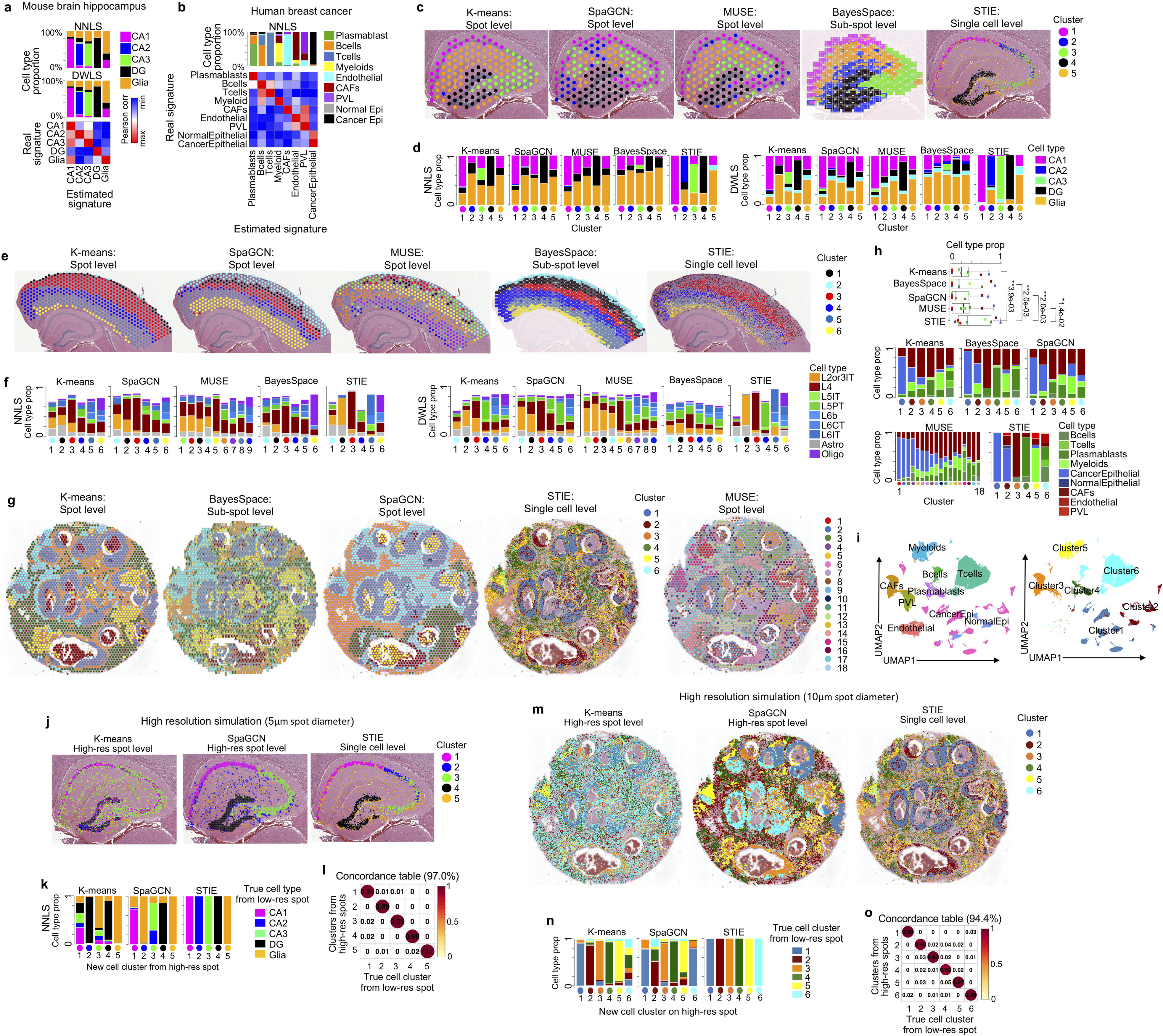
Single-cell level clustering in spatial transcriptomics by STIE. **a-b**, Cell type specific transcriptomic signature learning from 10X Visium mouse brain hippocampus FFPE (**a**) and 10X Visium human breast cancer FFPE (**b**). **c&e&g**, Spot-level clustering by K-means, SpaGCN, MUSE, subspot-level clustering by BayesSpace and single-cell-level clustering by STIE on 10X Visium FFPE mouse brain hippocampus (**c**), mouse brain cortex (**e**) and human breast cancer (**g**). **d&f&h**, Cell type deconvolution of spot-, subspot-, and single-cell-level clustering-derived CAGE in the mouse brain hippocampus (**d**), mouse brain cortex (**f**), and human breast cancer (**h**). For the mouse brain cortex, the cell types in the transcriptomic signature, which are not cortex layers and have small proportions, are not shown in the barplot. **i**, The UMAP plot of human breast cancer scRNA-seq data from 26 primary tumors^31^. The top panel is the original cell typing of 10,060 single cells, and the bottom panel is the subset of cells that are mapped to the six STIE clusters. **j&m**, Spot-level clustering by K-means (left), SpaGCN (middle), and single-cell-level clustering by STIE (right) on the simulated high-resolution spot spatial transcriptome data of the mouse brain hippocampus (**j**) and human breast cancer (**m**). **k&n**, Cell type deconvolution of spot- and single-cell-level clustering-derived CAGE in the mouse brain hippocampus (**k**) and human breast cancer (**n**). **l&o**, The consistency table of single-cell clusters between the simulated high-resolution spot-based STIE clustering and the original low-resolution spot-based STIE clustering as ground truth of the mouse brain hippocampus (**c**) and human breast cancer (**g**).

Furthermore, we similarly estimated signatures from the whole transcriptome besides signature genes. On this basis, we implemented the STIE clustering algorithm when given neither cell typing nor transcriptomic signatures: we iteratively re-estimate the clustering transcriptomic signature and refine the clustering signature-based single-cell deconvolution/convolution until the iteration converges (see Methods). We tested this clustering algorithm on the mouse brain hippocampus (Fig. 4c). As comparison, we also tested the other clustering methods on this dataset, including K-means that relies on only gene expression, SpaGCN^17^ and MUSE^33^ that also utilize the histological image, and BayesSpace at the sub-spot level (Fig. 4c; Supplementary Fig. 18-21). At k=5, STIE clustering was also close to the ground truth and matched the cell types of CA1, CA2, CA3, DG, and Glia in STIE deconvolution. K-means accurately uncovered the clusters of CA1, CA3, DG, and Glia with CA2 missed. SpaGCN roughly found the CA1, DG, and Glia clusters. MUSE automatically determined the number of clusters k=5 and found the CA1, DG, and Glia clusters. At the sub-spot level, BayesSpace clustering is more delicate with higher resolution, but it still failed to find the CA2 cluster even at a higher number of clusters (Supplementary Fig. 20). Furthermore, we calculated the average gene expression of each cluster (referred to as CAGE) and deconvoluted it based on the scRNA-seq-derived cell type transcriptomic signatures as another angle to evaluate the resolution of clustering methods (Fig. 4d). The rationale is that if the cluster is single-cell level and all cells within the cluster belong to the same cell type, the deconvolution will demonstrate a high proportion of single-cell type for each cluster; otherwise, it will be the saturated proportions of multiple cell types. As shown in Fig. 4d, the CAGEs by other methods at spot-level or subspot-level all demonstrated a mixture of cell types. In particular, all of them showed a high proportion of Glia, suggesting that most spot/subspot clusters mix with Glia cells, which may mislead the biological interpretation of the cluster. In contrast, the CAGE by STIE (i.e., the STIE-estimated whole transcriptomic signature for single-cell clusters) recovers all cell type signatures at the single-cell type level, i.e., STIE found the matched clusters for all cell types, CA1, CA2, CA3, DG, and Glia, and each cluster gives the high proportion of its matched cell type by the cell type deconvolution. Likewise, we evaluated STIE clustering on the 10X CytAssist dataset of mouse brain hippocampus (Supplementary Fig. 22), which accurately uncover the clusters of CA1, CA2, CA3, and DG largely matching the ground truth.

Moreover, we applied different clustering methods to the 10X Visium mouse brain cortex spatial transcriptome (Fig. 4e & Supplementary Fig. 23-26). At k=6, we found clear layers by K-means, SpaGCN, BayesSpace, and STIE. Specifically, K-means at the spot level has thinner cluster 1 (black), cluster 2 (cyan), and cluster 6 (yellow), but thicker cluster 3 (red), while SpaGCN and BayesSpace showed more even clusters, especially cluster 1-3, but SpaGCN identified a much thicker cluster 6 (yellow) than the others. MUSE also identified 6 clusters automatically, but its clusters are more blurred than the other methods. STIE clustering takes the K-means at spot level as initial values, so it largely resembles K-means, but STIE showed more expanded cluster 1 (black) and cluster 6 (yellow). We further checked the cell type deconvolution based on the scRNA-seq signatures of the mouse brain cortex by Allen Brain Atlas^25^. Despite different patterns by NNLS and DWLS, both spot-level K-means, SpaGCN, MUSE, and subspot-level BayesSpace showed saturated proportions of cortex layers, whereas STIE demonstrated a significantly higher proportion for the layers from L2 to L6 (Fig. 4f), which is also concordant with the consecutive spatial location of clusters. In addition, we tested STIE clustering on the mouse brain cortex 10X V2 Chemistry CytAssist FFPE datasets (Supplementary Fig. 27). On both section 1 and section 2, we observed the clear layers along with the good concordance between the CAGE and the scRNA-seq transcriptomic signatures. Further, to evaluate the impact of nuclear morphology on the STIE accuracy, we compared the above FFPE sections with the fresh frozen tissue section, which may not well preserve the morphology. We run STIE clustering at the single-cell level on the 10X Visium fresh frozen human dorsolateral prefrontal cortex^34^, which grouped the single cells into clear layers that resemble the manual annotation (Supplementary Fig. 28). By downgrading the single cells into spots, the clusters gave a moderate concordance with the manual annotation (ARI=0.30). However, STIE clustering on the human dorsolateral prefrontal cortex demonstrates worse performance and weaker robustness on the selection of nuclear morphological features, as compared to the 10X CytAssist FFPE mouse brain cortex (Supplementary Fig. 28-29). This is largely due to the low image quality of the fresh frozen section and the inaccurate nuclear segmentation (Supplementary Fig. 30), corroborating the importance of nuclear morphology to STIE in the accurate single cell clustering.

Third, we investigated the 10X Visium human breast cancer FFPE spatial transcriptome (Fig. 4g & Supplementary Fig. 31-34). Overall, all K-means, BayesSpace, SpaGCN, and STIE demonstrated clear clusters, where the clustering at k=6 demonstrates the best concordance with the STIE deconvolution (Fig. 3l). By comparing with the manual annotation, their clusters are largely consistent with the tumor region (cluster 1 of K-means with R2=0.63, cluster 1&2 of BayesSpace with R2=0.49, cluster 1&2 of SpaGCN with R2=0.58, and cluster 1&2 of STIE with R2=0.64; Supplementary Fig. 35). Although MUSE identified 18 clusters automatically, it also uncovers clusters partially matching the tumor regions well (cluster 1-4 with R2=0.38). By checking the cell type deconvolution, cluster 1 and cluster 3 in the spot/subspot level clustering showed relatively higher proportions of CancerEpithelial and CAFs, respectively, while the other clusters all mixed with different cell types, especially CAFs. In contrast, STIE clusters showed much higher cell type proportions (Fig. 4h & Supplementary Fig. 36), therefore matching well with the known cell types: cluster 1 (14.3% out of all cells) for CancerEpithelial, cluster 2 (21.3%) for CancerEpithelial, cluster 3 (14.2%) for CAF, cluster 4 (35.8%) for Plasmablasts, cluster 5 (10.2%) for Myeloids, and cluster 6 (4.2%) for mixed T-cells, B-cells and Plasmablasts. Of note, different from the deconvolution (Fig. 3l), STIE clustering identified two clusters for the tumor regions (cluster 1 and cluster 2), where cluster 2 was only located at the bottom tumor and the boundary of the other tumor regions. We aligned the clusters to ∼100,000 human breast cancer single cells from 26 primary tumors^31^ (Fig. 4i & Supplementary Fig. 37). SCINA^35^ was used to assign cell clusters based on the clustering-derived transcriptomic signature. A total of 60.7% of cells were identified with known clusters assigned (Fig. 4i right), matching 6 cell types: CancerEpithelial, CAF, Plasmablasts, Myeloids, T-cells, and B-cells (Fig. 4i left). We found a group of closely neighboring clouds for cluster 2 (brown) in the big category of CancerEpithelial, which is separated from cluster 1 (steel blue), revealing a putative subtype of breast cancer. The CAGE of cluster 2 comprises a mixture of CAF and CancerEpithelial (Fig. 4h). It may implicate the epithelial-mesenchymal transition (EMT)^36^, which is associated with tumor progression and migration.

STIE has shown good clustering performance on the current spot size, i.e., 55 μm in diameter. Next, we tested whether STIE can also achieve single-cell level clustering on the high-resolution spot. We simulated the high-resolution spot spatial transcriptome from the 5 clusters in the mouse brain hippocampus (Fig. 4c) and checked whether spot-level K-means, SpaGCN, MUSE, and single-cell-level STIE can recover those 5 clusters. We did not test the subspot level clustering, since the spot was even smaller than the cell size, with no need for further segmentation. Consequently, the K-means clusters on high-resolution spots are highly mixed together (Fig. 4j left), which heavily deviate from the truth (Fig. 4c right) and are even worse than K-means on low-resolution spots (Fig. 4c left). Consistently, the cell type deconvolution of CAGEs revealed that different cell types were mixed in all K-means clusters and that one cell type was lost (Fig. 4k left & Supplementary Fig. 38). These facts posed an even more difficult challenge for the K-means on the high-resolution spot. Compared to the K-means, SpaGCN performed much better clustering on high-resolution spots (Fig. 4j middle), which accurately found the region of CA1, CA3, and DG, but still missed CA2. The cell type deconvolution of CAGEs also recovers those four cell types despite slightly mixing with others (Fig. 4k middle), but two clusters were found for Glia with CA2 regions missed. However, as opposed to K-means and SpaGCN, STIE recovered all 5 clusters accurately, and all CAGEs were composed of single-cell clusters at a very high proportion (Fig. 4j-k). Specifically, the single cell clusters from the high-resolution spots are highly concordant with those from the low-resolution spots (Fig. 4l). A similar observation was also made for the simulated high-resolution spot data on human breast cancer (Fig. 4m-o & Supplementary Fig. 39). We also tested MUSE on the simulated high-resolution spatial transcriptomics data, which automatically determined 50 clusters for the mouse brain hippocampus and 159 clusters for the human breast cancer (Supplementary Fig. 40). The clusters are highly mixed and difficult to compare with the ground truth. Together, STIE can perform single-cell clustering for both low-and high-resolution spot spatial transcriptomics data, which outperforms spot-/subspot-level clustering in terms of both resolution and associated CAGEs.

### STIE reveals single-cell level insights in spatial transcriptomics

As discussed above, differing from the single-cell gene expression profiling, the spot cannot always capture the complete cells, resulting in loss of information from cells at the spot boundary or contamination of cells from outside. Therefore, the first question is raised: does the gene expression captured by the spot derive from all cells located in the spot, partially from cells at the center of the spot, or even more cells outside the spot? Since STIE deconvolutes whole-slide-wide single cells with both cell typing and spatial information retained, we can check different areas surrounding the spot center and their covering cells. The area that shows the best concordance between the spot gene expression and its covering cell typing can provide the answer to the actual area captured by the spot. We referred to the size of that area as the “*bona fide* spot size” to distinguish it from the reported spot size. Accordingly, we fit the STIE model by varying the *bona fide* spot sizes with the spot center fixed and checked the model fitting between gene expression and cell type deconvolution. In the mouse brain hippocampus 10X Visium spatial transcriptome, the area in 2x-3x spot diameter demonstrated the best model optimization with the least root-mean-square error (RMSE) (Fig. 5a). The analysis was repeated on the 10X Visium human breast cancer FFPE spatial transcriptome, and a consistent observation was made (Fig. 5b). Moreover, we investigated the new released 10X Visium V2 chemistry CytAssist datasets from two mouse brain replicates (Fig. 5c), and consistently observed the minimum RMSE around 2.5x spot diameter. Although a small variation was observed among datasets, the above facts suggest that the spot may capture even more gene expression from the spot surroundings, further motivating us to reevaluate the resolution of the spot. In STIE, we introduced the hyperparameter γ to represent the *bona fide* spot size, and we have set γ=2.5x reported spot diameter (55 μm) for all above analyses.

**Fig. 5.**
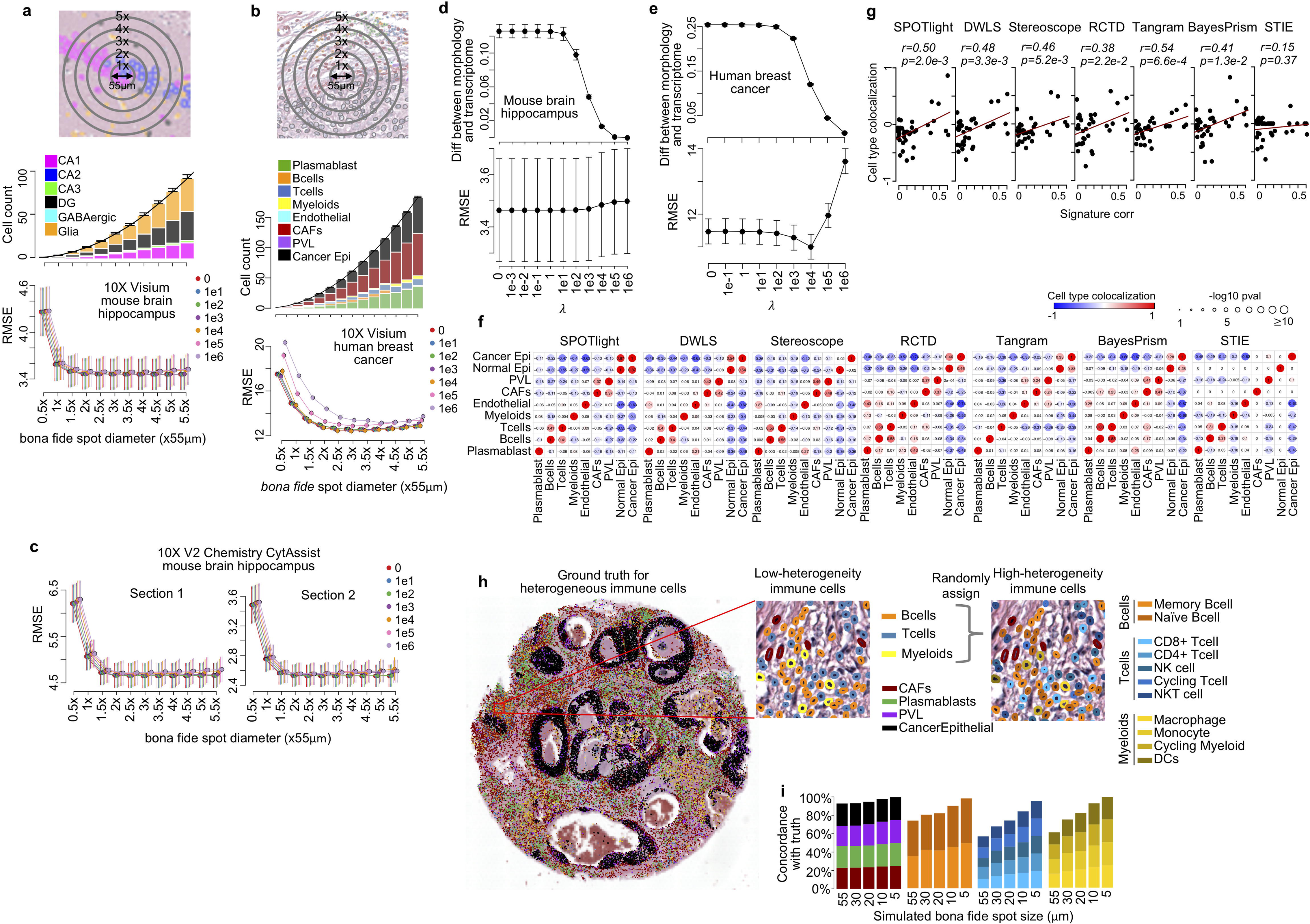
Relevant questions in spatial transcriptomics addressed by the single-cell level deconvolution. **a-b**, Identification of the *bona fide* area captured by spots in the mouse brain hippocampus (**a**) and human breast cancer (**b**). The x-axis represents the putative size of the *bona fide* area measured by the spot in the unit of a regular 10X Visium spot size (55 μm). The top panel represents the chart of *bona fide* spot size in the real tissue. The middle panel represents the cell count (y-axis) in the spot area (x-axis); the bottom panel represents the RMSE by fitting the STIE model using the cells within the corresponding area indicated by the x-axis. **c**, Identification of the *bona fide* area captured by spots in the 10X Visium V2 Chemistry CytAssist mouse brain hippocampus. **d-e**, The evaluation of image contribution to the cell type deconvolution in human breast cancer (**d**) and mouse brain hippocampus (**e**). The top panel represents the difference between cell-type proportions estimated from the spot gene expression and the nuclear morphological features; the bottom panel represents the RMSE of gene expression fitting. The x-axis represents the value of *λ* in Formula (7). **f**, Heatmap of the correlation between the cell type proportion within spots by SPOTlight, DWLS, Stereoscope, RCTD, Tangram, BayesPrism, and STIE. **g**, The association between transcriptomic signature similarity and cell-type colocalization by SPOTlight, DWLS, Stereoscope, RCTD, Tangram, BayesPrism, and STIE. **h-i**. High-resolution spots along with STIE holds the premise to distinguish nuanced cell types. **h.** Random assignments of Memory Bcell or Naïve Bcell to the Bcell; CD8+ Tcell, CD4+ Tcell, NK cells, Cycling Tcell, or NKT cell, to the Tcell; and Macrophage, Monocyte, Cycling Myeloid, or DCs to the Myeloid. **i**. The barplot represents the concordance of STIE deconvoluted/convoluted single cells with the simulation ground truth (**h**). The x-axis represents the simulated spot diameter, and the y-axis represents the concordance. The color refers to the cell type in the legend.

Second, unlike the traditional cell-type deconvolution method, STIE integrates nuclear morphological information with gene expression. Therefore, we wanted to investigate whether the nuclear morphology can really contribute to the deconvolution. According to Equation (7) (Methods), the parameter *λ* penalizes the difference between morphology- and transcriptome-based prediction. The extremely large *λ* renders the prediction entirely rely on nuclear morphology, which is first confirmed by the consecutively diminished difference between morphology- and transcriptome-based prediction along the increasing *λ* (Fig. 5d-e top). Therefore, we gradually attribute more to the morphology by increasing the shrinkage penalty *λ* and check the model fitting. Consequently, the RMSE slightly decreased and reached a minimum between 1e+03 and 1e+04 in the human breast cancer spatial transcriptomics dataset (Fig. 5e bottom), indicating the *λ* value of the best balanced contribution between transcriptome and morphology. Afterwards, the RMSE sharply increased with the extremely large *λ* value, suggesting that the optimized model cannot fully reply on the nuclear morphology. It is worth noting that the nuclear morphology has been incorporated into the STIE model even at *λ* =0 (Equation 9) when determining the single cell type. In 10X Visium and V2 Chemistry CytAssist mouse brains (Fig. 5d bottom & Supplementary Fig. 41), we all observed the flat RMSE at the beginning followed by its elevation at the large *λ*, suggesting the best model has been achieved at *λ*=0. The difference between STIE deconvolution at *λ*=0 and other methods relying on only gene expression (Fig. 3a-h) supports the additional contribution of nuclear morphology to the accurate cell typing. The similar difference can also be observed from the comparison between STIE clustering at *λ*=0 and K-means clustering on the CytAsssist mouse brain cortex (Supplementary Fig. 27). These facts corroborated the coordinated contribution of gene expression and nuclear morphology to cell typing in the spatial transcriptome.

Third, we wanted to investigate cell type deconvolution-based cell type colocalization. The cell type proportion can be estimated within each spot given the gene expression. Subsequently, the correlation between the predicted cell-type proportions has been widely used to measure cell type colocalization as a clue of cell type interaction^27, 37, 38^. Here, we compared STIE with the other six methods that take gene expression as input and deconvolute cell types at spot level. Similar to Stereoscope, we calculated for each method the Pearson correlation between cell type proportion within spots as a measure of cell type colocalization (Fig. 5f). Overall, the transcriptomic signature-based methods suggested more cell type colocalization with more saturated and pronounced values, whereas STIE was more conservative (Fig. 5f & Supplementary Fig. 42). Specifically, all methods found a negative correlation between CAF and CancerEpithelial, indicating that CAF spatially dissociates from the tumor cell area, which is consistent with the gross histological distributions of fibrous tissue area and invasive carcinoma foci in the original H&E image (Fig. 3n). In addition, a negative correlation between immune cells and CancerEpithelial was observed by all seven methods, suggesting less immune cell infiltration into tumor cell areas, which can be further related to the efficacy of immuno-oncology therapy. However, by checking the cell type transcriptomic signatures, we found that signature similarity was significantly correlated with cell type colocalization by all methods except STIE (Fig. 5g), revealing a potential co-linearity bias within transcriptomic signatures for the methods relying only on gene expression. For instance, the original tissue slide is limited to the cancer area and may not contain “normal epithelia” histologically. However, due to the high correlation of the signatures between CancerEpithelial and NormalEpithelial (*r=0.61* with *p=5.1e-130*), DWLS, SPOTlight, and RCTD all suggested their high colocalization. In contrast, STIE identified very few NormalEpithelial cells by utilizing the nuclear shape to further distinguish them from the CancerEpithelial, even though they are largely similar in the transcriptome. Therefore, STIE suggests no colocalization between NormalEpithelial and CancerEpithelial in this dataset. In addition, the misidentification of colocalization may also result from the lack of sparsity in statistical inference or failure to characterize cell counts by gene expression-based methods. For example, the cell type with an extremely small proportion is still kept in one spot, and the high proportions of cell types are still estimated for those cell-sparse areas (Fig. 3a-h). These facts implied the benefits of accurate cell typing at single-cell level via exploiting both gene expression and nuclear morphological information, which facilitates less biased downstream exploration.

At last, it would be very useful for digital pathology if STIE could distinguish more nuanced cell types, for instance, the heterogeneous T and B cells that are morphologically similar but transcriptomically different. Intuitively, the current low-resolution spot covers multiple whole cells simultaneously, making it almost infeasible to distinguish two nuanced T or B cells covered by the same spot. However, this could be potentially addressed by the upcoming high-resolution spot-based spatial transcriptomics, which is corroborated by the following simulation analysis. Based on the STIE-deconvoluted Tcell, Bcell, and Myeloid from the human breast cancer low-resolution spatial transcriptomics, we randomly assigned them the new nuanced immune cell type: Memory Bcell or Naïve Bcell to the Bcell; CD8+ Tcell, CD4+ Tcell, NK cells, Cycling Tcell, or NKT cell, to the Tcell; and Macrophage, Monocyte, Cycling Myeloid, or DCs to the Myeloid. The randomly assigned cell types were used as ground truth in the following investigation (Fig. 5h). We used the scRNA-seq data^31^ to rebuild a new highly-heterogeneous cell-type transcriptomic signature, and further, simulated spatial transcriptomics datasets with spot diameters of 55 μm, 30 μm, 20 μm, 10 μm, and 5 μm, respectively, for which the gene expression of each spot was simulated based on the fraction of its covering cell types and the corresponding high-heterogeneous cell type transcriptomic signatures. Of note, the spot size and gene expression are different between simulated datasets, while the pathology image and nuclear morphology remain the same. Using the high-heterogeneity cell type signature, we run STIE on each simulation to investigate if we can recover those nuanced cell types. The accuracy of low-heterogeneity cell types, including CAFs, Plasmablasts, PVL and Cacner Epithelial, is consistently high with small variation across spot sizes (Fig. 5i). As opposed, the accuracy of high-heterogeneity immune cells demonstrated a high variation, and with the spot size becoming smaller, the accuracy is largely improved. At 5μm, the nuanced immune cells are convoluted very accurately. The high-resolution spot still covers multiple cells partially, but it is less likely to completely cover two cells compared to the low-resolution spot, so that the cell type could be determined by the spot that captures the largest fraction of the cell and gives the highest probability. Thus, the high-resolution spot along with STIE single-cell convolution provides a higher power in distinguishing nuanced cell types.

### STIE enables the investigation of spatially resolved cell-cell interactions

Given the STIE-obtained single cells with spatial information retained, we further investigated the spatially resolved cell-cell interaction in the tissue. We use STIE to re-estimate the cell-type whole-transcriptomic signature from human breast cancer spatial transcriptomics and take it as input for CellChat^39^, a toolkit and a database of interactions among ligands, receptors, and their cofactors, to quantitatively infer the cell-cell interaction. We investigated the interactions at *λ*=1e3 and *λ*=1e4 for STIE, respectively, given their similar measurements in the gene expression and nuclear morphology fittings (Supplementary Fig. 10). By further aggregating the ligand-receptor into pathways, we found that, in both settings, the interactions of extracellular matrix (ECM) pathways, such as COLLAGEN and FN1, predominantly occur between CAF, CancerEpithelial, and PVL, as well as the immune pathways, such as SPP1 and MIF, between immune cells, CAF and CancerEpithelial (Fig. 6a & Supplementary Fig. 43). We spatially mapped the interaction strength to the cell pairs under a cutoff of cell-cell distance that may permit the potential cell-cell interaction. The COLLAGEN pathway was taken as an illustrative example (Fig. 6b). The outgoing signal is sent mainly from CAFs with a widespread distribution on the tissue (Fig. 6b left), while the incoming signal is received locally by the CancerEpithelial at the tumor region and the PVL at the tumor boundary, confirming that the spatially resolved interaction is driven not only by the ligand-receptor interaction strength but also by the cell abundance and spatial location. Accordingly, we incorporated the information of the neighboring cell pairs into the cell-cell interaction strength (Fig. 6c left) and obtained the global spatial cell-cell interaction strength (Fig. 6c right), which further suggests that spatial intercellular interactions primarily occur among CAFs and CancerEpithelial cells in this tissue.

**Fig. 6.**
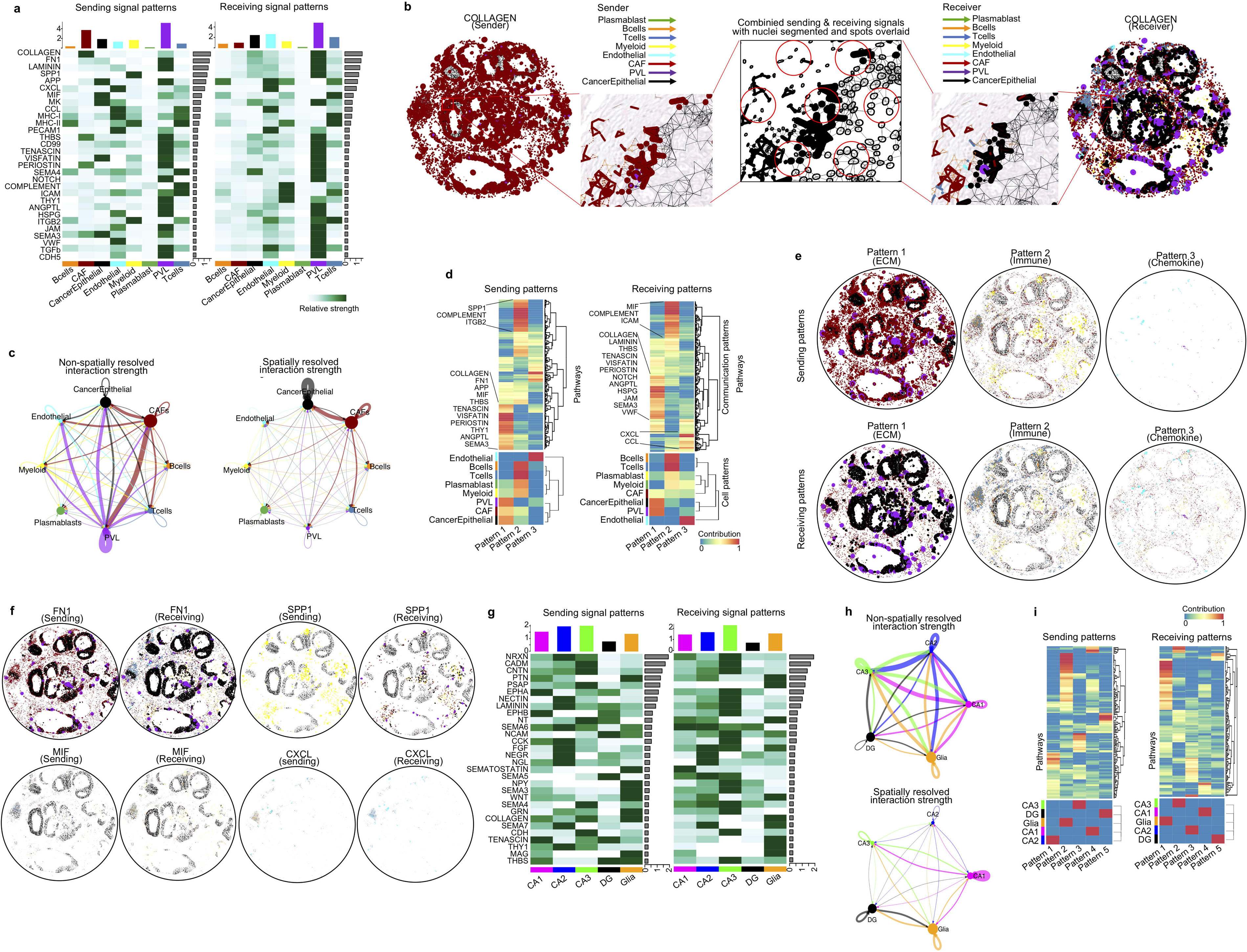
Cell-cell communication with spatial information retained. **a-f**, The spatially resolved cell-cell interaction in human breast cancer at *λ*=1e3 for STIE. **a,** Heatmap of the cell-cell interaction strength of pathways in sender cell types (left) and receiver cell types (right). In sender and receiver heatmaps, the bar plots on top and right represent the total amount of cell-cell interaction within the cell type and pathway, respectively. **b**, Cell-cell interaction in the COLLAGEN pathway. The color of the edge is indicated by the sender cell type on the left panel and the receiver cell type on the right panel, respectively, and the thickness is proportional to the interaction strength. The middle panel is the combined sending and receiving signals with nuclear segmentation and spots overlaid (The black color of edges and circles does not represent cell types). **c**, The non-spatially resolved interaction strength between cell types (left) and the total amount of spatially resolved cell-cell interaction strength (right), which is the strength of interaction (left) times the number of cell pairs at proximity. The size of the dot is proportional to the number of cell types, and the edge thickness is proportional to the interaction strength (left) and the amount of interaction (right). **d**, Heatmap of the contribution of pathways (top) and cell types (bottom) to the pattern of senders (left) and receivers (right). The enriched pathways in each pattern further overlapped with the top 30 strongest pathways (in Fig. 6a) and are listed beside the heatmap. **e**, The sender and receiver cell-cell interaction patterns across the tissue. The color of the edge is indicated by the sender cell type and the receiver cell type in the sender and receiver patterns, respectively. **f**, Cell-cell interactions in the FN1, SPP1, MIF and CXCL pathways. **g-i**, Spatial cell-cell interactions in the mouse brain hippocampus. **g,** Heatmap of the cell-cell interaction strength of pathways in sender cell types (left) and receiver cell types (right). **h**, The non-spatially resolved interaction strength between cell types (top) and the total amount of spatially resolved cell-cell interactions (bottom). **i**, Heatmap of the contribution of pathways (top) and cell types (bottom) to the pattern of senders (left) and receivers (right). In all above analyses, only cell-cell interactions within 3μm were counted to consider the potential autocrine and paracrine signaling.

To simplify the interpretation of complex intercellular interactions, we used non-negative matrix factorization (NMF) implemented by CellChat to identify the global interaction patterns beyond only exploring individual pathways. The sending and receiving patterns are learned independently and do not necessarily correspond to each other. We uncovered three sending and three receiving patterns of signaling pathways, respectively (Supplementary Fig. 44-45 & Fig. 6d left and right), revealing how the signaling pathways coordinate to drive communication for certain sender cell types and respond to incoming signals for certain target cell types. First, a large portion of ECM signaling among CAFs, CancerEpithelial, and PVL is characterized by communication between sending pattern 1 and receiving pattern 1, which represent multiple pathways, including but not limited to COLLAGEN, FN1, THBS, TENASCIN, PERIOSTIN, and ANGPTL. Second, an immune signaling pattern among T-cells, B-cells, Plasmablast, Myeloids, CAFs, and CancerEpithelial is characterized by sending pattern 2 and receiving pattern 2, representing multiple pathways, including SPP1, MIF, COMPLEMENT, ITGB, and ICAM. Third, a weak pattern among Endothelial, CAFs, Myeloids, and Plasmablast was also uncovered, characterized by receiving pattern 3 with the most minor pathways, e.g., CXCL and CCL. Mapping patterns to the spatial location provides quick insight into their global abundance (Fig. 6e), with pattern 1 being the most predominant, pattern 2 less pronounced, and pattern 3 the sparsest for both sending and receiving signals. At *λ*=1e4, two patterns were uncovered, corresponding to the dominant pattern 1 (ECM signaling) and pattern 2 (immune signaling) at *λ*=1e3, with the weak pattern 3 eliminated (Supplementary Fig. 46-48).

There are multiple lines of biological evidence supporting the intercellular interaction in human breast cancer. For example, the stiffened collagenous stroma has been reported as a promoter of malignant transformation and a poor prognostic feature of breast cancer^40, 41^. Accordingly, the collagen signaling between cancer cells and stromal cells in our analysis may indicate a potential impact of the molecular interaction on patient prognosis. Fibronectin (FN) is another component of the ECM that induces an EMT response and tumor progression to breast cancer cells. It is reported to be linked with collagenous bundles^40, 42, 43^, which is consistent with our data that FN signaling is activated along with collagen signaling among cancer cells, CAFs, and PVLs. Furthermore, FN contributes to the development of EMT through cooperation with signals initiated by the type-I TGFβ receptor that is also found in our data. Thus, our analysis clearly visualizes the spatial cancer-stroma association via ECM-related molecular pathways in breast cancer. In addition, our data illustrate that osteopontin (SPP1) signaling is mainly sent from myeloid cells, although osteopontin has been reported to be secreted from various cells^44^ (Fig. 6f). Interestingly, the high signal of the osteopontin pathway from myeloid cells to cancer cells was observed in tumor necrotic areas in our analysis, and many of these myeloid cells could be morphologically classified as macrophages in the H&E staining slide. Considering that osteopontin from TAMs is reported to be a poor prognostic factor in breast cancer patients^45^, the necrotic area of breast cancer tissue can be one of the hotspots of osteopontin signaling leading to poor prognosis of the patient. Moreover, differing from the intracellular MIF, extracellular MIF is reported to play a pro-oncogenic role in promoting breast cancer cell-stroma interactions^46^.

In addition to human breast cancer, we investigated the mouse brain hippocampus as another testing tissue with potentially different mechanisms. As expected, the interaction primarily occurred among the neuron-associated pathways (Fig. 6g), e.g., NRXN, CADM, CNTN, PTN, and PSAP. As supporting evidence, transcripts encoding the synaptic adhesion molecules neurexin-1,2,3 are commonly expressed in principal cells and interneurons of the mouse hippocampus, and the conditional ablation of NRXN alternative splice insertions results in differential hippocampal network activity by changing the synaptic interaction between neurons and impairment in a learning task^47^. The non-spatially resolved intercellular communication indicated connections among the majority of cell types with similar strengths (Fig. 6h top), while after considering the cell spatial location, intercellular communication between cell types became less pronounced and tended to occur more frequently within cell types (Fig. 6h bottom). The spatially resolved interaction is more reasonable because the hippocampal cell types are distributed locally and separated from each other. Consistently, five sending patterns and five receiving patterns are dedicatedly identified for five cell types, respectively (Supplementary Fig. 49-51 & Fig. 6i).

To conclude, we adapted the CellChat toolkit to fit the STIE-obtained single-cell level spatial transcriptomics so that we can explore the spatially resolved cell-cell interaction by integrating gene expression with both cell spatial information and prior knowledge of interactions between signaling ligands and receptors. Thus, the cell-cell interaction is explored at the single-cell level, holding the premise to reveal more informative and meaningful biological insights over the original spot-level spatial transcriptomics and scRNA-seq data.

## Discussion

In this paper, we used both real and simulation data, demonstrating that due to fixed position and fixed size of spots, the spot-based spatial transcriptomics essentially does not capture single cells. The single cells cannot be resolved by only enhancing spot resolution via computational imputation or technical improvement, i.e., single-cell resolution is not single-cell level. To bridge the gap, we integrated the histology image generated along with the spatial transcriptomics by aligning the gene expression to the image-based nucleus segmentation, thereby achieving the single-cell level deconvolution and convolution for the low- and high-resolution spot-based spatial transcriptomics, respectively. The STIE method relies on cell/nucleus segmentation to locate cells and morphological information to distinguish cells. The current 10X Visium supports FFPE that better preserves the cell morphological information compared to fresh frozen tissue section, therefore holding the great premise for more reliable nucleus segmentation. STIE takes as input the generalized nucleus segmentation by “Multi-Organ Nucleus Segmentation”^22^, aiming to provide broader support to different tissues. STIE also allows for the segmentation of other cells/nuclei as input. The dedicated segmentation model tailored for a specific organ may further improve the accuracy.

The resolved single cell with retaining spatial location and cell typing facilitates addressing the critical relevant questions: first, how large an area can the spot capture? We found that the area captured by the 10X Visium spot is larger than its reported spot size, i.e., ∼2x-3x reported spot diameter, which demonstrates the best concordance between spatial transcriptome and nuclear morphology. The larger spot-capturing area is supported by independent 10X Visium datasets, suggesting that the resolution of spot-based transcriptomics should be re-evaluated. Likewise, the simulation spatial transcriptomics data used in this paper may have even higher spot resolution than we reported, since we simulated the *bona fide* spot size rather than spot size. The fact that the *bona fide* spot size is larger than the reported spot size has both cons and pros: it further downgrades the resolution of the spot due to covering even more cells; however, it reduces the loss of information by capturing the gap area between spots. The integration of imaging and transcriptomics can balance the trade-off to achieve enhanced resolution and minimize the loss of information. In addition, we demonstrated that the inclusion of nuclear morphology information improves the cell type deconvolution compared with relying only on gene expression by modulating the ratio of contribution to the model between nuclear morphology and gene expression. Thus, one question is raised: if STIE can resolve the single cells and rescue the gap area from the low-resolution spot spatial transcriptomics, do we really need the high-resolution spot? Our result suggested that compared to the low-resolution spot, the high-resolution spot spatial transcriptomics combining with the STIE convolution has superior power in distinguishing the nuanced cell types, such as CD4+ and CD8+ Tcells. This is a promising advantage, which may lead to many useful applications. Given the improved single-cell level deconvolution with both spatial information and cell types, we checked the cell type colocalization. We found that the cell type deconvolution methods, which only take gene expression as input, tend to be biased by the underlying similarity among cell-type transcriptomic signatures, thereby overestimating cell type colocalization, while by incorporating nucleus morphology as orthogonal features, we better distinguish cells from their resembling transcriptome, enabling less biased cell type colocalization and misleading conclusions.

In addition to cell deconvolution, STIE can perform single-cell level clustering when given no cell-type transcriptomic signatures, i.e., signature-free single-cell deconvolution/convolution. Since the spot covers multiple cells simultaneously, the resulting cluster inevitably comprises a mixture of cell types, making the clustering average gene expression misleading due to the mixed cell-type gene expression. The same challenge can also be raised for the high-resolution spot. STIE addressed the challenge well and achieved single-cell level clustering. More importantly, the STIE clustering-derived gene expression showed the high concordance with scRNA-seq-derived cell type signatures, suggesting that STIE can extract the cell type-specific gene expression variation from the spatial transcriptomics similar to the scRNA-seq. However, the cluster can be too small to be found as significant, sometimes, such as cluster 2 (CA2) in the mouse brain hippocampus that are missed by some clustering methods, while given the transcriptomic signature, we could identify the clear cloud of CA2 by most deconvolution methods. Accordingly, signature-based cell deconvolution is recommended, especially for small groups. However, the predefined signature may not cover all possible cell types, which may also bias or misclassify the cell groups. As such, two methods should be combined to balance the trade-off and gain a better understanding.

Finally, given the deconvoluted/convoluted single cells with spatial information retained, we explored the spatially resolved cell-cell interaction at the single-cell level rather than spot level. We did not attempt to discover the ligand-receptor interaction de novo or estimate the best distance of the cell-cell interaction but instead relied on prior knowledge, which is well validated and reliable. The dominant ECM pathway interaction between CAFs and tumors was uncovered in human breast cancer, and neuron-associated pathways were found in the mouse brain hippocampus.

Taken together, STIE provides a timely and effective solution for the currently most predominant spot-based spatial transcriptomics technique to fill the gap between spot resolution and single-cell level. We expect it to make a rich set of applications in diverse domains, such as genomics, biochemistry, and clinical studies, and lead to broadly novel biological insights.

## Methods

### 10X Visium ST data preprocessing

The raw reads and images of 10X ST Visium FFPE for mouse brain, mouse kidney and human breast cancer and the processed gene-level read count matrix and image of 10X CytAssist Spatial Gene Expression (V2 Chemistry) for mouse brain section 1&2 were downloaded from www.10xgenomics.com/resources. The raw data were preprocessed using Spaceranger ver. 1.3.0 with parameters in the downloaded “web_summary.html”. We used the automatic tissue detection by SpaceRanger workflow to filter out the spots located outside tissue areas based on the histological examination. Further, considering that the contamination of background RNA other than cellular RNA may cause automatic tissue detection to falsely identify spots outside the tissue, we also performed manual annotation, to exclude the location that do not overlap tissues. For the specific area of tissue, e.g., the mouse brain cortex and mouse brain hippocampus, we used Loupe Browser ver. 5.1.0 Visium Manual Alignment to manually annotate and select the area of interest and generate the .json file for the following analysis. The spaceranger ver. 1.3.0 count command was run with the parameter “--loupe-alignment” set to be the .json file.

The processed gene-level read count matrix, histology image and nuclear segmentation of 10X Visium frozen human dorsolateral prefrontal cortex were downloaded from http://research.libd.org/globus. The nuclear morphological features were extracted using ImageJ.

## Image-based nucleus segmentation using deep learning

The ImageJ plugin DeeplmageJ^21^ with the “Multi-Organ Nucleus Segmentation” model^22, 48–50^ was used to perform nucleus segmentation. The Multi-Organ Nucleus Segmentation model was proposed by the “Multi-Organ Nucleus Segmentation Challenge^22^”, aiming to train the generalized nucleus segmentation model in H&E stained tissue images to address the challenge that nucleus segmentation algorithms working well on one dataset can still perform poorly on other datasets with variation resulting from different organs, disease conditions, and even digital scanner brands or histology technicians. The cell location and morphology features were extracted for every single cell using ImageJ ROI Manager. To reduce memory consumption, the large H&E images were split into small images of 3000×3000 pixels. To avoid cells being split on the image boundary, each split image has four 100-pixel margins overlapping with its adjacent images. The overlapped cells were checked from the four 100-pixel margins based on their location distance and cell diameter. The larger cell among overlapped cells was kept for the following analysis. The ImageJ macro was incorporated into the STIE R package to run the ImageJ plugin for each split image.

### Construction of cell-type transcriptomic signature from the scRNA-seq data

Given the single-cell data set, we used the buildSignatureMatrixMAST() R function developed by DWLS to select the marker genes and build the matrix of cell-type transcriptomic signature. The cell types in the single-cell data are expected to represent all cell types in the spatial transcriptomics. The clusters of the single cells that reveal the constituent cell types are required as input. Upon characterization of the cell types, differential expression analysis is performed to identify marker genes for each cell type. We define marker genes as genes with an FDR adjusted p-value of <0.01 (defined using the hurdle model in the MAST R package), and a log2 mean fold change >0.5. To create the final signature matrix *G*, the expression values of these chosen genes are averaged across each cell type, so that each resulting candidate matrix is an *N*×*T* matrix, where *N* is the number of genes and *T* is the number of cell types.

### STIE model

We first listed the variables used in the model (see Supplementary Information for detailed derivation):

- *N*, *C*, *T*, *S*, and *F* represent the number of genes, cells, cell types, spots, and morphological features, respectively.
- *G* = {*G_it_*}_1≤*i*≤*N*;1≤*t*≤*T*_ is a matrix of cell-type gene expression signatures, where *G_it_* is the expression of the *i-th* signature gene of the *t-th* cell type.
- *E* = {*E_is_*}_1≤*i*≤*N*;1≤*s*≤*S*_ is a matrix of gene expression from the ST, where *E_is_* is the expression of the *i-th* gene in the *s*-th spot.
- *c* = {*c_s_*}_1≤*s*≤*S*_ is the cell index on the ST histology image, where *c_s_* represents the *c*-th cell in *the s-th* spot, and *c*_s_ represents the total cell count in *the s-th* spot. The parameter *γ* is used to represent the *bona fide* area captured by the spot, which can be different from the reported spot size.
- *M* = {*M_cf_*}_1≤*c*≤*C*;1≤*f*≤*F*_ is a matrix of nuclear morphological features obtained from the ST histology image, where *M_cf_* is the value of the *f-th* morphological feature of the *c*-th cell.
- Let *q* = {*q_c_*}_1≤*c*≤*C*_ represent the hidden cell type of one single cell that generates the observed morphological features *M_c_* and gene expression *E_c_*, with *q_c_* = *t* representing the *c-th* cell that takes the cell type *t*.

We formulate that 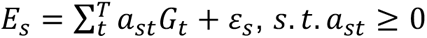 *and ε_s_*∼*N* (0, σ^2^), where *a_st_* is the non-negative regression coefficient for the *s*-th spot and the *t*-th cell type. On the other hand, we assume that the probability of the nuclear morphological feature follows a Gaussian distribution, *P* (*M_c_s__* | *q_c_s__* = *t*) ∼*N* (*μ_t_*, Σ*_t_*), where *μ_t_* and Σ*_t_* are the mean and variance of the nuclear morphological features of the *t-th* cell type, respectively. Thus, the parameter of STIE is a 3-tuple: *θ* = {*a_st_*, *μ_t_*, Σ*_t_* }.

We used the Expectation-Maximization (EM) algorithm to solve the STIE model and estimate *θ*. The Q function takes the following form:

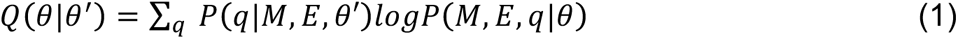

We assume that each cell is generated independently and that *E* and *M* are conditionally independent given its cell type, *q*. Thus, the Q function is rewritten as:

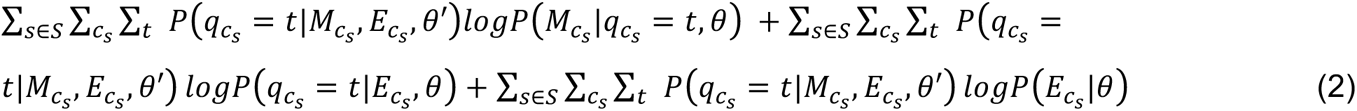

The transcriptome of cells in the same spot are profiled in bulk and observed as the same gene expression profile, i.e., *E_c_s__* = *E_s_*. Likewise, *P*(*q_i_s__* = *t*|*E_i_s__*, *θ*) of the cell *i*_s_ and *P* (*q_j_s__* = *t*|*E_i_s__*, *θ*) of the cell *j_s_* from the same spot *s* are also indistinguishable, and therefore, we obtain that for any cell *c_s_* in the spot *s*, *P* (*q_c_s__* = *t*|*E_c_s__*, *θ* = *P* (*q_c_s__* = *t*|*E_s_*, *θ*) = *a_st_*/Σ*_t_ a_st_*. So, the Q function is rewritten as:

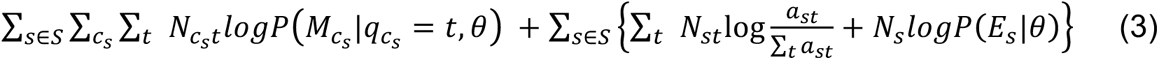

where 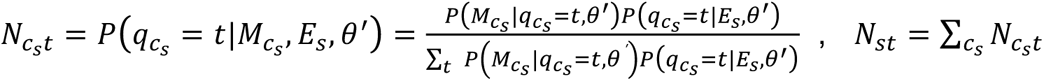, and *N_s =_* Σ*_c_s__* Σ*_t_ N_c_s_t_*.

In the M-step, we first take the derivative of the Q function (Equation 3) with respect to µ*_t_*, and Σ*_t_*, respectively. These two parameters are only in the first term, and the derivative of the second term is zero. To optimize it, we obtained the following equations:

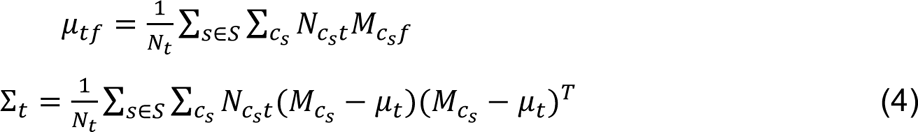

where *N_t_* = Σ_*s*∈*S*_ Σ_*c_s_*_ *N_c_s_t_*

Since the parameter *a_st_* for spatial gene expression are only in the second term of Equation (3), we optimize the second term to solve out *a_st_*. Each spot can be solved independently as following:

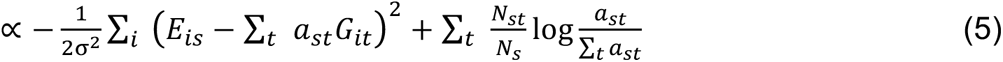

This is not a convex optimization. But according to the Gibbs’ inequality, Σ*_i_ p_i_logp_i_* ≥ Σ*_i_ p_i_logq_i_*, the right part reaches the maximum at 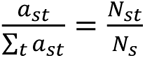. The term 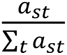 represents the cell type proportion estimated from gene expression (denoted by *Prop_t_* (*E_s_*|*θ*′)), while 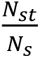 represents the cell type proportion estimated from both gene expression and nuclear morphology (denoted by *Prop_t_* (*M_s_*, *E_s_*|*θ*^’^)). This is very similar to our expectation that the proportions of cell type estimated from gene expression and that from nuclear morphology are the same (Supplementary Note 1). Therefore, we further assume that the maximum is achieved around *Prop_t_* (*E_s_*|*θ*^’^) = *Prop_t_* (*M_s_*|*θ*^’^) = *Prop_t_* (*M_s_*, *E_s_*|*θ*^’^), and our aim is to find their consensus *Prop_t_* (*M_s_*, *E_s_*|*θ^′^*) as the final estimation of cell type proportions. Accordingly, we also searched a local solution for the left part of Formula (5) in the above small area. Similar to the method DWLS^19^, we take an approximate solution of the left part via the non-negative least square, and further, add the local area derived from the right part as a constraint to restrict their solution, i.e.,

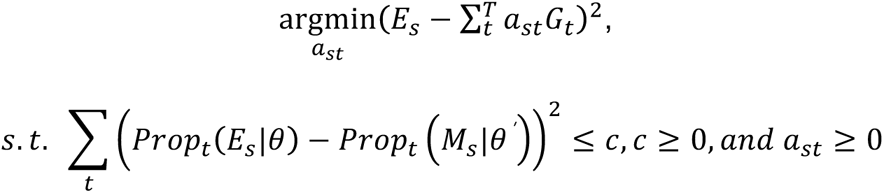

Similar to Tikhonov regularization, we use a Lagrange multiplier and rewrite the problem as follows:

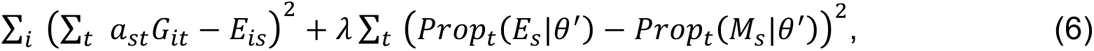

where *Prop_t_* (*E_s_*|*θ*^’^) = *a_st_*/ Σ*_t_ a_st_* and *Prop_t_* (*M_s_*|*θ*^’^) = Σ*_c_s__ P* (*q_c_s__* = *t*|*M_c_s__*, *θ*^’^) /*C_s_* represent the proportion of cell type *t* in spot *s*, which are estimated from gene expression and nuclear morphology, respectively.

Theoretically, there is a one-to-one correspondence between *c* and *λ*. Thus, we transformed Formula (5) into an easier quadratic optimization and simultaneously optimized the left and right parts of Formula (5) around *Prop_t_* (*E_s_*|*θ*^’^) = *Prop_t_* (*M_s_*|*θ*^’^).

By assuming that 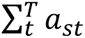 approximately equals the current estimate 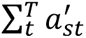, the formula can be represented in the format of inner products:

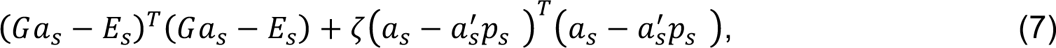

where we take 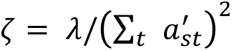 for brevity; *a_s_* = (*a*_*s*1_, …, *a_sT_*)^*T*^, representing the *T*×1 vector of regression coefficients for the *s*-th spot; *G* = (*G_it_*), representing the matrix of signature gene expression whose first index refers to the marker gene and the second index refers to the cell type; 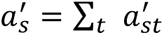, representing the sum of the current estimates of *a_s_* ; and 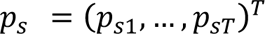 with 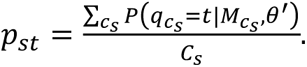.

By expanding all items and matching the standard form of quadratic linear programming,

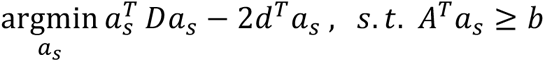

### we got that

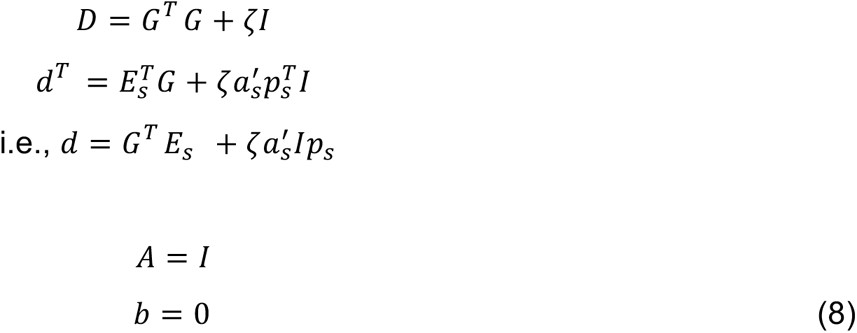

Given the standard form, we solved the quadratic linear programming using the R package “quadprog” (https://cran.r-project.org/web/packages/quadprog/index.html). The function is solve.QP(D,d,A,b).

### Filling up missing cells outside spots via spot neighborhood information

Given the STIE model with the parameters *θ* = {*a_st_*, *μ_t_*, Σ*_t_* }, the cell types were assigned as following:

First, for the cell lying inside the *bona fide* spot, we assigned the cell type using the gene expression parameter *a_st_* of the *bona fide* spot that covers the cell, and the nuclear morphological parameters, *μ_t_* and Σ*_t_*, estimated from all *bona fide* spot-covering cells, which is formulated as 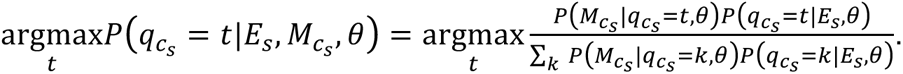

Second, for the cell, which still lies outside the bona fide spot, we kept enlarging the spot size until the missing cells are covered. We also considered both the gene expression parameter *a*_st_ of the enlarged spot that covers the cell, and the nuclear morphological parameters, *μ_t_* and Σ*_t_* to assign the cell type. Of note, we do not update the parameters, even though the spot size is enlarged.

The cell covered by different enlarged spots is assigned based on its highest probability, i.e., for a single cell *c_s_*, and the spot set *S* covering *c_s_*, the cell type is assigned by the following formula:

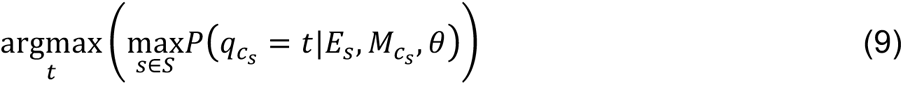

### Cell type signature estimation from spatial transcriptomics and the spatial transcriptomics clustering at the single-cell level

Given the cell type/cluster proportion in each spot, *a_st_*, the signature gene expression *G_it_* was estimated using the following non-negative least square solution:

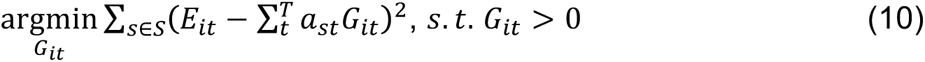

When carrying out clustering with both unknown gene expression signature *G_it_* and unknown cell type proportion of spots, *a*_st_, the initial values of clusters are first given using the spot level clustering, e.g., K-means and Louvain clustering, the cells within the spot are assigned the same initial cluster, and the initial value of cluster signature was set to be the average gene expression of spots belonging to the cluster. In each iteration, the clustering whole transcriptomic signature was re-estimated using Formula (10) in the M-step, and the cluster of each single cell was re-assigned in the E-step.

### High-resolution spatial transcriptomics data simulation from the real data

To take into account different nuclear sizes of cell types, real 10X Visium Spatial transcriptomics data from the mouse brain hippocampus, mouse brain cortex, mouse kidney, and human breast cancer were used to simulate high-resolution spot spatial transcriptomics data. Spot diameters of 30 μm, 20 μm, 10 μm and 5 μm were simulated to cover the whole tissue evenly, where the distance between two spot centers was set to be 2x spot diameter. The spot is defined to cover the cell if the distance of the centroids between the spot and the nucleus is smaller than the sum of the spot radius and the nucleus major radius. The cell is approximately treated as a circle with the cell radius being the nucleus major radius. The cell area covered by the spot was measured as the area of intersection between the spot and the cell, and the spot-covering cell proportion was defined as the area of the intersection divided by the cell area.

Given one high-resolution spot, its transcriptome was simulated by the following formula:

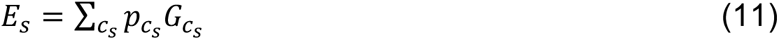

where *c_s_* represents the cell covered by spot *s*, *p_c_s__* represents the cell area proportion covered by spot *s*, and *G_c_s__* represents the whole transcriptomic signature of cell type/cluster *c_s_*.

### Low-resolution spatial transcriptomics data simulation

We simulated the low-resolution ST data to test the STIE and other tools. We simulated 10,000 cells, which are randomly generated from 10 cell types and located in 1,000 spots. We further simulated the gene expression signature comprising of 500 marker genes based on the 10 distinguishable uniform distributions, respectively, representing the average gene expression within each cell type. For each cell type, we generated 100 single cells as the scRNA-seq data (100×10 single cells in total), whose gene expression is simulated from the negative binomial distribution by taking as mean the corresponding gene expression signature. For each spot in the spatial transcriptomics, we simulated its bulk gene expression based on the negative binomial distribution, whose mean is the sum of products between the number of cell types and the corresponding gene expression signatures. Meanwhile, for each cell in the spatial transcriptomics, we simulated 3 nuclear morphological features based on the Gaussian distribution, with the mean uniformly generated between 0 and 3. To investigate the relative contribution of nuclear morphological features and spatial gene expression to the final cell typing at different scenarios, we tweaked the variations of the Gaussian distribution for morphological features and the negative binomial distribution for gene expression, respectively, to generate four combinations between morphological features and spatial gene expression with either high or low noise. To test the nuclear morphological feature selection, we generated 6 more features: 3 intermediate features that are the sum of any two true features along with random noise, and 3 irrelevant features that are completely random noise.

### Spatially resolved cell-cell interactions across the sample tissue

We first used STIE to learn the whole-transcriptomic signature of each cell type/cluster from the spatial transcriptome data. Although the cell type transcriptomic signature pre-defined from scRNA-seq was used for cell type deconvolution, we re-estimated the signature to account for the variation in the real spatial transcriptome data, as well as the whole transcriptome to cover the potential interaction beyond the signature genes. We replicated each cell type transcriptomic signature for times of the corresponding cell count and used it as the single-cell gene expression for CellChat as input so that CellChat can evaluate the interaction significance by permuting cells and recalculating the average gene expression of each permuted cell type.

Calculation of intercellular interaction probability follows the steps in CellChat^39^. Briefly, the ligand-receptor mediated signaling interactions were calculated using the law of mass action. The random walk-based network propagation technique was used to project the single-cell level spatial gene expression onto a protein-protein network from STRINGdb^51^. The communication probability 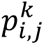 between cell types *i* and *j* for a particular ligand-receptor pair *k* was modeled by CellChat and adapted as follows:

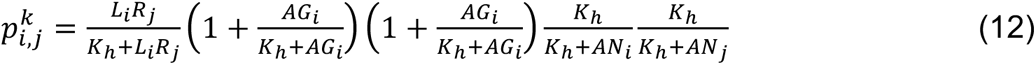

where *L_i_* and *R_j_* represent the expression level of ligand *L* and receptor *R* in cell types, *i* and *j*, respectively, which are approximated by the geometric mean of the expression level of their subunits. *AG* and *AN* represent the average expression of multiple soluble agonists and antagonists for the ligand-receptor pair. The Hill function was used to model the interactions between ligand and receptor as well as the modulations of agonists and antagonists, with a parameter *K_h_* set to be 0.5. The probability of ligand-receptor pairs from the same signaling pathway was summarized, thereby obtaining a communication probability matrix *P_T_*_×*T*×*K*_ between *T*×*T* cell type pairs and *K* signaling pathways.

Of note, CellChat assumes for unsorted single-cell data that abundant cell populations tend to collectively send stronger signals than rare cell populations. However, the spatial cell-cell interaction strength may depend on their distance to a greater extend, compared to their abundance. Accordingly, we did not include the term cell count in cell types *i* and *j* in Equation (12), as CellChat did. Moreover, given a distance cutoff, we recalculate the global cell-cell interaction by considering the spatial proximity.

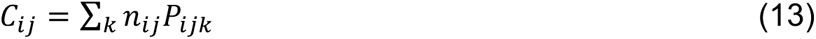

where *n_ij_* represents the number of cell pairs within a distance cutoff, and *C_ij_* represents the weighted total count of spatially resolved interactions between cell types *i* and *j*. We left the distance between nucleus centroids along their major axes as a parameter and set it as default to be 3 μm as the cell-cell attachment distance^52^.

In addition, non-negative matrix factorization (NMF) was applied to the intercellular communication probability matrix to infer the latent outgoing and incoming signal patterns of ligand-receptor pairs or signaling pathways for sender cells and target cells, respectively, which reveal how these ligand-receptor pairs or signaling pathways work together to drive communication for certain sender cell groups and respond to incoming signals for certain target cell groups, respectively. The Cophenetic and Silhouette metrics were used to select the number of latent patterns. Two contribution matrices, 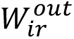 and 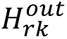, were obtained for outgoing patterns via NMF, representing the contribution of cell group *i* and the contribution of signaling pathway *k* in the incoming pattern *r*, respectively; likewise, two contribution matrices, 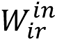 and 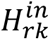, were obtained for the incoming patterns. Furthermore, we calculated the cell communication strength in each outgoing and incoming pattern as follows:

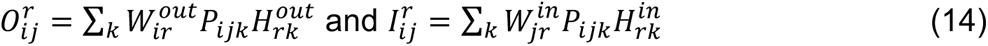

Most of the above functions are implemented and extended based on CellChat.

## Data availability

The authors analyzed the publicly available spatial transcriptome and single-cell RNA-seq datasets. The data were acquired from the following websites or accession numbers: (1) 10X Visium adult mouse brain FFPE (https://www.10xgenomics.com/resources/datasets/adult-mouse-brain-ffpe-1-standard-1-3-0); (2) 10X Visium adult mouse kidney FFPE (https://www.10xgenomics.com/resources/datasets/adult-mouse-kidney-ffpe-1-standard-1-3-0); (3) 10X Visium human breast cancer FFPE (https://www.10xgenomics.com/resources/datasets/human-breast-cancer-ductal-carcinoma-in-situ-invasive-carcinoma-ffpe-1-standard-1-3-0); (4) 10X Visium frozen postmortem human dorsolateral prefrontal cortex (the Globus endpoint ‘jhpce#HumanPilot10x’); (5) 10X CytAssist Spatial Gene Expression (V2 Chemistry) mouse brain coronal section1 FFPE (https://www.10xgenomics.com/resources/datasets/mouse-brain-coronal-section-1-ffpe-2-standard) and section 2 FFPE (https://www.10xgenomics.com/resources/datasets/mouse-brain-coronal-section-2-ffpe-2-standard); (6) Human breast cancer spatial transcriptomics (https://www.spatialresearch.org/resources-published-datasets/doi-10-1126science-aaf2403/); (7) H&E images for primary breast cancer frozen tissue section in HDST (https://singlecell.broadinstitute.org/single_cell/study/SCP420/hdst#study-download); (8) Single nucleus RNA-seq dataset of 1,402 cells in the adult mouse hippocampus (https://singlecell.broadinstitute.org/single_cell/study/SCP1/-single-nucleus-rna-seq-of-cell-diversity-in-the-adult-mouse-hippocampus-snuc-seq#study-download); (9) Single cell RNA-seq dataset of 14,249 adult mouse cortical cell taxonomy from the Allen Institute (https://www.dropbox.com/s/cuowvm4vrf65pvq/allen_cortex.rds?dl=1); (10) Single cell RNA-seq of 26 primary tumors from three major clinical subtypes of human breast cancer (GSE176078). (11) The marker genes of 31 preliminary cell clusters across 7 broad cell types on the single nucleus RNA-seq data from DLPFC using 5,231 nuclei (https://github.com/LieberInstitute/spatialLIBD).

## Code availability

STIE is publicly available as an open-source R package at https://github.com/zhushijia/STIE.

## Acknowledgements

This work was supported by the following grants: National Institute of Health: CA233794, CA255621, CA226052, and R01CA258584/TW; Cancer Prevention and Research Institute of

Texas: RR180016 and RP200554; European Commission: ERC-2014-AdG-671231 and ERC-2020-ADG-101021417.

## Author contributions

This study was conceived by S.Z., Y.H., and G.X. and led by S.Z. S.Z. designed the model and algorithm and implemented the STIE software. S. Z designed the experiments. S.Z. performed the data analysis. N.K. annotated the image. S.Z. and N.K. interpreted the result. S.Z. wrote the manuscript with input from N.K. and T.W. and S.W.

## Notes

### Competing Interest Statement

Advisory board: Helio Health, Espervita Therapeutics
Founding share holder: Alentis Therapeutics
Research funding: Morphic Therapeutics

