## Supplementary Information for "STIE: Single-cell level deconvolution, convolution, and clustering in spatial transcriptomics by aligning spot level transcriptome to nuclear morphology"

**Supplementary Note 1**,

Both gene expression and cell morphology act crucial roles in determining the identities and functions of cells. On the one hand, cell morphology and cell function are closely related. Historically, the cell morphology was one of the first ways to identify the cell types. Some of the direct evidences are the descriptive names of cells, e.g., squamous, stellate, and dendritic cells. Cell morphology serves as a crucial marker of cellular identity across a range of biological contexts, such as, diagnosing cancer by identifying irregular or enlarged nuclear morphology, determining differentiation status of stem cells, and elucidating neural circuits through axonal and dendritic structures. On the other hand, a cell typically expresses only a fraction of its genes, and the different types of cells in multicellular organisms arise because different sets of genes are expressed, making the gene expression a more direct indicator of cell types. More recently, single-cell transcriptome profiling has enabled unbiased definition of cell types using transcriptomic features. Ideally, the cell typing based on cell morphology and gene expression are the same, and therefore, STIE models jointly the morphological information and spatial gene expression in the spatial transcriptomics, aiming to find their consensus on the cell typing. Of note, there also exist the scenarios, where the cell types are transcriptomically distinct but morphologically similar, such as the nuanced immune cells. Compared with the current low-resolution spot spatial transcriptomics, STIE working on the high-resolution spot could provide the superior power in distinguishing the nuanced cell types (Fig. 4j-k).

The definition of cell type proportions can be different between low- and high-resolution spots. In the low-resolution spot, the cell type proportion refers to the cell count of each cell type divided by total cell count covered by the spot, while in the high-resolution spot, the cell type proportion refers to the cell area of each cell type divided by total cell area covered by the spot. For example, 5 cell type 1 (C1), 3 cell type 2 (C2) and 2 cell type 3 (C3) in one low-resolution spot give the proportion of 50% C1, 30% C2 and 20% C3, while 0.5 C1, 0.3 C2 and 0.2 C3 in the high-resolution spot give the same proportion of 50% C1, 30% C2 and 20% C3. Both proportions can be estimated by STIE in the similar way.

**Supplementary Note 2**, Compared to the fresh frozen tissue slice, the FFPE preserves and stabilizes the fragile structure inside and between the cells in the tissue, making the FFPE better for investigating the tissue and cell morphology. In addition, the tissue slice made from the fresh frozen sample is often thicker (>=10 μm) than the ST-FFPE H&E slide (5 μm). The density of nuclei (or cells) is high and the nuclear contour is blurred, both of which may further deteriorate the nuclear/cellular segmentation.

*scRNA-seq derived cell type transcriptomic signatures:*

- Mouse brain hippocampus: The cell type transcriptomic signature (the file "Major_cell_types_marker_genes.txt") was downloaded from the link (https://singlecell.broadinstitute.org/single_cell/study/SCP1/-single-nucleus-rna-seq-of-cell-diversity-in-the-adult-mouse-hippocampus-snuc-seq#study-download). It is the average gene expression for 773 marker genes from the major cell types (CA1, CA2, CA3, DG, and Glia) of the mouse hippocampus, which is derived from the single nucleus RNA-seq dataset comprising 25,392 genes and 1,402 cells in the adult mouse hippocampus. After intersection with the 10X Visium mouse brain FFPE spatial transcriptomics, the cell-type signature remains 670 marker genes.
- Mouse brain cortex: The scRNA-seq dataset comprising 34,617 genes and 14,249 adult mouse cortical cell taxonomy from the Allen Institute (the file "allen_cortex.rds") was downloaded from the link (https://www.dropbox.com/s/cuowvm4vrf65pvq/allen_cortex.rds?dl=1). The buildSignatureMatrixMAST() function in the DWLS R package was used to build cell type transcriptomic signature, with variable "subclass" from the "meta.data" table being the cell type. The default parameters of an FDR adjusted p-value of <0.01 (defined using the hurdle model in the MAST R package), and a log2 mean fold change >0.5 were used to determine the marker genes.
- Human breast cancer: The scRNA-seq dataset comprising 29,733 genes and 100,064 cells of 26 primary tumors from three major clinical subtypes of human breast cancer was downloaded from GEO (GSE176078). It comprises four files, count_matrix_sparse.mtx (raw count matrix), count_matrix_barcodes.tsv (cell barcode), count_matrix_genes.tsv (gene annotation) and metadata.csv (cell type labelling). The 100,064 cells were down-sampled to keep 15% cells for each cell type. The buildSignatureMatrixMAST() function in DWLS R package was used to build cell type transcriptomic signature, with the "celltype_major" column in the "metadata.csv" file being cell type. The default parameters of an FDR adjusted p-value of <0.01 (defined using the hurdle model in the MAST R package), and a log2 mean fold change >0.5 were used to determine the marker genes.

*The other methods used in the paper:*

- K-means: The K-means clustering on 10X Visium ST data was implemented in spaceranger ver. 1.3.0. The base R *stats::kmeans()* function was used for the simulated high-resolution spot spatial transcriptomics data.
- SPOTlight (*ver 0.1.7*): The *Seurat::SCTransform()* was first used to normalize the scRNA-seq data, followed by the *Seurat::FindAllMarkers()* function to find cell type gene markers. Further, the *SPOTlight::spotlight_deconvolution()* function was used to deconvolute the spot level gene expression via "nsNMF" factorization method.
- DWLS: The *buildSignatureMatrixMAST()* function was used to build cell type transcriptomic signatures from the scRNA-seq data with default parameters; the *solveDampenedWLS()* function was used to deconvolute both spot level gene expression and clustering derived CAGE.
- Stereoscope (*ver 0.2.0*): The python package, *stereoscope*, was used to deconvolute the spot level gene expression, with parameters, *--sc_cnt*, *--sc_labels*, and *--st_cnt* pointing to scRNA-seq count matrix, cell type label matrix and spatial transcriptomics count matrix, respectively.
- RCTD (*spacexr ver 2.0.0*): The *spacexr::Reference()* and *spacexr::SpatialRNA()* functions were run on scRNA-seq and spatial transcriptomics data, respectively, to build the corresponding objects. The *spacexr::create.RCTD()* function was used to build the RCTD object and *spacexr::run.RCTD()* was used to deconvolute the spot level gene expression.
- Tangram (*ver 1.0.3*): The spatial data and single cell data were pre-processed using the pp_adatas() function. The map_cells_to_space () function was used to map the single cells onto the Visium spots. The returned matrix is a cell-by-spot structure, where each item [*i, j*] gives the probability for cell *i* to be in spot *j*. To compare with the other methods, in each spot, we sum up the probabilities of single cells of the same cell type and further normalize the probabilities of cell types to their sum. The normalized probabilities are treated as the cell type proportions in the spot.
- BayesPrim (*ver 2.0*): The runBayesPrism() function of the R package, *BayesPrim*, was used to deconvolute the spot level gene expression. The parameter of cell states was set to the same with cell types. The parameter of species was set to be "mm" and "hs" for mouse and human samples, respectively.
- BayesSpace (*ver 1.4.1*): The *BayesSpace::spatialCluster()* function was used to perform clustering on spot level gene expression. The *BayesSpace::enhanceFeatures()* function

was used to impute the gene expression from spot level to subspot level with default parameters.

- xfuse (ver 0.2.1): The command “xfuse run” along with parameters of “metagenes” and “gene_maps” in the config file was used to impute the gene expression in the micrometer resolution by following the tutorial (https://github.com/ludvb/xfuse).
- SCINA (*ver 1.2.0*): The *preprocessCore::normalize.quantiles()* function was first used to normalize the scRNA-seq read count. The *SCINA::SCINA()* function was used to assign cell types to the scRNA-seq data, with parameter settings: “*rm_overlap=TRUE*” and “*allow_unknown=TRUE*” to remove overlapping signatures and assign “unknown” to the cells with unclear identities, respectively.
- SpaGCN (*ver 1.2.5*): The SpaGCN was installed under python (version=3.8.8), with versions of the dependent packages installed by following the tutorial. The spatial gene expression, spatial location, and histology image were used as input. The easy mode detect_spatial_domains_ez_mode() function was run to perform the clustering, where the parameter “shape” was set to be "hexagon" for Visium data. Following up, the function spatial_domains_refinement_ez_mode() was run to refine the clustering.
- MUSE: For the morphology, we identified corresponding H&E regions for each spot using ST positions, resized image tiles to 299 × 299 pixels and learned deep embeddings using Inception v3. For the transcript, we used median normalization and the log1p transform and selected the top 500 most variable genes. From the morphology and transcriptome modalities, the *muse.muse_fit_predict()* function was used to learn the MUSE features, which were further used to perform clustering by the *phenograph.cluster()* function.

**STIE model:**

We first listed the variables used in the model:

- $N$ , $C$, $T$ , $S$, and $F$ represent the number of genes, cells, cell types, spots, and morphological features, respectively.
- $G = \{G_{it}\}_{1 \le i \le N; 1 \le t \le T}$ is a matrix of cell-type gene expression signatures, where $G_{it}$ is the expression of the $i$-*th* signature gene of the $t$-*th* cell type.
- $E = \{E_{is}\}_{1 \le i \le N; 1 \le s \le S}$ is a matrix of gene expression from the ST, where $E_{is}$ is the expression of the $i$-*th* gene in the $s$-th spot.
- $c = \{c_s\}_{1 \le s \le S}$ is the cell index on the ST histology image, where $c_s$ represents the $c$-th cell in *the* $s$-*th* spot, and $C_s$ represents the total cell count in *the* $s$-*th* spot. The parameter $\gamma$ is used to represent the *bona fide* area captured by the spot, which can be different from the reported spot size.
- $M = \{M_{cf}\}_{1 \le c \le C; 1 \le f \le F}$ is a matrix of nuclear morphological features obtained from the ST histology image, where $M_{cf}$ is the value of the $f$-*th* morphological feature of the $c$-th cell.
- Let $q = \{q_c\}_{1 \le c \le C}$ represent the hidden cell type of one single cell that generates the observed morphological features $M_c$ and gene expression $E_c$, with $q_c = t$ representing the *c-th* cell that takes the cell type $t$.

We formulate that $E_s = \sum_t^T a_{st} G_t + \varepsilon_s,\ s.t.\ a_{st} > 0\ and\ \varepsilon_s \sim N(0, \sigma^2)$, where $a_{st}$ is the non-negative regression coefficient for the $s$-th spot and the $t$-th cell type. On the other hand, we assume that the probability of the nuclear morphological feature follows a Gaussian distribution, $P\left(M_{c_s} \mid q_{c_s} = t\right) \sim N(\mu_t, \Sigma_t)$ , where $\mu_t$ and $\Sigma_t$ are the mean and variance of the nuclear morphological features of the $t$-*th* cell type, respectively. Thus, the parameter of STIE is a 3-tuple: $\theta = \{a_{st}, \mu_t, \Sigma_t\}$.

The observed data pdf given $\theta$ can be formulated as:

$$P(M, E|\theta) = \sum_q P(M, E|q, \theta) P(q|\theta) \qquad (1)$$

We used the Expectation-Maximization (EM) algorithm to solve the STIE model and estimate $\theta$. The Q function takes the following form:

$$Q\left(\theta|\theta'\right) = \sum_q P\left(q|M, E, \theta'\right) logP(M, E, q|\theta) \qquad (2)$$

By expanding all cells, the Q function is rewritten as:

$$\sum_{q_1} \sum_{q_2} \cdots \sum_{q_C} P(q_1, q_2, \dots, q_C | M_1, M_2, \dots, M_C, E_1, E_2, \dots, E_C, \theta') logP(M_1, M_2, \dots, M_C, E_1, E_2, \dots, E_C, q_1, q_2, \dots, q_C | \theta)$$

(3)

We assume that each cell is generated independently, so, we obtained that

$$=\sum_{q_1}\sum_{q_2}\dots\sum_{q_C} P(q_1,q_2,\dots,q_C|M_1,M_2,\dots,M_C,E_1,E_2,\dots,E_C,\theta')\{logP(M_1,E_1,q_1|\theta) + logP(M_2,E_2,q_2|\theta) + \cdots + logP(M_C,E_C,q_C|\theta)\} \quad (4)$$

$$=\sum_{q_1} logP(M_1,E_1,q_1|\theta)\sum_{q_2}\dots\sum_{q_C} P(q_1,q_2,\dots,q_C|M_1,M_2,\dots,M_C,E_1,E_2,\dots,E_C,\theta') + \sum_{q_2} logP(M_2,E_2,q_2|\theta)\sum_{q_1}\sum_{q_3}\dots\sum_{q_C} P(q_1,q_2,\dots,q_C|M_1,M_2,\dots,M_C,E_1,E_2,\dots,E_C,\theta') + \cdots + \sum_{q_C} logP(M_C,E_C,q_C|\theta)\sum_{q_1}\dots\sum_{q_{C-1}} P(q_1,q_2,\dots,q_C|M_1,M_2,\dots,M_C,E_1,E_2,\dots,E_C,\theta') \quad (5)$$

$$=\sum_{q_1} P(q_1|M_1,E_1,\theta')logP(M_1,E_1,q_1|\theta) + \sum_{q_2} P(q_2|M_2,E_2,\theta')logP(M_2,E_2,q_2|\theta) + \cdots + \sum_{q_C} P(q_C|M_C,E_C,\theta')logP(M_C,E_C,q_C|\theta) \quad (6)$$

The Q function is rewritten as:

$$\sum_{s\in S}\sum_{c_s}\sum_t P\left(q_{c_s}=t|M_{c_s},E_{c_s},\theta'\right)logP\left(M_{c_s},E_{c_s},q_{c_s}=t|\theta\right) \quad (7)$$

We assume that $E$ and $M$ are conditionally independent given the cell type, i.e., the value of $q$. Therefore,

$$P(M,E,q|\theta) = P(M|q,\theta)P(E|q,\theta)P(q|\theta) = P(M|q,\theta)P(q|E,\theta)P(E|\theta) \quad (8)$$

Thus, the Q function becomes

$$\sum_{s\in S}\sum_{c_s}\sum_t P\left(q_{c_s}=t|M_{c_s},E_{c_s},\theta'\right)logP\left(M_{c_s}|q_{c_s}=t,\theta\right) + \sum_{s\in S}\sum_{c_s}\sum_t P\left(q_{c_s}=t|M_{c_s},E_{c_s},\theta'\right) logP\left(q_{c_s}=t|E_{c_s},\theta\right) + \sum_{s\in S}\sum_{c_s}\sum_t P\left(q_{c_s}=t|M_{c_s},E_{c_s},\theta'\right) logP\left(E_{c_s}|\theta\right) \quad (9)$$

The transcriptome of cells in the same spot are profiled in bulk and observed as the same gene expression profile, i.e., $E_{c_s}=E_s$. Likewise, $P\left(q_{i_s}=t|E_{i_s},\theta\right)$ of the cell $i_s$ and $P\left(q_{j_s}=t|E_{i_s},\theta\right)$ of the cell $j_s$ from the same spot $s$ are also indistinguishable, and therefore, we obtain that for any cell $c_s$ in the spot $s$, $P\left(q_{c_s}=t|E_{c_s},\theta\right) = P\left(q_{c_s}=t|E_s,\theta\right) = a_{st}/\sum_t a_{st}$. So, the Q function becomes:

$$\sum_{s\in S}\sum_{c_s}\sum_t P\left(q_{c_s}=t|M_{c_s},E_s,\theta'\right)logP\left(M_{c_s}|q_{c_s}=t,\theta\right) + \sum_{s\in S}\sum_t log\frac{a_{st}}{\sum_t a_{st}}\sum_{c_s} P\left(q_{c_s}=t|M_{c_s},E_s,\theta'\right) + \sum_{s\in S} logP(E_s|\theta)\sum_{c_s}\sum_t P\left(q_{c_s}=t|M_{c_s},E_s,\theta'\right) \quad (10)$$

$$=\sum_{s\in S}\sum_{c_s}\sum_t N_{c_st}logP\left(M_{c_s}|q_{c_s}=t,\theta\right) + \sum_{s\in S}\left\{\sum_t N_{st}\log\frac{a_{st}}{\sum_t a_{st}} + N_s logP(E_s|\theta)\right\} \quad (11)$$

where $N_{c_st} = P\left(q_{c_s}=t|M_{c_s},E_s,\theta'\right)$, $N_{st}=\sum_{c_s} N_{c_st}$, and $N_s = \sum_{c_s}\sum_t N_{c_st}$

Next, we demonstrate how to calculate $N_{c_st}$. In one spot $s$, given the bulk gene expression $E_s$ in the spot and its morphological features $M_{c_s}$, one single cell $c_s$ is of cell type $t$, taking the following probability:

$$N_{c_s t} = P\left(q_{c_s} = t \middle| E_s, M_{c_s}, \theta'\right) \tag{12}$$

$$= \frac{P\left(E_s, M_{c_s} \middle| q_{c_s} = t, \theta'\right) P\left(q_{c_s} = t \middle| \theta'\right)}{\sum_t \ P\left(E_s, M_{c_s} \middle| q_{c_s} = t, \theta'\right) P\left(q_{c_s} = t \middle| \theta'\right)} \tag{13}$$

$$= \frac{P\left(M_{c_s} \middle| q_{c_s} = t, \theta'\right) P\left(E_s \middle| q_{c_s} = t, \theta'\right) P\left(q_{c_s} = t \middle| \theta'\right)}{\sum_t \ P\left(M_{c_s} \middle| q_{c_s} = t, \theta'\right) P\left(E_s \middle| q_{c_s} = t, \theta'\right) P\left(q_{c_s} = t \middle| \theta'\right)} \tag{14}$$

$$= \frac{P\left(M_{c_s} \middle| q_{c_s} = t, \theta'\right) P\left(E_s, q_{c_s} = t \middle| \theta'\right) / P\left(E_s \middle| \theta'\right)}{\sum_t \ P\left(M_{c_s} \middle| q_{c_s} = t, \theta'\right) P\left(E_s, q_{c_s} = t \middle| \theta'\right) / P\left(E_s \middle| \theta'\right)} \tag{15}$$

$$= \frac{P\left(M_{c_s} \middle| q_{c_s} = t, \theta'\right) P\left(q_{c_s} = t \middle| E_s, \theta'\right)}{\sum_t \ P\left(M_{c_s} \middle| q_{c_s} = t, \theta'\right) P\left(q_{c_s} = t \middle| E_s, \theta'\right)} \tag{16}$$

In the M-step, we first take the derivative of the Q function with respect to $\mu_t$ and $\Sigma_t$, respectively. Of note, these two variables are only in the first item, i.e., the derivative of the second term with respect to these two variables is zero.

The pdf of the Gaussian normal of morphological features is that

$$P\left(M_{c_s}, q_{c_s} = \mathrm{t} \middle| \theta\right) = \frac{1}{2\pi^{F/2} |\Sigma_t|^{1/2}} \exp\left(-\frac{1}{2}\left(M_{c_s} - \mu_t\right)^T \Sigma_t^{-1} \left(M_{c_s} - \mu_t\right)\right) \tag{17}$$

Thus, the first item of the Q function becomes:

$$\sum_{s\in S} \sum_{c_s} \sum_t \ N_{c_s t} \left(-\frac{F}{2} log 2\pi - \frac{1}{2} log|\Sigma_t| - \frac{1}{2}\left(M_{c_s} - \mu_t\right)^T \Sigma_t^{-1} \left(M_{c_s} - \mu_t\right)\right) \tag{18}$$

To optimize the first item, we obtained the following equations. The derivation is very similar to that of the Gaussian Mixture Model, so, we omit the detail.

$$\mu_{tf} = \frac{1}{N_t} \sum_{s\in S} \sum_{c_s} N_{c_s t} M_{c_s f} \tag{19}$$

$$\Sigma_t = \frac{1}{N_t} \sum_{s\in S} \sum_{c_s} N_{c_s t} (M_{c_s} - \mu_t)(M_{c_s} - \mu_t)^T \tag{20}$$

where $N_t = \sum_{s\in S} \sum_{c_s} N_{c_s t}$

Like the parameter of morphological features that are only in the first item, the parameter $a_{st}$ for spatial gene expression are only in the second term. So, we only need to optimize the second term to solve out $a_{st}$. Each spot can be solved independently by maximizing the following:

$$\propto -\frac{1}{2\sigma^2} \sum_i \ \left(E_{is} - \sum_t \ a_{st} G_{it}\right)^2 + \sum_t \ \frac{N_{st}}{N_s} \log \frac{a_{st}}{\sum_t a_{st}} \tag{21}$$

This is not a convex optimization. But according to the Gibbs' inequality, $\sum_i \ p_i log p_i \geq \sum_i \ p_i log q_i$, the right part reaches the maximum at $\frac{a_{st}}{\sum_t a_{st}} = \frac{N_{st}}{N_s}$. The term $\frac{a_{st}}{\sum_t a_{st}}$ represents the cell type proportion estimated from gene expression (we denoted it as $Prop_t(E_s|\theta')$), while $\frac{N_{st}}{N_s}$ represents the cell type proportion estimated from both gene expression and nuclear morphology (denoted by $Prop_t(M_s, E_s|\theta')$). Further, we assume that $Prop_t(E_s|\theta') = Prop_t(M_s|\theta') = Prop_t(M_s, E_s|\theta')$, that is, the proportions of cell type estimated from gene expression and that from nuclear morphology are the same (Supplementary Note 1), and our aim is to find their consensus $Prop_t(M_s, E_s|\theta')$ as the final estimation of cell type proportions. Accordingly, we also searched a local solution for the left part of Formula (21) in the above small area, i.e., $\|Prop_t(E_s|\theta) - Prop_t(M_s|\theta')\|_2^2 \leq c \; and \; c \geq 0$. We reformulated the Formula (21) as a constrained minimization problem, which takes an approximate solution of the left part via the non-negative least square along with the local area derived from the right part as a constraint to restrict their solution:

$$\underset{a_{st}}{\operatorname{argmin}} \frac{1}{2\sigma^2}(E_s - \textstyle\sum_t^T a_{st}G_t)^2 \tag{22}$$

$$\textstyle\sum_t \left(Prop_t(E_s|\theta) - Prop_t(M_s|\theta')\right)^2 \leq c,\ c \geq 0 \text{ and } a_{st} \geq 0 \tag{23}$$

where $Prop_t(E_s|\theta) = P\left(q_{c_s} = t|E_s, \theta\right) = a_{st}/\sum_t a_{st}$ and $Prop_t(M_s|\theta') = \sum_{c_s} P\left(q_{c_s} = t|M_{c_s}, \theta'\right)/\sum_{c_s}\sum_t P\left(q_{c_s} = t|M_{c_s}, \theta'\right) = \sum_{c_s} P\left(q_{c_s} = t|M_{c_s}, \theta'\right)/C_s$ represent the proportions of cell type $t$ in spot $s$, which are estimated from gene expression and nuclear morphology, respectively. The constraint also serves as penalty to balance the gene expression and nuclear morphology more directly.

Since $2\sigma^2$ is a constant, we ignore it in the above minimization. Further, similar to Tikhonov regularization, we use a Lagrange multiplier and rewrite the problem as following:

$$\underset{a_{st}}{\operatorname{argmin}} \sum_i \left(\sum_t a_{st}G_{it} - E_{is}\right)^2 + \lambda \sum_t \left(\frac{a_{st}}{\sum_t^T a_{st}} - \frac{\sum_{c_s} P(q_{c_s}=t|M_{c_s},\theta')}{C_s}\right)^2, \tag{24}$$

Theoretically, there is a one-to-one correspondence between $c$ and $\lambda$. Thus, we transformed Formula (21) into an easier quadratic optimization and simultaneously optimized the left and right parts of Formula (21) around $Prop_t(E_s|\theta') = Prop_t(M_s|\theta')$.

By assuming that $\sum_t^T a_{st}$ approximately equals the current estimate $\sum_t^T a'_{st}$, we obtained

$$\sum_i \left(\sum_t a_{st}G_{it} - E_{is}\right)^2 + \zeta \sum_t \left(a_{st} - \sum_t a'_{st} \frac{\sum_{c_s} P(q_{c_s}=t|M_{c_s},\theta')}{C_s}\right)^2, \text{ where } \zeta = \lambda/\left(\sum_t a'_{st}\right)^2$$

The formula can be represented in the format of inner products:

$$(Ga_s - E_s)^T(Ga_s - E_s) + \zeta\left(a_s - a'_s p_s\right)^T\left(a_s - a'_s p_s\right) \tag{25}$$

where $a_s = (a_{s1}, \ldots, a_{sT})^T$, representing the $T\times1$ vector of regression coefficients for the $s$-th spot; $G = (G_{it})$, representing the matrix of signature gene expression whose first index refers to the marker gene and the second index refers to the cell type; $a'_s = \sum_t a'_{st}$, representing the sum of the current estimates of $a_s$; and $p_s = (p_{s1}, \ldots, p_{sT})^T$, where $p_{st} = \frac{\sum_{c_s} P(q_{c_s}=t|M_{c_s},\theta')}{C_s}$.

After expanding, we obtained that

= $a_s^T G^T G a_s \; - 2E_s^T G a_s \; + E_s^T E_s \; + \zeta a_s^T I a_s$ -$2\zeta p_s^T a_s' I a_s$ +$\zeta p_s^T a_s' a_s' p_s$ (26)

= $a_s^T (G^T G + \zeta I) a_s \; - 2(E_s^T G + \zeta p_s^T a_s' I) a_s \; + E_s^T E_s \; + \zeta p_s^T a_s' a_s' p_s$ (27)

Since $E_s^T E_s$ and $\zeta p_s^T a_s' a_s' p_s \; = \lambda p_s^T p_s$ are constants, we obtained

∝ $a_s^T (G^T G + \zeta I) a_s \; - 2(E_s^T G + \zeta p_s^T a_s' I) a_s$ (28)

To match the standard form of quadratic linear programming:

$$\underset{a_s}{\operatorname{argmin}}\, a_s^T D a_s - 2d^T a_s \,, \;\; s.t. \; A^T a_s \geq b \tag{29}$$

We got that

$$D = G^T G + \zeta I \tag{30}$$

$$d = G^T E_s \; + \zeta a_s' I p_s \tag{31}$$

$$A = I \tag{32}$$

$$b = 0 \tag{33}$$

Given the standard form, we solved the quadratic linear programming using the R package "quadprog" (https://cran.r-project.org/web/packages/quadprog/index.html). The function is solve.QP(D,d,A,b).

As shown above, to maximize the whole Q-function in the M-step, the estimation formula of parameters are derived separately for the morphological feature and spatial gene expression, since they are only in the first and second items, respectively. However, in the iterations of EM algorithm, they are not updated independently. In each E-step, we use the parameters of morphological features estimated from the last step to update the parameter of gene expression, and vice versa, therefore, guaranteeing the convergence of the algorithm.

***Selection of hyperparameters and morphological features:***

**(1) Selection of γ:** The hyperparameter γ aims to model the actual area covered by the spot, which may differ from the reported spot size. The difference might result from the sample leaking from the spot area to the surroundings. We select γ as the area giving the best concordance between the prediction and true spatial gene expression, which is measured by the RMSE. We evaluated γ on the 10X Visium mouse brain and 10X Visium human breast cancer FFPE spatial transcriptomics datasets (Fig. 5a-b), and observed the similar patterns across all different λ, all of which reach the minimum around γ=2.5x. Moreover, we evaluated γ on the new 10X Visium V2 Chemistry CytAssist datasets from two mouse brain replicates (Fig. 5c) and consistently observed the similar patterns, with the minimum RMSE around γ=2x~2.5x. These facts suggest that the selection of γ is independent of λ and stable in different 10X Visium datasets.

**(2) Selection of λ:** The hyperparameter λ balances the information from gene expression and histological features. Given the morphological features, we used a heuristic strategy to select λ via evaluating the balance between two criteria: RMSE of the spatial gene expression fitting and log-likelihood of the morphological feature fitting (Supplementary Fig. 3).

We used the simulation to benchmark the selection of λ. We randomly generated the spatial gene expression data for 1,000 spots based on the negative binomial distribution, 10,000 cells that randomly locate on those 1,000 spots, and 3 nuclear morphological features for each cell based on the Gaussian distribution. To investigate the relative contribution to the cell typing at different scenarios, we generated four combinations between morphological features and spatial gene expression with either high or low noise. Overall, with the elevated λ for morphology, the weight of gene expression decreases in the model, making the gene expression less fitting, and therefore, the RMSE is elevated (Supplementary Fig. 3a-d, second panels). Meanwhile, the increasing weight of morphology enhances the fitting of morphological features and therefore, increases the corresponding morphology log-likelihood (Supplementary Fig. 3a-d, third panels). Specially, in the first scenario (Supplementary Fig. 3a), where both gene expression and morphological features are good, both RMSE and morphology log-likelihood increase stably, but at λ=1e6, the fitting of morphology increases relatively more than the RMSE, suggesting the sacrifice of RMSE gain much more morphological fitting. So, we chose λ=1e6 for this dataset. Consistently, the correlation between prediction and ground truth increases with inclusion of morphology until λ=1e6, while after λ=1e6, more inclusion of morphology decreases the correlation. Similarly, in the second (Supplementary Fig. 3b) and fourth (Supplementary Fig. 3d) scenarios, the morphological fitting increases relatively more than the RMSE at both λ=1e6, so, we chose λ=1e6 for both those scenarios. It suggests that when gene expression is poor or morphology is good, the model fitting can benefit from the inclusion of morphology. According to the above criterion, we chose λ=1e5 for the third scenario (Supplementary Fig. 3c), since it also gains more morphological fitting with a small sacrifice in the gene expression fitting. Of note, the gene expression is good and much better than morphology in the third scenario, and more inclusion of morphology does not significantly contribute to the total model fitting, i.e., the best setting, λ=1e5 gives very similar performance with that under λ=1e0.

**(3) Selection of nuclear morphological features:** The strategy is that given λ, we rank the nuclear morphological features based on the RMSE by running STIE on each feature individually. Next, based on their ranking, we used a greedy strategy to gradually add more features to the model. Like the selection of λ, the best morphological features are selected by evaluating the gene expression and morphological fittings simultaneously. We investigated the morphological features over different λ.

To benchmark the nuclear morphological feature selection, we performed the simulation analysis based on the simulation scenarios in (2). 6 more features were generated based on the 3 true features, including 3 intermediate features that are the summation of any two true features along with the random noise, and 3 irrelevant features that are completely random noise (Supplementary Fig. 4). The observations were made as follows: First, given relatively small values of λ, all true features were correctly ranked in the front, followed by the intermediate features, and the irrelevant features go to the very last. Second, given the high values of λ, the model falsely ranked the features. However, we can simply eliminate them due to their high RMSE and low morphological likelihood. Last, given three true features, all scenarios reach the minimum RMSE of gene expression fitting and maximum likelihood of morphology. Thus, we find the correct feature combination by simultaneously considering the gene expression and morphology fitting.

**(4) Selection of nuclear morphological features and λ via grid search in the real datasets:** In the real datasets, the selection is a bit simpler, as we only focused on two categories of morphological features, shape and size. As shown above, we selected the morphological features and λ using the similar strategy by considering the fitting of gene expression and nuclear morphology, and therefore, we used that strategy to select the morphological feature and λ simultaneously via the grid search.

In the real datasets, we extracted multiple morphological features for the nuclei, and observed that these features are highly correlated (Supplementary Fig. 7), which is consistent with their definition and mathematical calculation. These features can be grouped into three categories: size (Area, Major, Minor, Width, Height, Feret, and Perimeter), shape (Round and Circular) and angle (FeretAngle and Angle). To reduce the redundancy and improve the efficiency, we performed PCA for each category and took the 1st PC as the surrogate of each category. Further, we assume that the "angle" of nuclei does not contribute to the cell typing. Also, in case of overfitting to the irrelevant feature, we exclude 'angle', and only used two aggregated features, i.e., 'size' and 'shape'.

In the mouse brain hippocampus deconvolution and clustering (Supplementary Fig. 8a-f), the feature 'shape' was ranked before 'size', and 'shape' gives the lower RMSE and higher morphological likelihood at small $\lambda$, so we used 'shape' as the morphological feature and $\lambda=0$ (red triangle). In the mouse/human brain cortex deconvolution and clustering (Supplementary Fig. 9a-d), the feature 'shape' was ranked before 'size'. The RMSE at 'shape' is not the minimum. However, the RMSE does not change significantly at 'shape+size', while the morphological likelihood dropped more, so, we chose 'shape' as morphological feature and $\lambda=0$ due to higher morphological likelihood. In the human breast cancer clustering (Supplementary Fig. 10b), both RMSE and morphological likelihood were reduced when increasing features. We chose "shape+size" as morphological features and $\lambda=1e4$ (orange rectangle), to confer more reduction in RMSE with sacrificing morphological fitting. To be consistent with clustering, we also chose the same setting for the deconvolution (Supplementary Fig. 10a), which also gives the lowest RMSE.

Of note, it is well-known that tuning hyperparameters often rely on both good heuristics and domain knowledge. Although we provide a heuristic strategy above, we highly recommend borrowing the domain knowledge from the biologist/pathologist. For example, the hyperparameter could be tuned via matching the cell typing of a small subset of cells manually annotated.

We implemented an R function STIE_search(), which performed the above strategy of the grid search across hyperparameter $\lambda$ and nuclear morphological feature combinations.

**Human dorsolateral prefrontal cortex data processing**

The processed gene-level read count matrix, histology image, and nuclear segmentation of 10X Visium frozen postmortem human dorsolateral prefrontal cortex were downloaded from http://research.libd.org/globus. The nuclear morphological features were extracted using ImageJ. We focused on the layer-enriched genes prioritized by the 'enrichment' model[1] (7 models per gene, 1 per layer), which tested the differences in expression between one layer versus all other layers. We further ranked the layer-enriched genes based on their FDR by the enrichment model and selected the top 100 genes for each layer. The union of layer-enriched genes of seven layers (670 in total) were used in the clustering.

**STIE clustering on the frozen human brain cortex and FFPE mouse brain cortex spatial transcriptomics**

The frozen section may not preserve well the morphology compared to the FFPE, and therefore, we compared the STIE performance between the frozen section and FFPE to evaluate the impact of accurate nuclear morphology on the STIE performance. First, we compared the quality of the histology image between fresh frozen postmortem human brain cortex (Supplementary Fig. 30a) and FFPE mouse brain cortex (Supplementary Fig. 30b). As expected, FFPE is much better than the frozen section. The background of frozen section is highly noisy with small dots all over the image, and the nuclear shape is blurred with less clear boundary. Thus, even though the nuclear positions can be identified, the morphological information cannot be determined accurately. For example, the cytoplasm is misclassified as nuclear (Supplementary Fig. 30a), making the morphological information falsely calculated, e.g., size and shape.

We run STIE clustering at single-cell level on the 10X Visium frozen human brain cortex spatial transcriptomics dataset, by taking 'shape' as the morphological feature and $\lambda=0$ (Supplementary Fig. 9). The layers (Supplementary Fig. 28a) are clearly observed, which are partially concordant with the manual annotation (Supplementary Fig. 28b) with cells saturating to the surrounding layers. Meanwhile, we run K-means clustering at spot-level on the gene expression (Supplementary Fig. 28c). To make comparison, we used a simple way to aggregate the STIE-derived single cells to the spot (Supplementary Fig. 28d): we first calculated the proportions of STIE clusters in each spot, and then run K-means on the proportions to obtain the spot-level clusters. This could be further improved by considering neighborhood information, but beyond the scope of this paper. Although the morphological information is not accurate, STIE demonstrates higher concordance with the manual annotation than K-means (ARI=0.30 for STIE; ARI=0.22 for K-means). On the other hand, we run STIE clustering on the 10X Visium mouse brain cortex

FFPE spatial transcriptomics, which has better image quality and nuclear segmentation. We chose ‘shape’ as morphological feature and λ=0 (Supplementary Fig. 9). Similarly, STIE on the mouse brain cortex FFPE also outperformed K-means, with the cortex layers more clearly defined. STIE starts with the spot-level K-means clustering as initial values, but STIE outperformed K-means in both mouse brain cortex FFPE and human brain cortex frozen section, suggesting that STIE further refined the K-means clustering by incorporating the morphological features as additional information.

Of note, the human brain frozen section and the mouse brain FFPE are not directly comparable due to different species and underlying biological mechanisms. Therefore, we compared these two data with themselves under different parameter settings. We assume that the good extraction of nuclear morphology may contribute to the deconvolution similarly, with different morphological features changing the result to a relatively small extent. Currently, the best morphological features are both “shape” for the above two data. We further add more features to the STIE, i.e., “shape+size”, and check whether the results are stable or not. Consequently, in the mouse brain cortex FFPE (Supplementary Fig. 29d), the STIE result using “shape+size” is still largely consistent (76% concordance by average, Supplementary Fig. 29e) with that using only “shape”, suggesting “size” also has consistent distinguishable capability. As opposed, in the human brain cortex FF (Supplementary Fig. 29a-b), the STIE result using “shape+size” becomes much less concordant (52% concordance, Supplementary Fig. 29c) with that using “shape”. This is largely because the “size” is not well extracted, further misleading the classification. We also downgraded the STIE single-cell level clustering to the spot-level and compared with the manual annotation (ARI=0.26). Put together, STIE on FFPE demonstrates better and more stable performance than STIE on the frozen section, because FFPE better preserves the histology and provides more accurate nuclear segmentation.

The above analysis only aims to compare the performance of STIE on the FFPE with that on the fresh frozen sections. Further, we found that the performance of clustering could be improved by more properly setting the initial values, which is consistent with the common knowledge of the traditional clustering methods. Instead of the spot-level K-means clustering as the initial value, we used the spot-level SpaGCN clustering and follow up with the single-cell level clustering by STIE. To compare with the spot-level manual annotation, we further downgraded the single-cell level (Supplementary Fig. 29f) to the spot level (Supplementary Fig. 29g). The ARI is 0.5, which is comparable to the other spot-level clustering methods.

**Supplementary Figure 1.** Computational resolution enhancement cannot achieve single-cell level. The H&E image of human breast cancer with the enlarged tumor area (left) and xfuse-imputed gene expression summary with the enlarged gene expression summary in the tumor area (right).

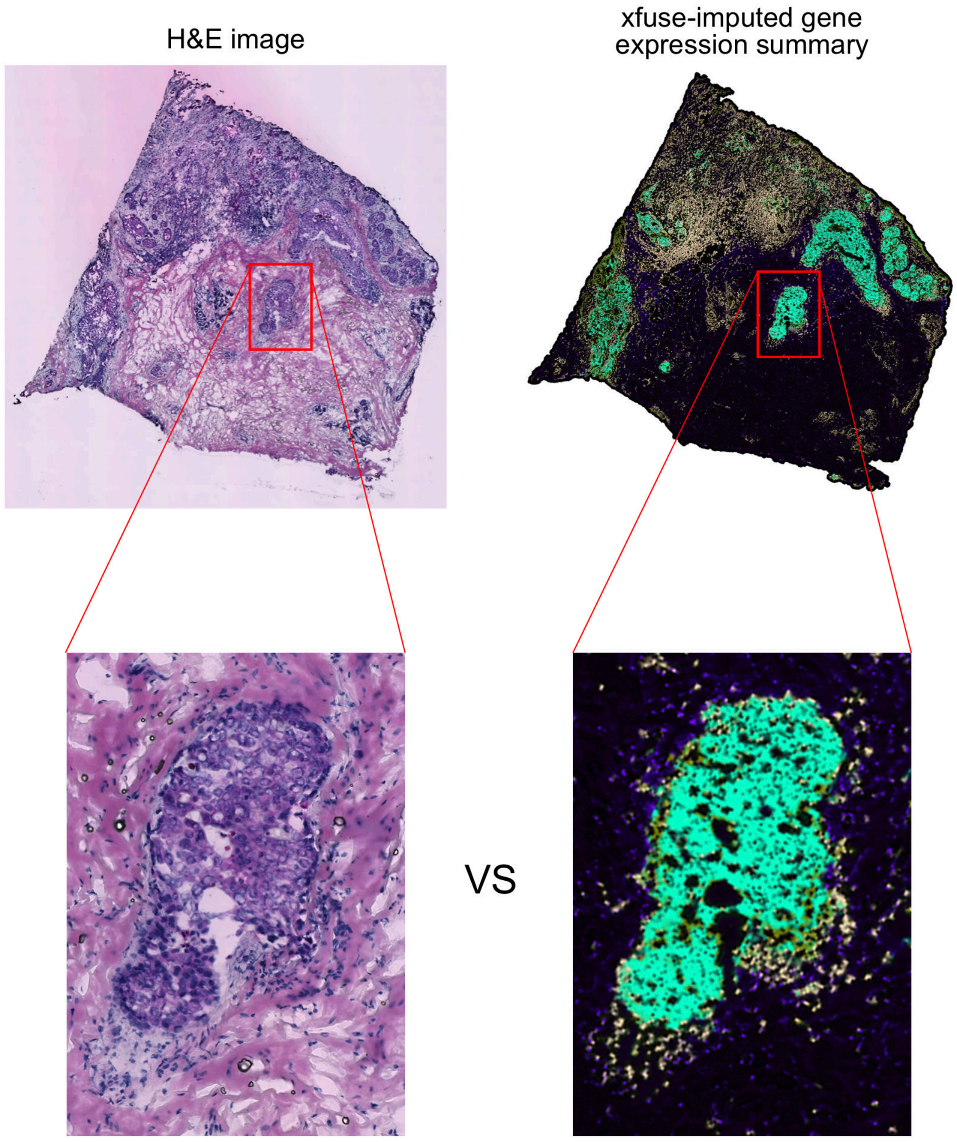

**Supplementary Figure 2.** H&E images of human breast cancer fresh tissue section in HDST (left) and nucleus segmentation (right). The middle panel is the enlarged region with nucleus segmentation overlaid (red circle).

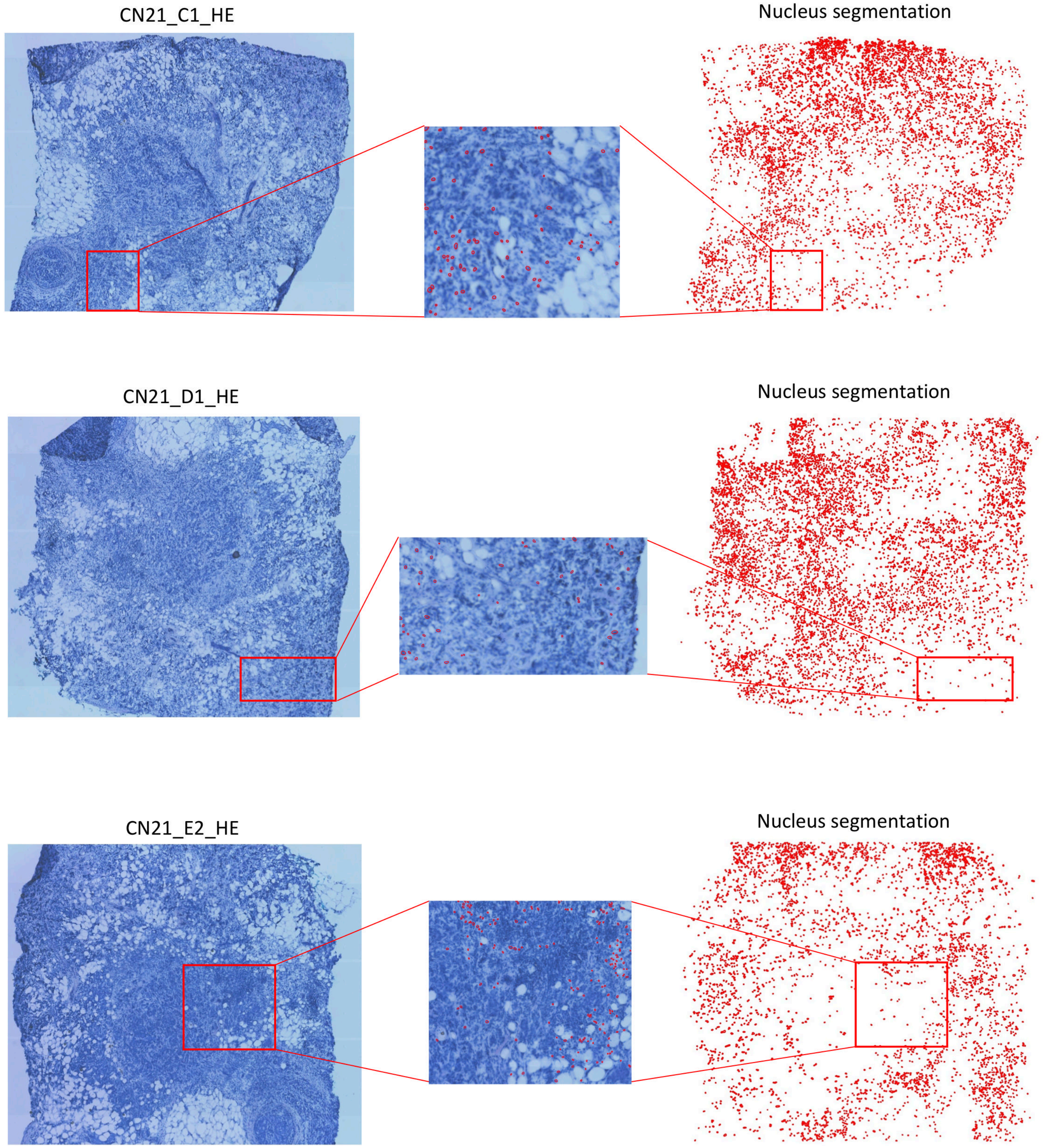

**Supplementary Figure 3.** Selection of λ on the simulation dataset and comparison with the other tools at the spot level. In the top panel, the boxplot represents the Pearson correlation (y-axis) between the true cell type proportion and the prediction using morphological features and gene expression. The first two boxplots in grey use only the morphological feature by K-means and gene expression by NNLS, respectively. The boxplots in white use both morphological features and gene expression by STIE at different λ (x-axis). Four scenarios were simulated based on the correlation between the ground truth and the K-means using the morphological feature (the first boxplot) and the correlation between the ground truth and the NNLS using the gene expression (the second boxplot): **a**, Morphology correlation (median=0.722) and gene expression correlation (median=0.708); **b**, Morphology correlation (median=0.643) and gene expression correlation (median=0.488); **c**, Morphology correlation (median=0.446) and gene expression correlation (median=0.708); **d**, Morphology correlation (median=0.390) and gene expression correlation (median=0.488). The red arrow points out the best parameter setting for STIE. The middle panel represents the RMSE of gene expression fitting, and the bottom panel represents the log-likelihood of the nuclear morphology.

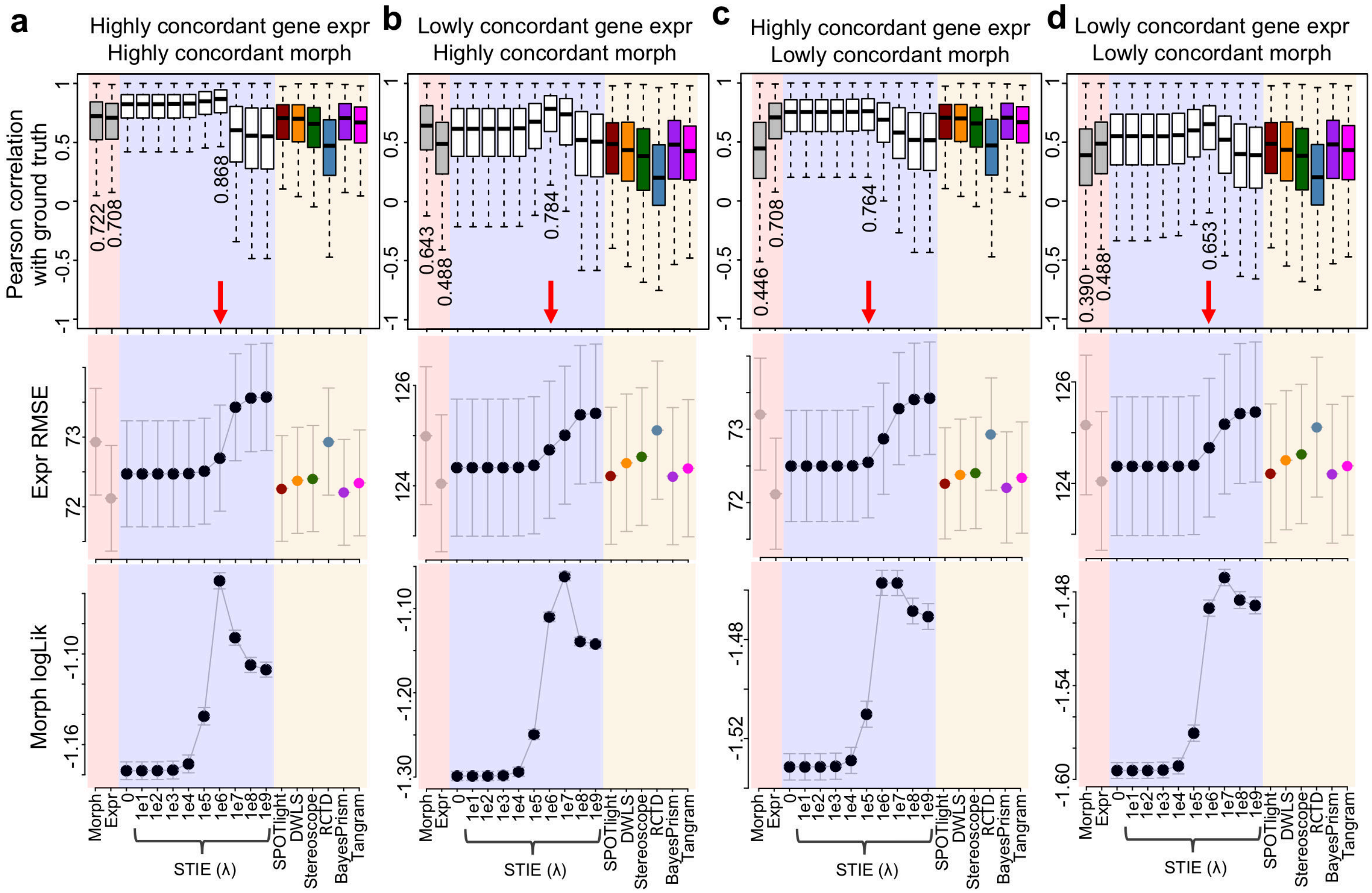

**Supplementary Figure 4.** Nuclear morphological feature selection on simulation data. There are 9 features used in this analysis: 3 true features (darkred), 3 intermediate features (steelblue) that are the sum of any two true features along with the random noise, and 3 irrelevant features (black) that are completely random noise. STIE was run under different values of λ. The panels from left to right represent the simulation data of four combinations between morphological features and spatial gene expression with either high or low noise. The upper panel represents the RMSE of gene expression fitting, and the bottom panel represents the log-likelihood of the nuclear morphology. The x-axis represents the incremental features from 1 feature to 9 features based on their RMSE ranking. The red ellipse indicate the correct feature selection.

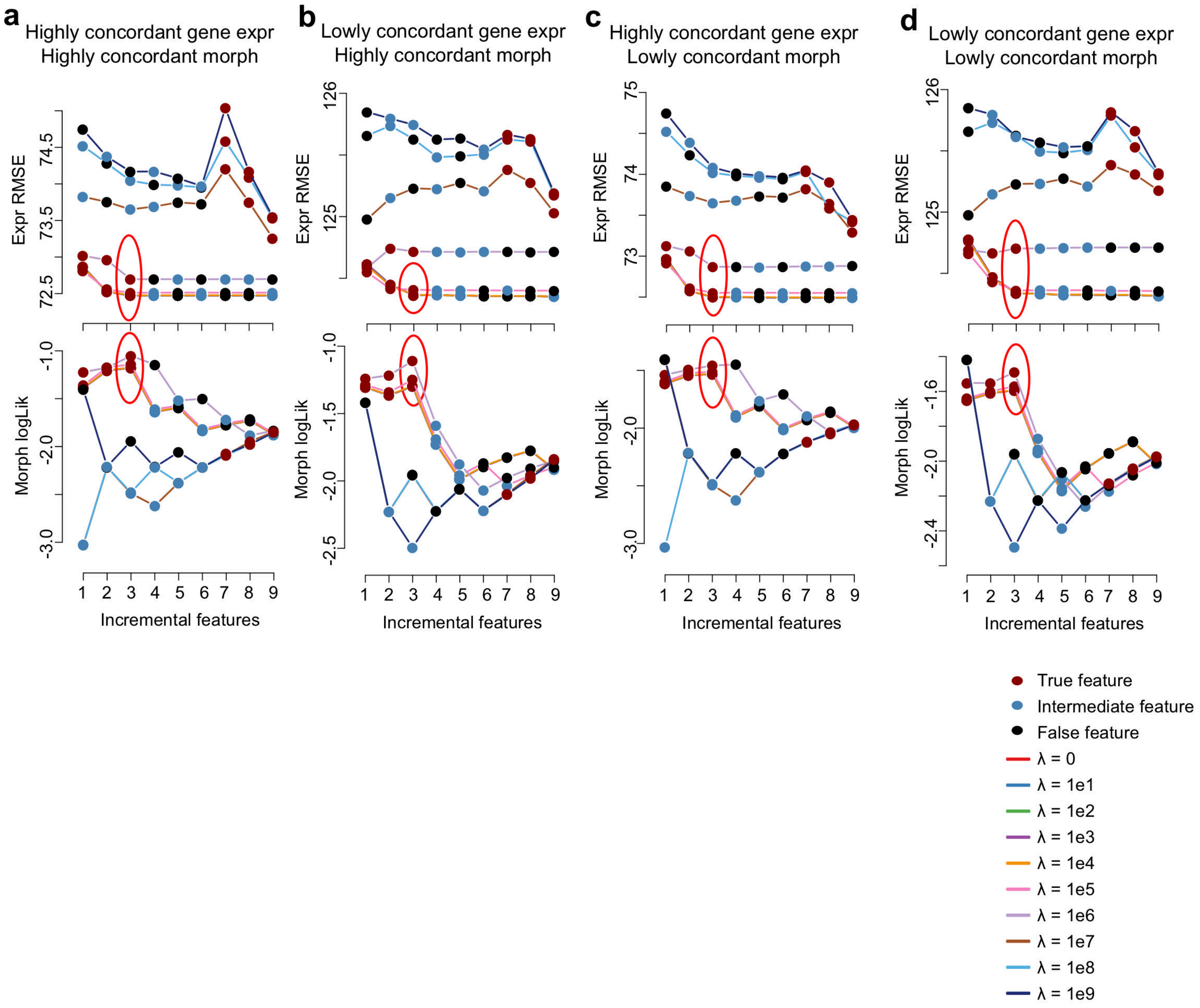

**Supplementary Figure 5.** Convergence analysis of STIE deconvolution on the simulation data. Three categories of parameters were analyzed: the non-negative regression coefficients on spatial gene expression (top panel), the mean (middle panel) and standard deviation (bottom panel) of morphological features under Gaussian distribution. The difference of parameter values (Y-axis) between the current and prior steps were calculated and plotted along the iteration steps (X-axis). The error bar represents the stand error.

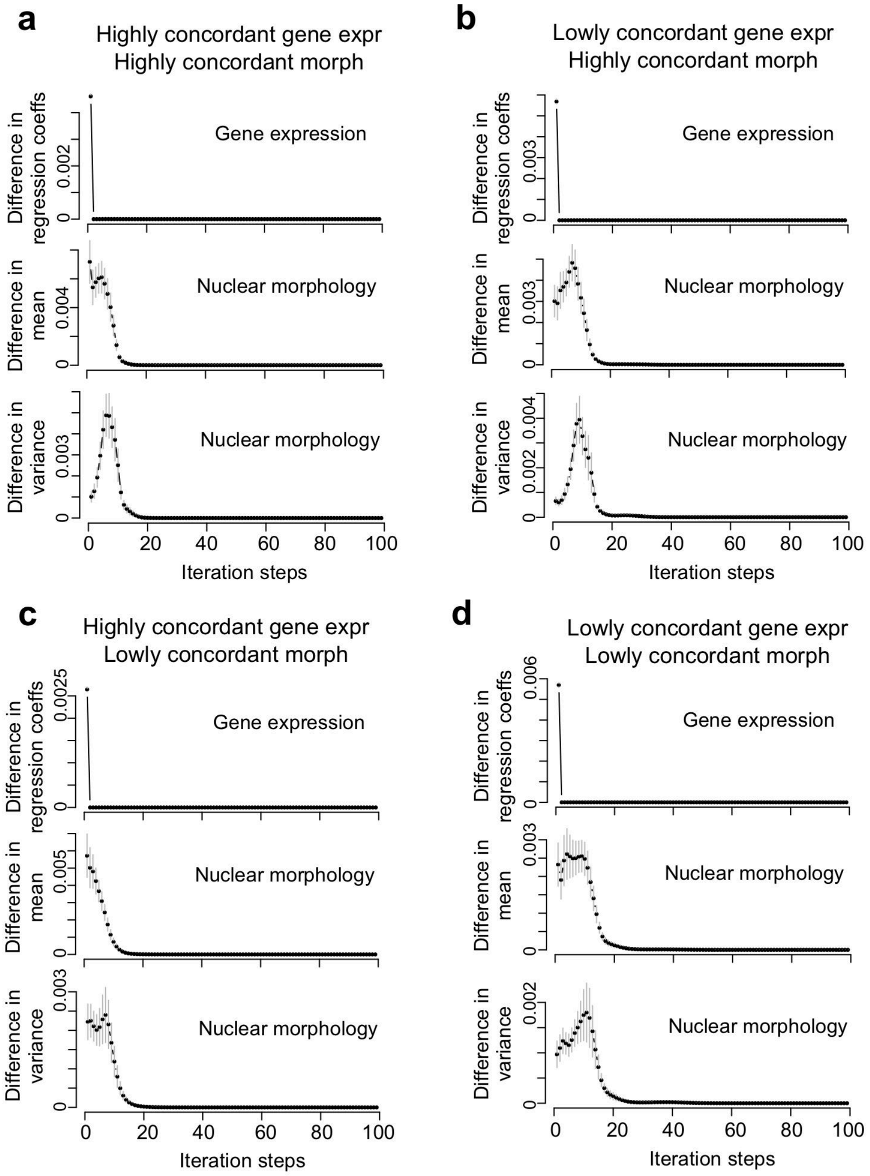

**Supplementary Figure 6.** Running time of STIE deconvolution on the simulation data over different numbers of spots (**a**), cell types (**b**), and marker genes (**c**). **a**, Given the number of spots, 1,000 marker genes, 10 cell types, and the single cells of 10x number of spots were simulated. **b**, Given the number of cell types, 1,000 marker genes, 1,000 spots, and 10,000 single cells were simulated. **c**, Given the number of marker genes, 10 cell types, 1,000 spots, and 10,000 single cells were simulated. The running time is estimated using the R function system.time().

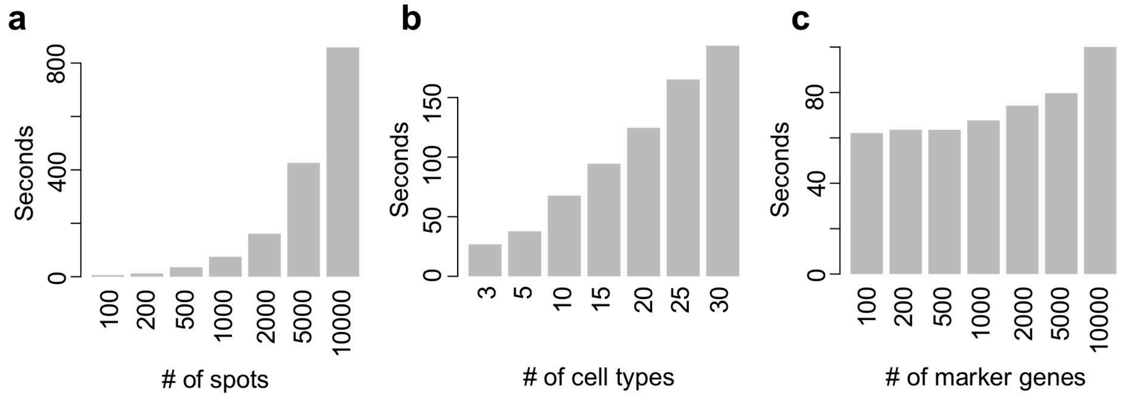

**Supplementary Figure 7.** Correlation heatmap between nuclear morphological features in the real dataset, **a**, 10X Visium mouse brain FFPE, **b**, 10X Visium human breast cancer FFPE, and **c-d**, 10X V2 Chemistry CytAssist mouse brain section 1&2. Each item represents the R^2 between two features, with the size of circles proportional to the values.

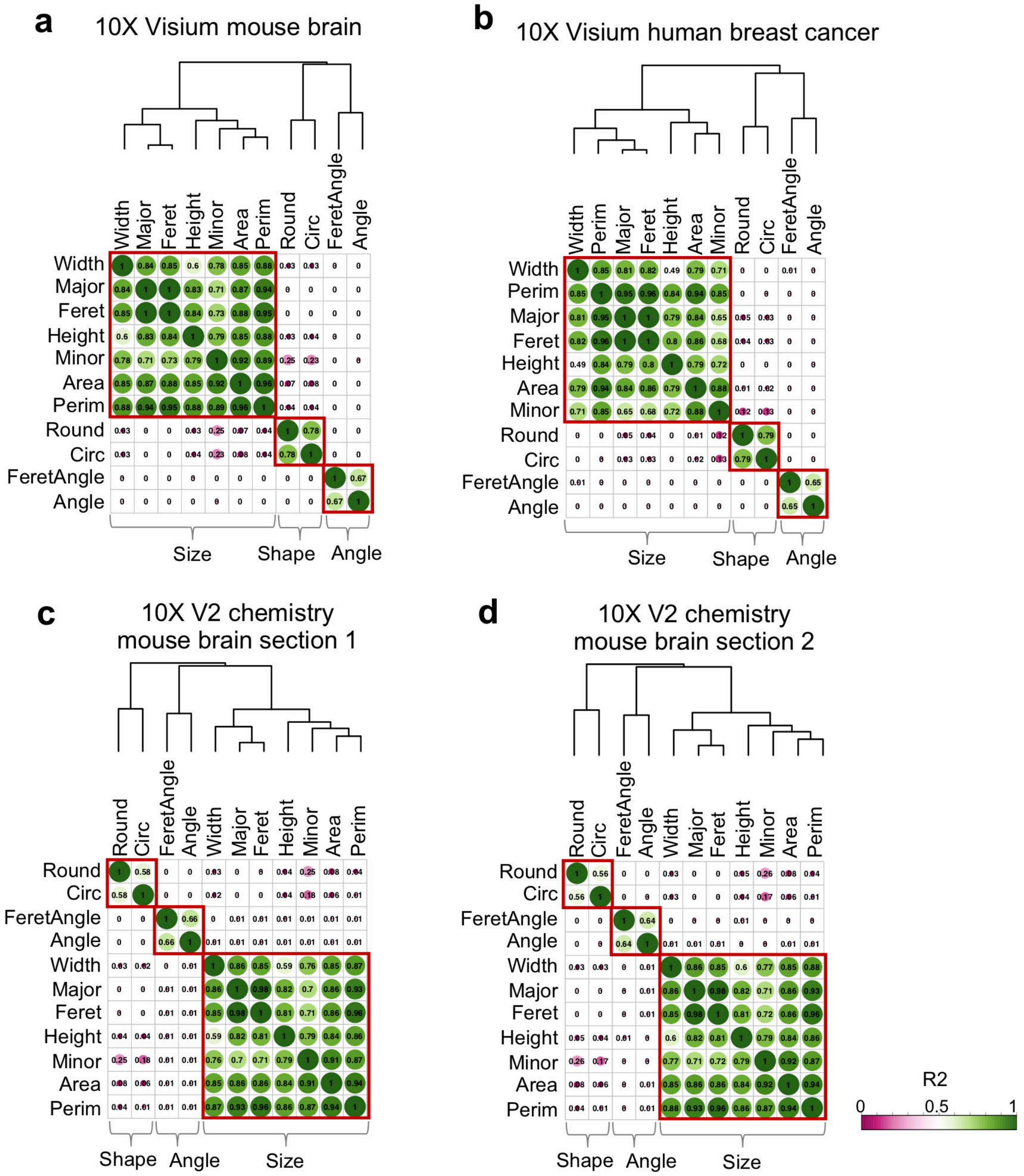

**Supplementary Figure 8.** Selection of nuclear morphological features and λ via grid search for the mouse brain hippocampus. **a-c**, STIE deconvolution on 10X Visium mouse brain hippocampus FFPE (**a**), 10X Visium V2 Chemistry CytAssist mouse brain hippocampus FFPE section 1 (**b**) and section 2 (**c**). **d-f**, STIE clustering on 10X Visium mouse brain hippocampus FFPE (**d**), 10X Visium V2 Chemistry CytAssist mouse brain hippocampus FFPE section 1 (**e**) and section 2 (**f**). The selected morphological feature and λ are marked in the circle.

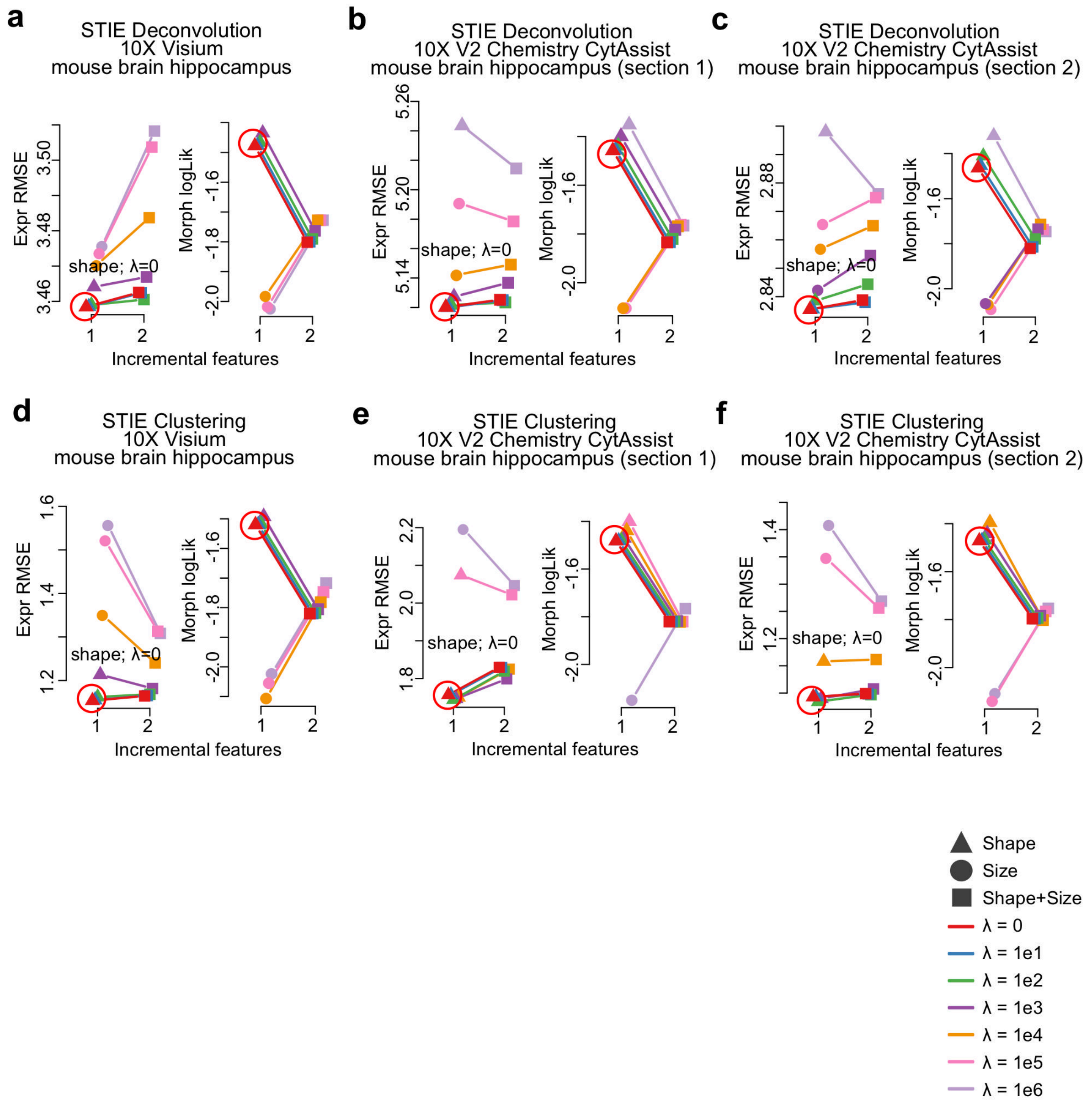

**Supplementary Figure 9.** Selection of nuclear morphological features and λ via grid search for the mouse brain cortex. **a-d**, STIE clustering on 10X Visium mouse brain cortex FFPE (**a**), 10X Visium V2 Chemistry CytAssist mouse brain cortex FFPE section 1 (**b**) and section 2 (**c**), and 10X Visium fresh frozen human brain cortex (**d**). The selected morphological feature and λ are marked in the circle.

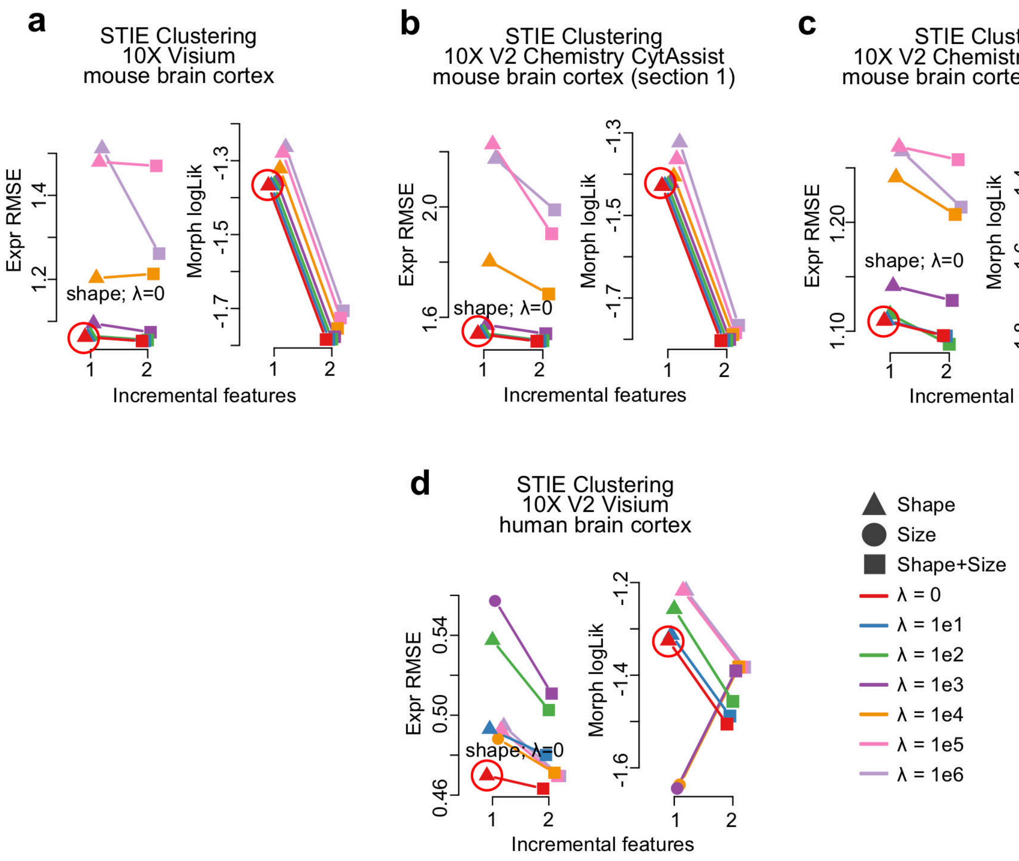

**Supplementary Figure 10.** Selection of nuclear morphological features and λ via grid search for the human breast cancer. **a**, STIE deconvolution on 10X Visium human breast cancer FFPE. **b**, STIE clustering on 10X Visium human breast cancer FFPE. The selected morphological feature and λ are marked in the circle.

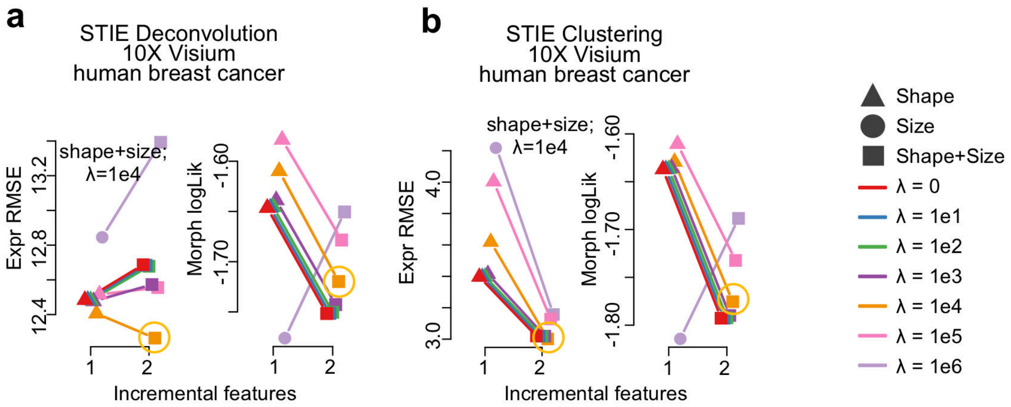

**Supplementary Figure 11.** Running time and memory usage of methods on the real dataset. Each job was submitted to the slurm queue, and the sacct command was used to display the running time and maximum memory size. The following STIE running time is based on the predefined nuclear morphological features and λ. When selecting morphological features and λ, STIE_search may take longer time (~10X) depending on the searching space but may hold the similar memory usage.

**a** Deconvolution on 10X Visium mouse brain hippocampus FFPE

| | Time | Memory(max) |
|---|---|---|
| SPOTlight | 04m55s | 2,415M |
| DWLS | 12m55s | 3,362M |
| Stereoscope | 04h38m36s | 5,648M |
| RCTD | 53s | 3,718M |
| Tangram | 54s | 7,224M |
| BayesPrism | 04m45s | 4,298M |
| STIE | 44s | 11,684M |

**b** Clustering on 10X Visium mouse brain hippocampus FFPE

| | Time | Memory(max) |
|---|---|---|
| BayesSpace | 07m19s | 3,374M |
| SpaGCN | 54s | 7,327M |
| MUSE | 02m54s | 971M |
| STIE | 02m05s | 10,989M |

**c** Deconvolution on the 10X Visium human breast cancer FFPE

| | Time | Memory(max) |
|---|---|---|
| SPOTlight | 04m28s | 5,202M |
| DWLS | 05h07m48s | 3,994M |
| Stereoscope | 1d07h50m37s | 3,207M |
| RCTD | 33m27s | 9,749M |
| Tangram | 16m50s | 12,507M |
| BayesPrism | 04h57m09s | 22,246M |
| STIE | 8m24s | 12,916M |

**d** Clustering on the 10X Visium human breast cancer FFPE

| | Time | Memory(max) |
|---|---|---|
| BayesSpace | 17h01m38s | 17,987M |
| SpaGCN | 01m03s | 14,525M |
| MUSE | 35m14s | 975M |
| STIE | 16m52s | 15,271M |

**Supplementary Figure 12.** The similarity between scRNA-seq derived cell type transcriptomic signatures in the mouse brain hippocampus.

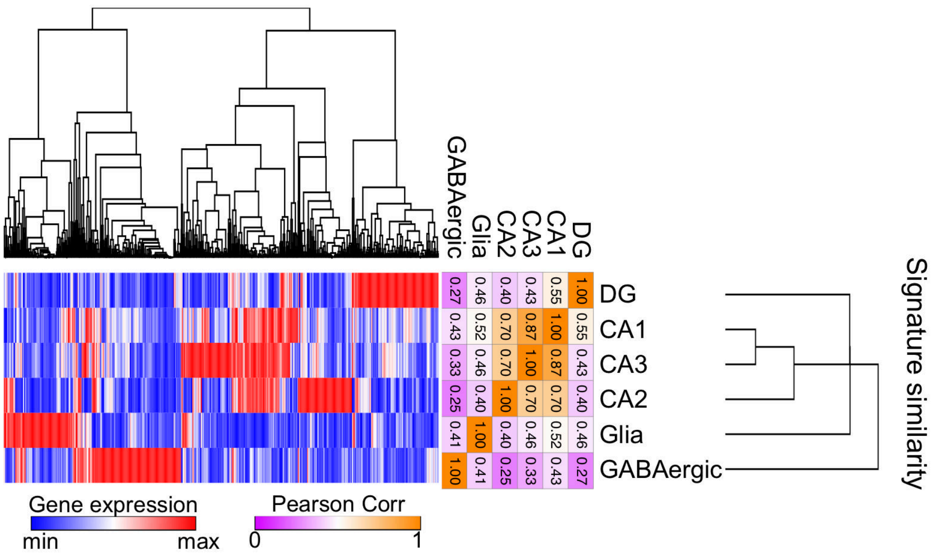

**Supplementary Figure 13**. Investigation of marker gene selection on STIE deconvolution. **a-c.** STIE deconvolution using new transcriptomic signatures constructed by SPOTlight. **a**. Overlapping between new and original marker genes. **b**. STIE deconvolution using new transcriptomic signatures. **c**. Concordance of STIE deconvolution between using original and new transcriptomic signatures. **d-e**. Concordance of STIE deconvolution between using randomly sampled marker genes and original full marker genes. **d**. STIE deconvolution on the gene expression and original morphological information. **e.** STIE deconvolution on the gene expression and the shuffled morphological information.

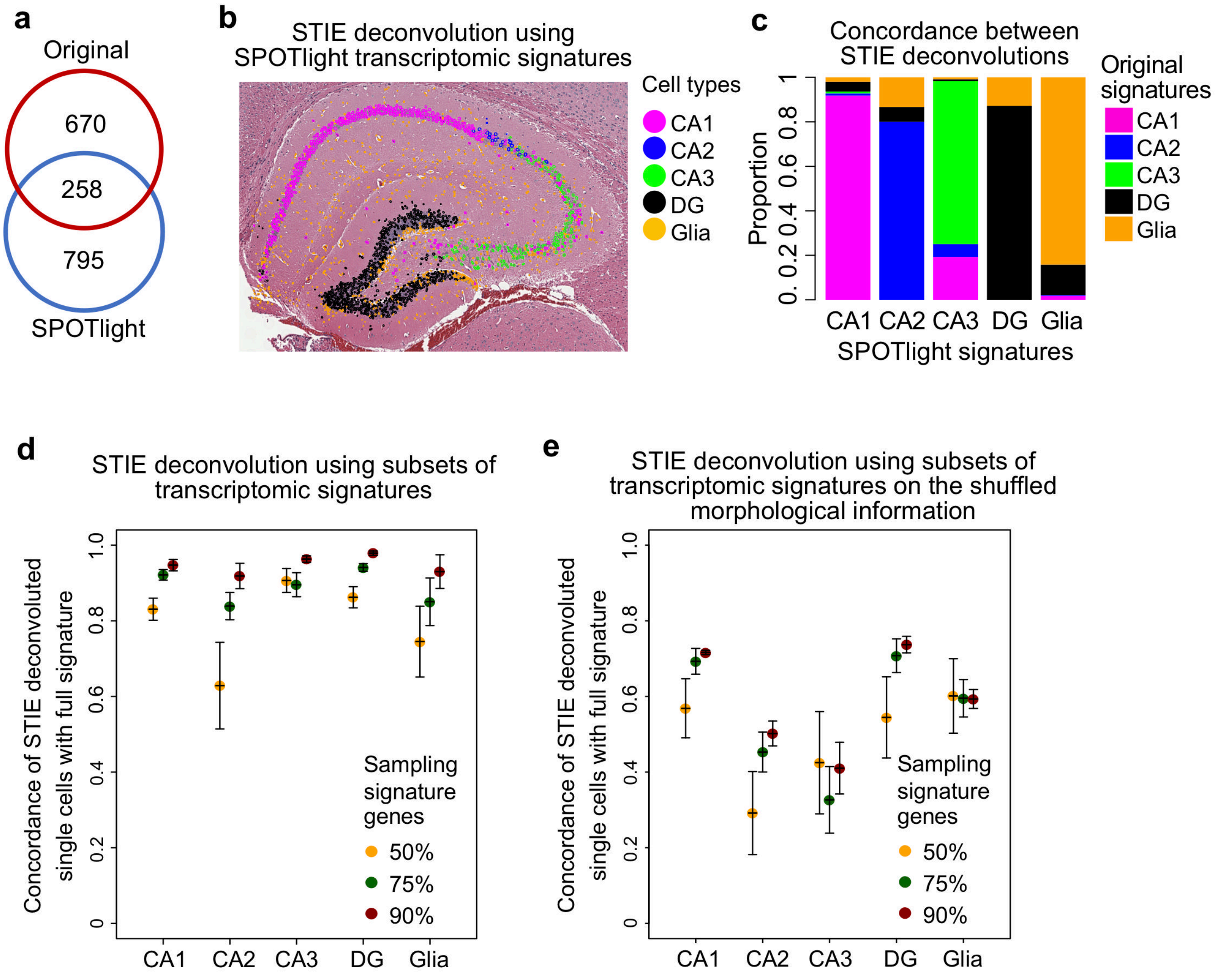

**Supplementary Figure 14.** The influence of image alignment to STIE deconvolution on the human breast cancer 10X Visium FFPE. The image was shifted to misalign to the spot. We rerun STIE deconvolution and evaluate its concordance with the STIE deconvolution under accurate alignment. It shows 88% and 84% concordance with the accurate alignment, even if the cells are misplaced by a half spot and a whole spot, respectively. A slight misalignment, 10% spot size, does not significantly influence the cell typing (98% concordance). The x-axis represents the shift distance of the image. The y-axis represents the single-cell level concordance of the STIE deconvoluted cell types with that under accurate alignment.

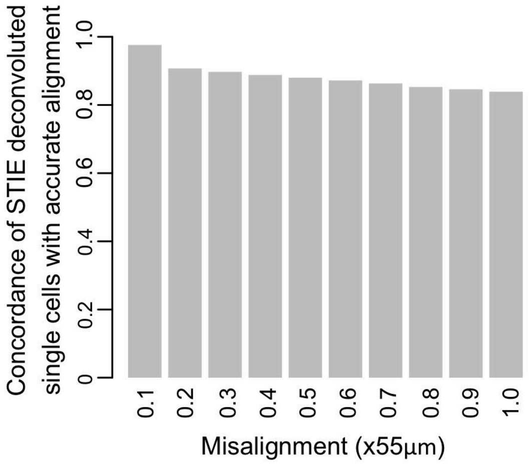

**Supplementary Figure 15.** Deconvoluted cell type proportion of 10X Visium human breast cancer FFPE spatial transcriptomics by SPOTlight, DWLS, Stereoscope, RCTD, Tangram, and BayesPrism. The piecharts are overlaid on the tissue slide. The small area is enlarged on the top right of each image. The color refers to the cell type.

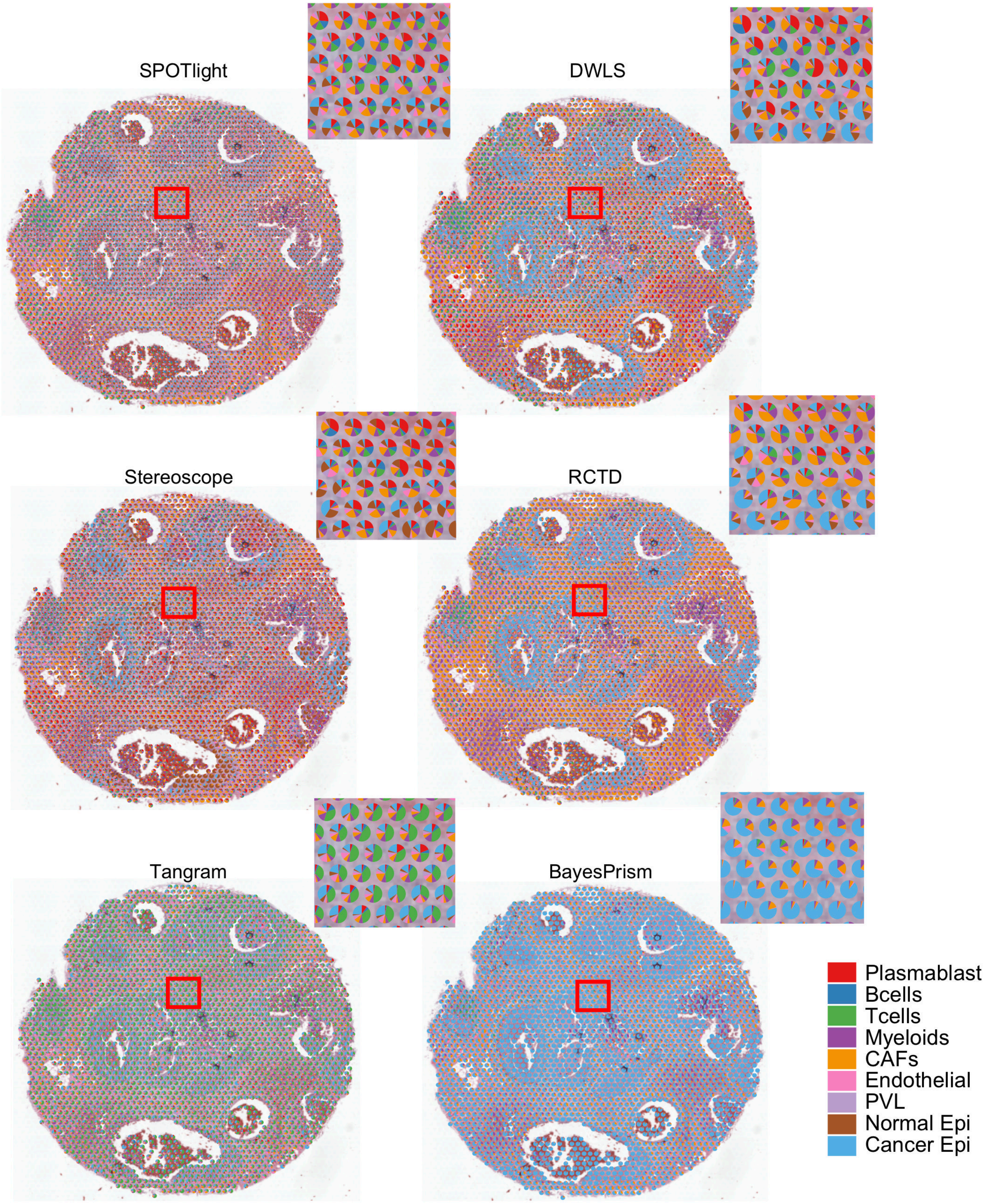

**Supplementary Figure 16.** Concordance of the tumor region between the prediction and the ground truth measured by Adjusted Rand Index (ARI) (**b**) and R2 (**c**). **a.** Refined manual annotation of tumor region. **b**. Different cutoffs of cell type proportion in spots were applied to determine the tumor region at spot level. The binary value of tumor regions was compared with the manual annotation to calculate the ARI. The best ARI value was reported in the legend. **c**. The R2 between the continuous value of Cancer Epithelial cell proportion and the manual annotation.

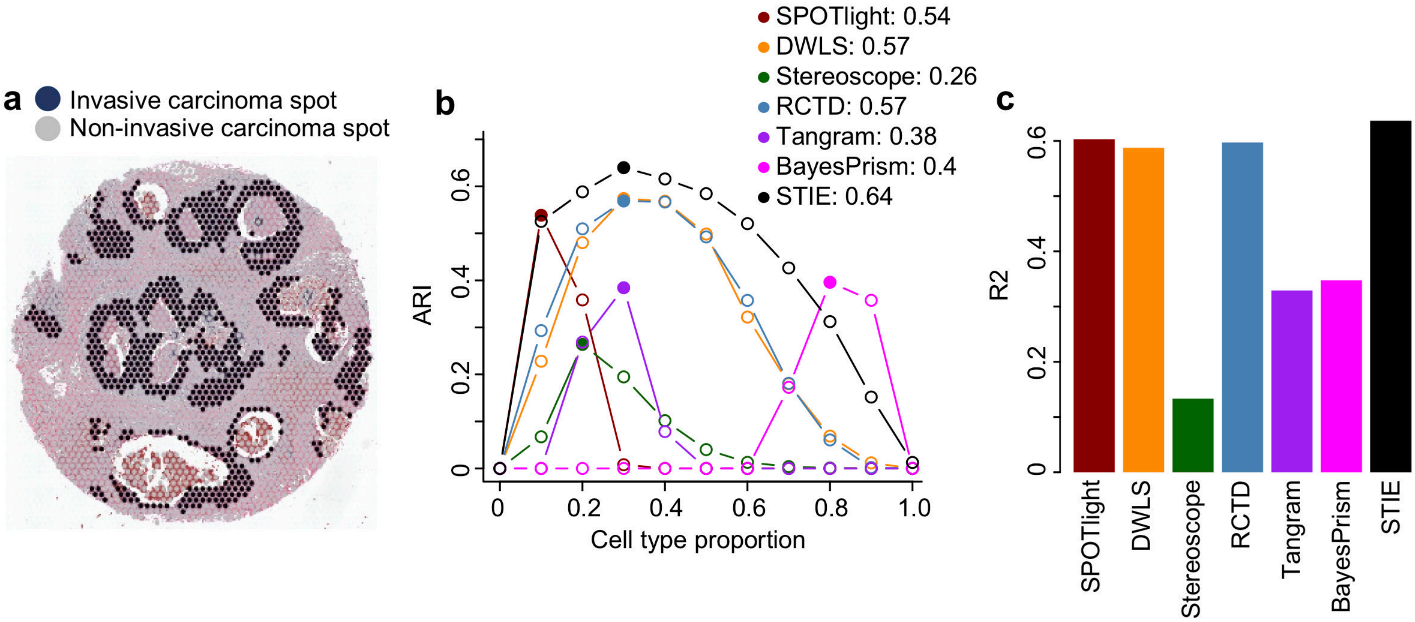

**Supplementary Figure 17** Single-cell level cell type signature learning from 10X Visium human breast cancer FFPE spatial transcriptomics data. The signatures were deconvoluted based on the scRNA-seq derived transcriptomic signatures via NNLS and DWLS.

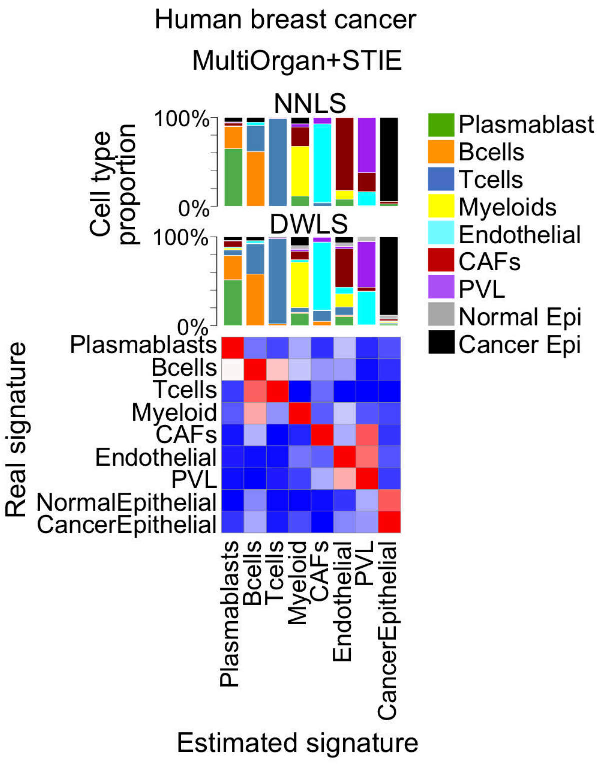

**Supplementary Figure 18.** Spot level K-means clustering on 10X Visium mouse brain hippocampus FFPE.

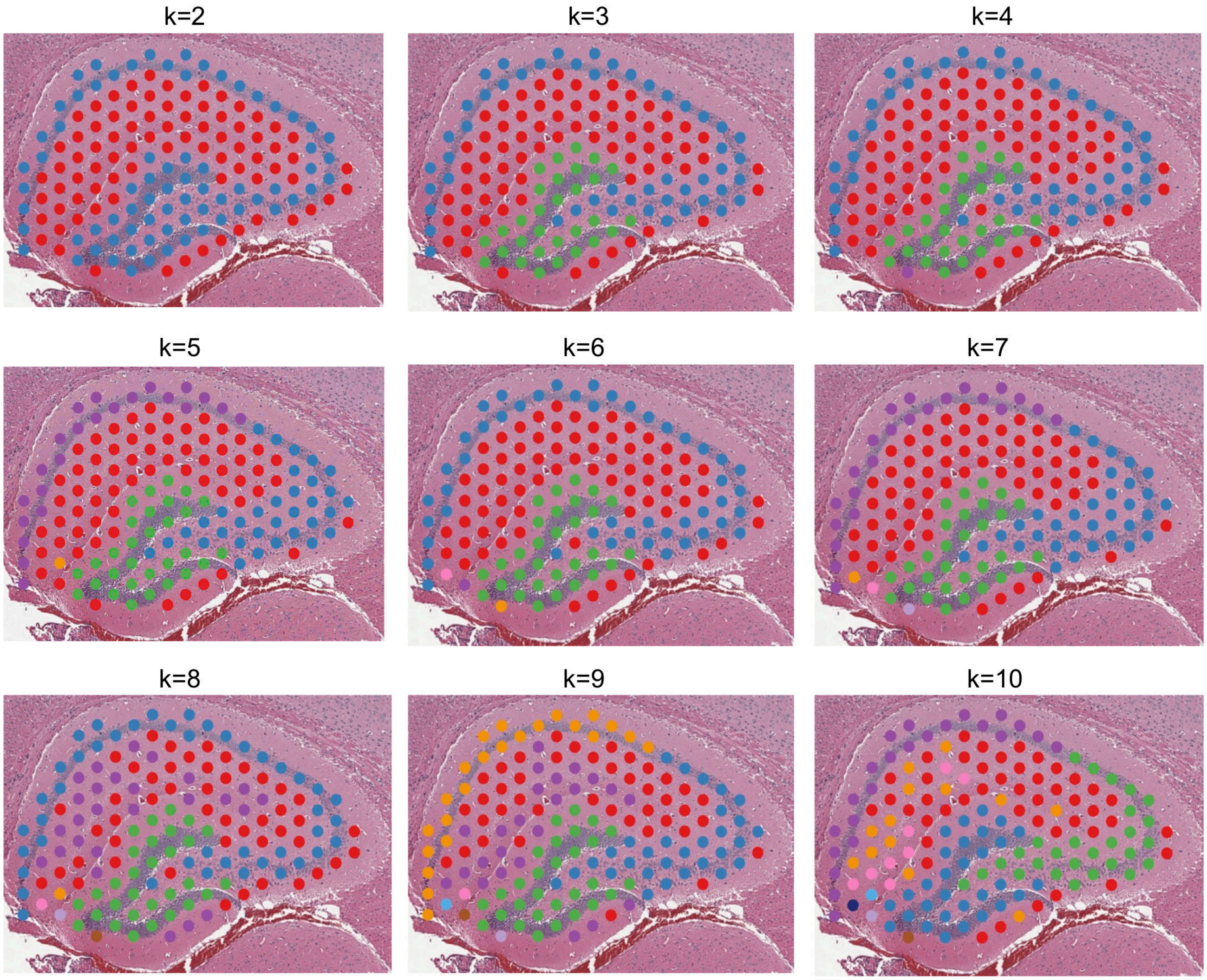

**Supplementary Figure 19.** Spot level SpaGCN clustering on 10X Visium mouse brain hippocampus FFPE.

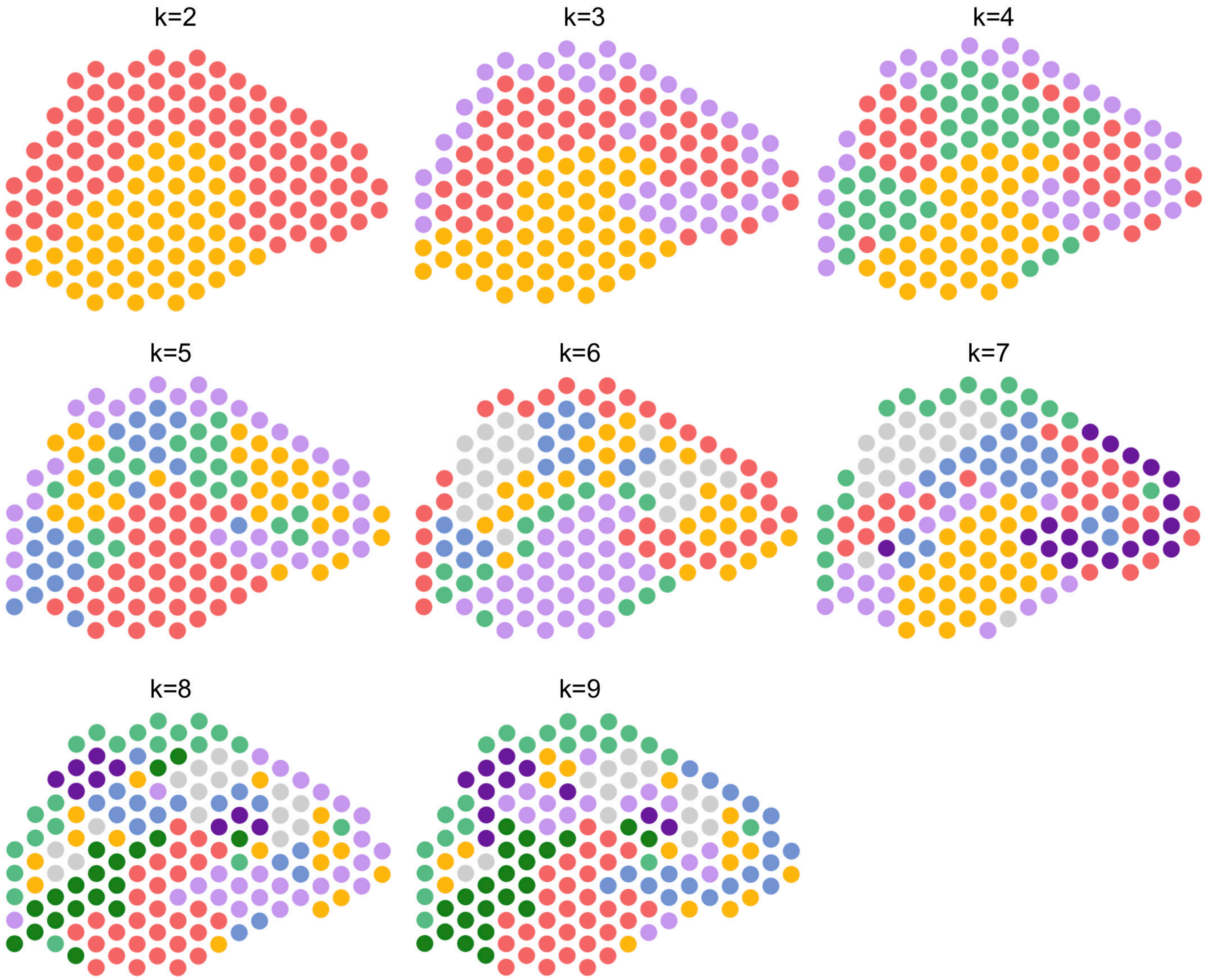

**Supplementary Figure 20.** Subspot level BayesSpace clustering on 10X Visium mouse brain hippocampus FFPE.

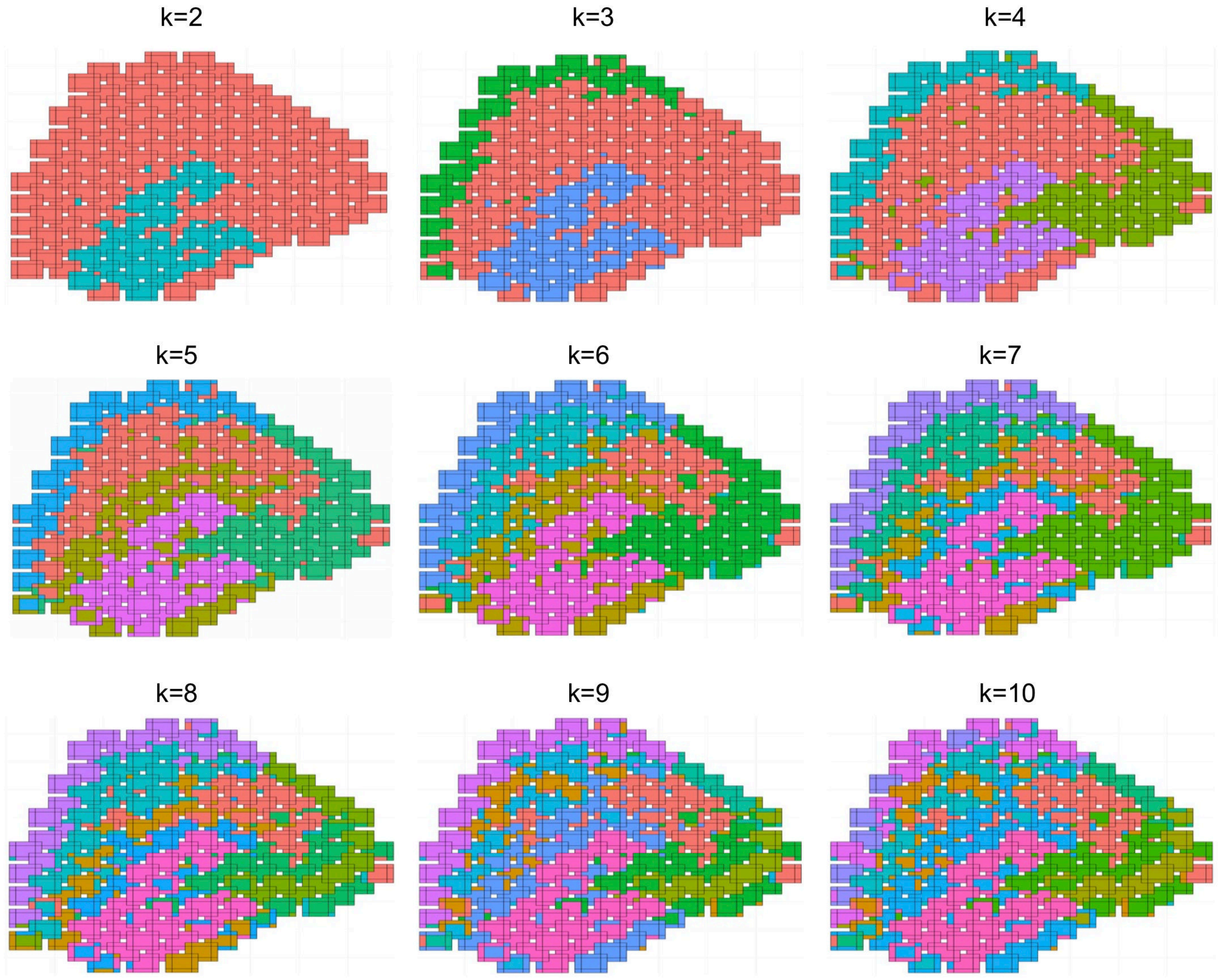

**Supplementary Figure 21.** Single-cell level STIE clustering on 10X Visium mouse brain hippocampus FFPE.

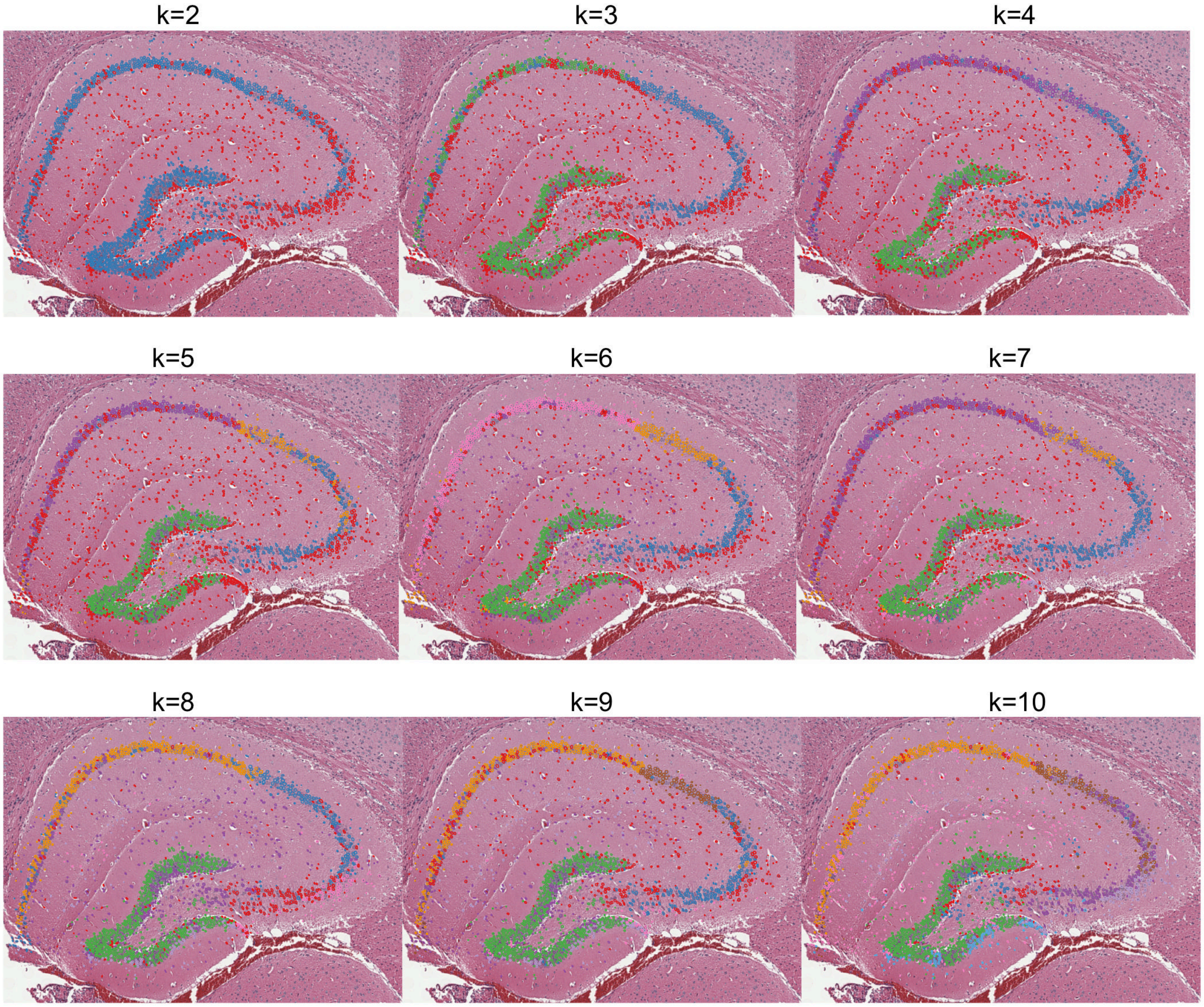

**Supplementary Figure 22.** Clustering on 10X V2 Chemistry CytAssist FFPE mouse brain hippocampus at k=5. **a.** Spot-level K-means. **b.** Single-cell level STIE clustering. **c.** The CAGEs by K-means and STIE were deconvoluted based on the scRNA-seq derived transcriptomic signatures using NNLS and DWLS for section 1 (top panel) and section 2 (bottom panel).

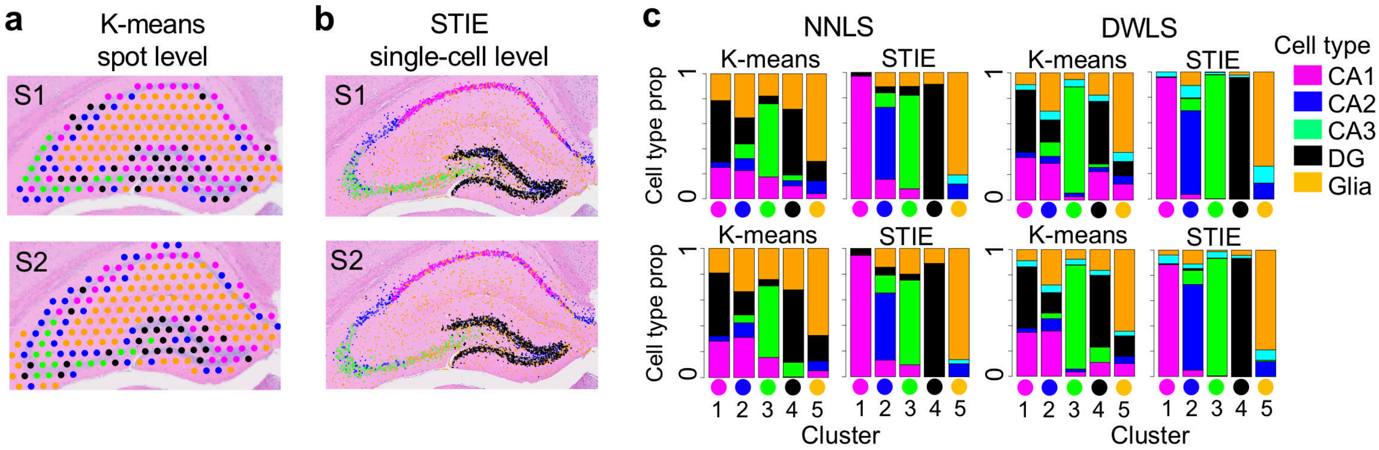

**Supplementary Figure 23.** Spot level K-means clustering on 10X Visium mouse brain cortex FFPE.

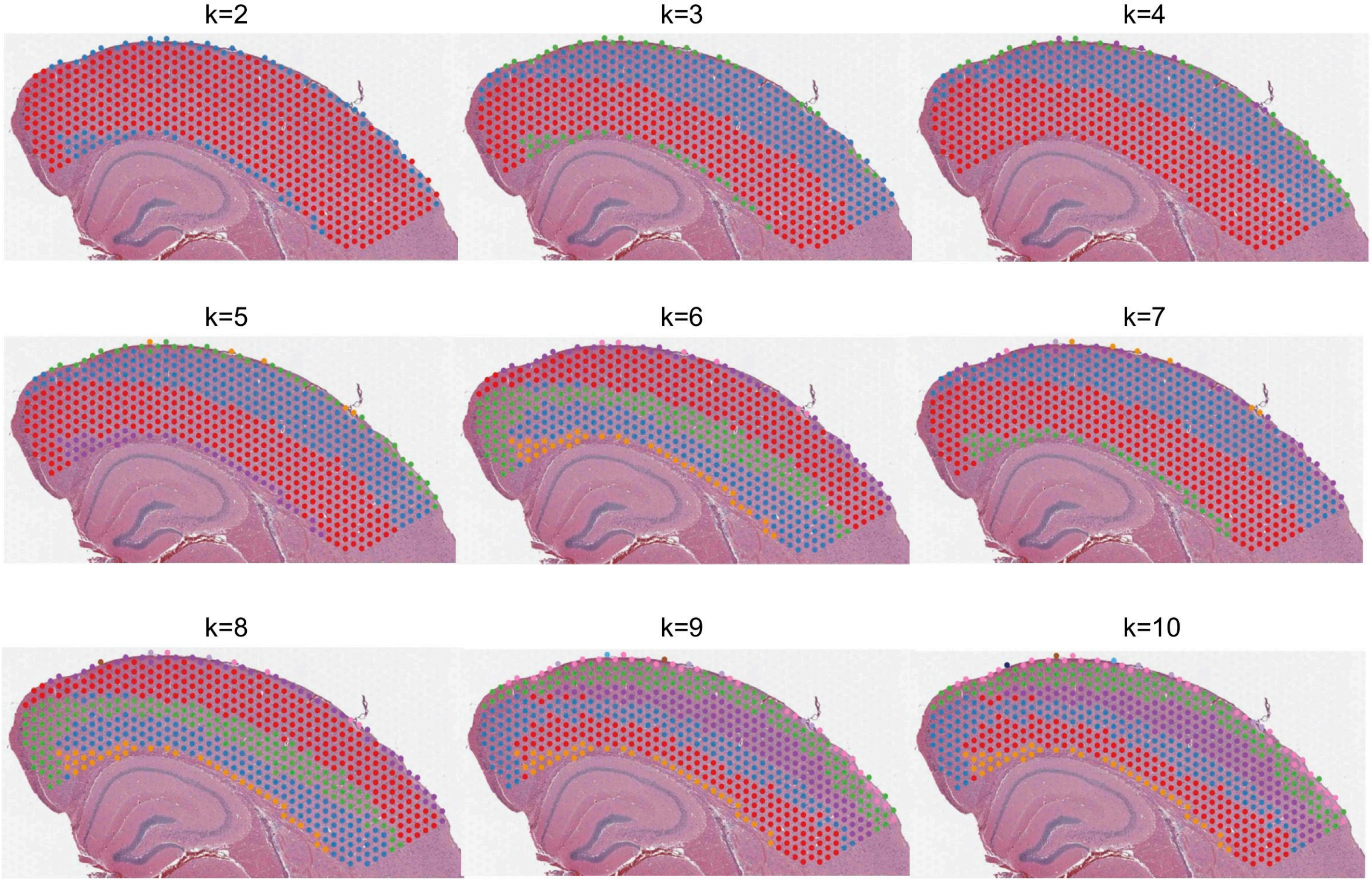

**Supplementary Figure 24.** Spot level SpaGCN clustering on 10X Visium mouse brain cortex FFPE.

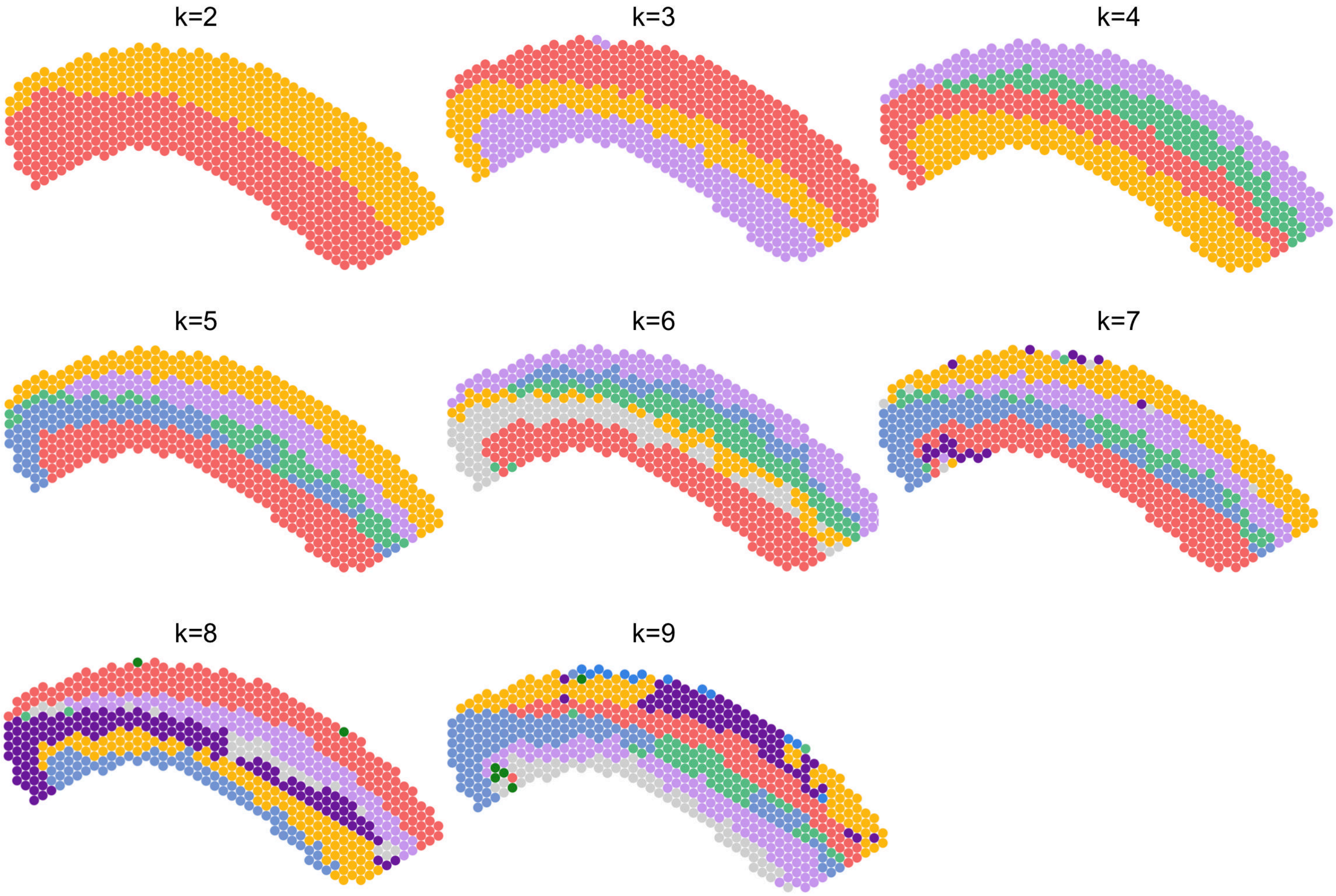

**Supplementary Figure 25.** Subsppot level BayesSpace clustering on 10X Visium mouse brain cortex FFPE.

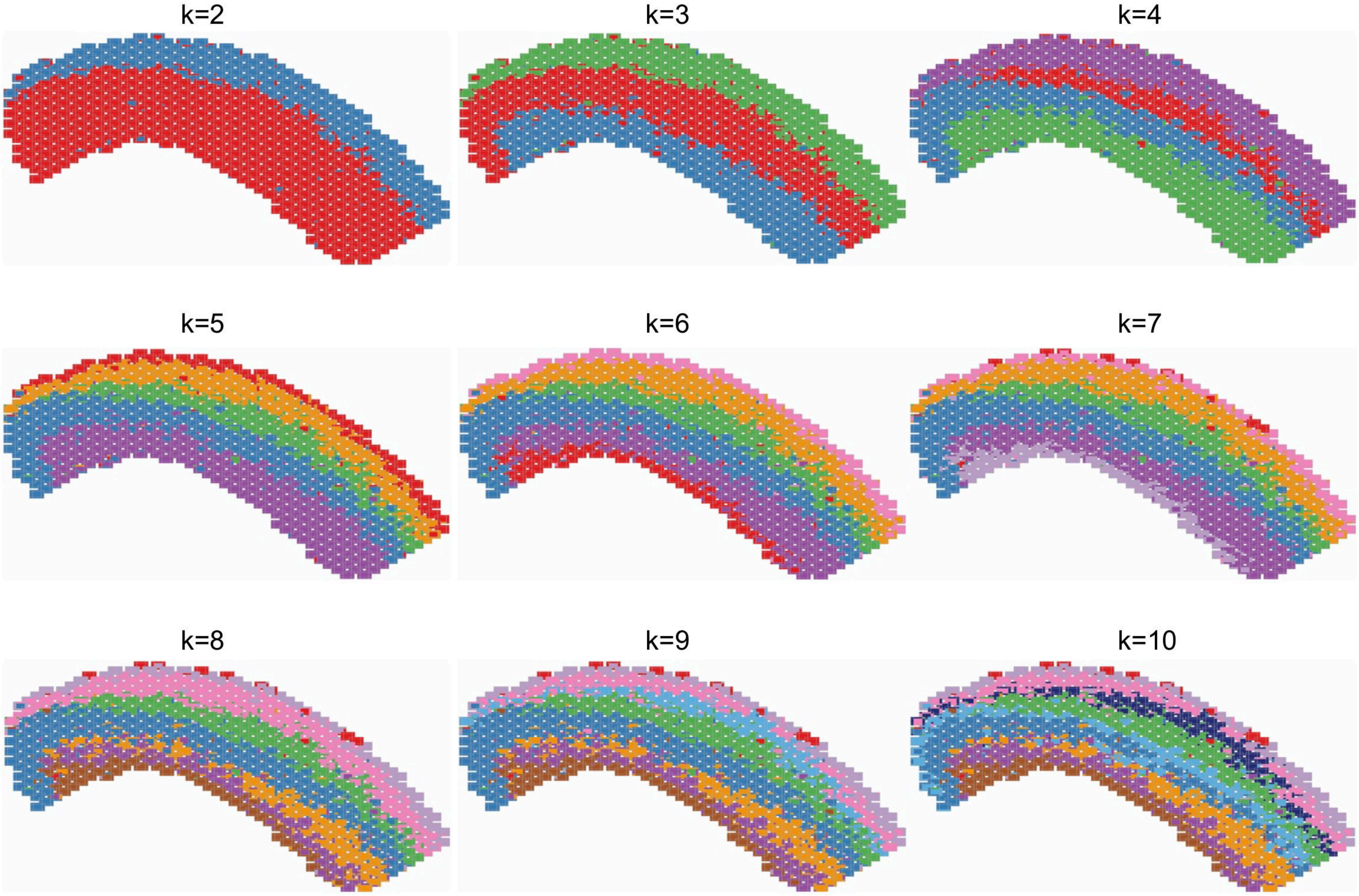

**Supplementary Figure 26.** Single-cell level STIE clustering on 10X Visium mouse brain cortex FFPE.

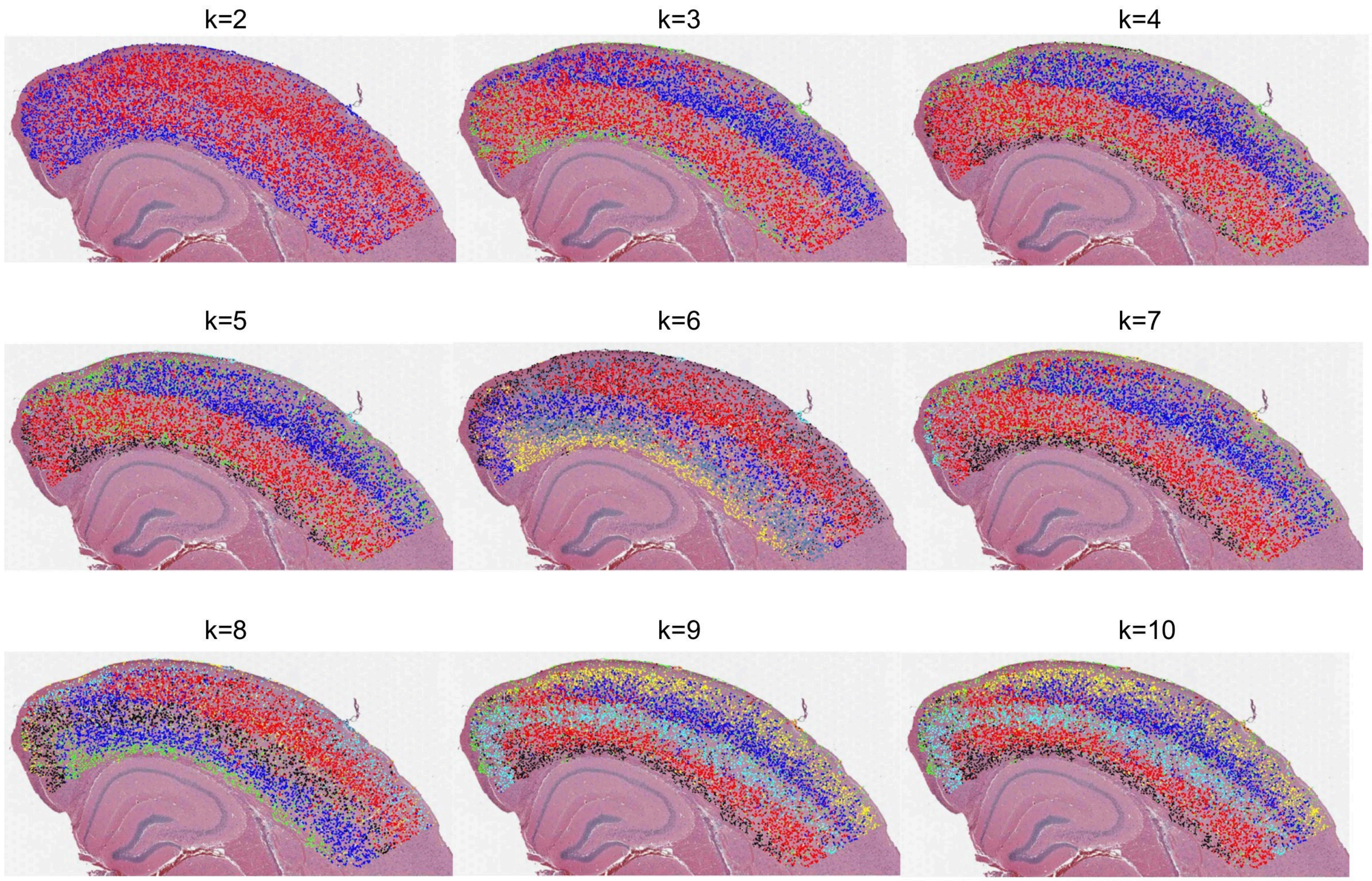

**Supplementary Figure 27.** Clustering on 10X V2 Chemistry CytAssist FFPE mouse brain cortex at k=6. **a&c.** Spot-level K-means (left) and single-cell level STIE clustering (right) for section 1 (**a**) and section 2 (**c**). **b&d.** The CAGEs by K-means and STIE were deconvoluted based on the scRNA-seq derived transcriptomic signatures using NNLS for section 1 (**b**) and section 2 (**d**).

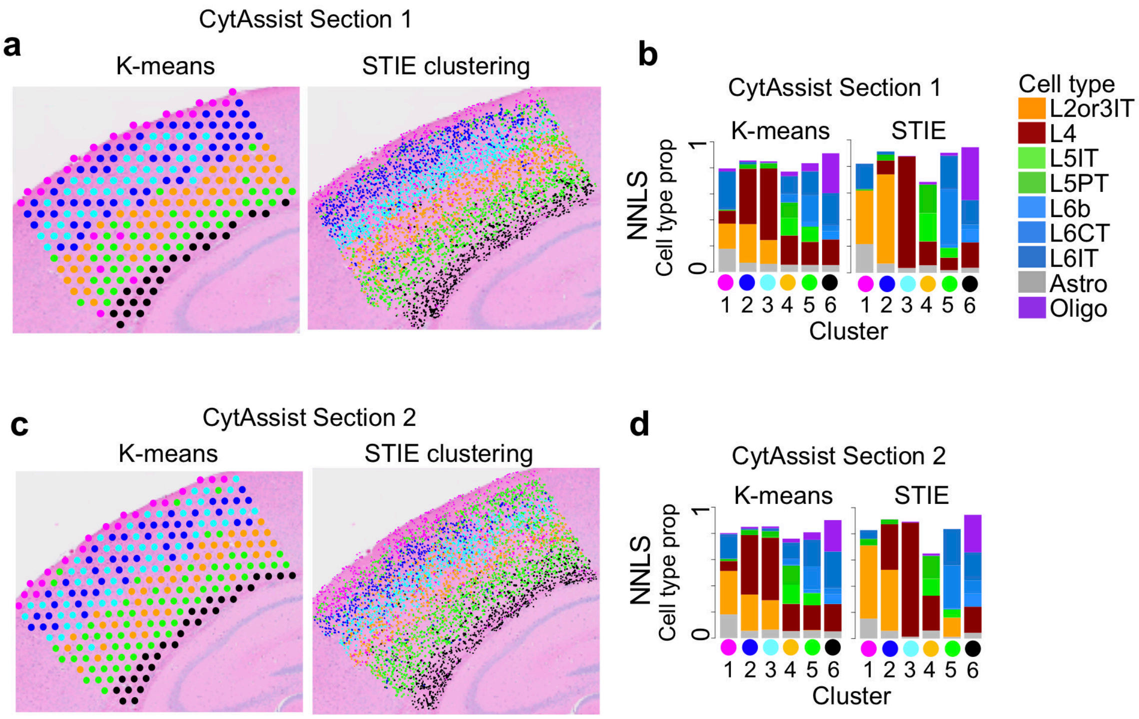

**Supplementary Figure 28.** STIE ("shape" as the morphological feature) on the human brain frozen section and mouse brain FFPE. **a.** The single-cell level clustering by STIE on the human brain frozen section. **b.** Manual annotation. **c.** K-means on human brain frozen section only using gene expression at the spot level. **d.** STIE on human brain frozen section at spot level. **e.** The K-means (spot level) and STIE (single-cell level) clustering on the mouse brain FFPE.

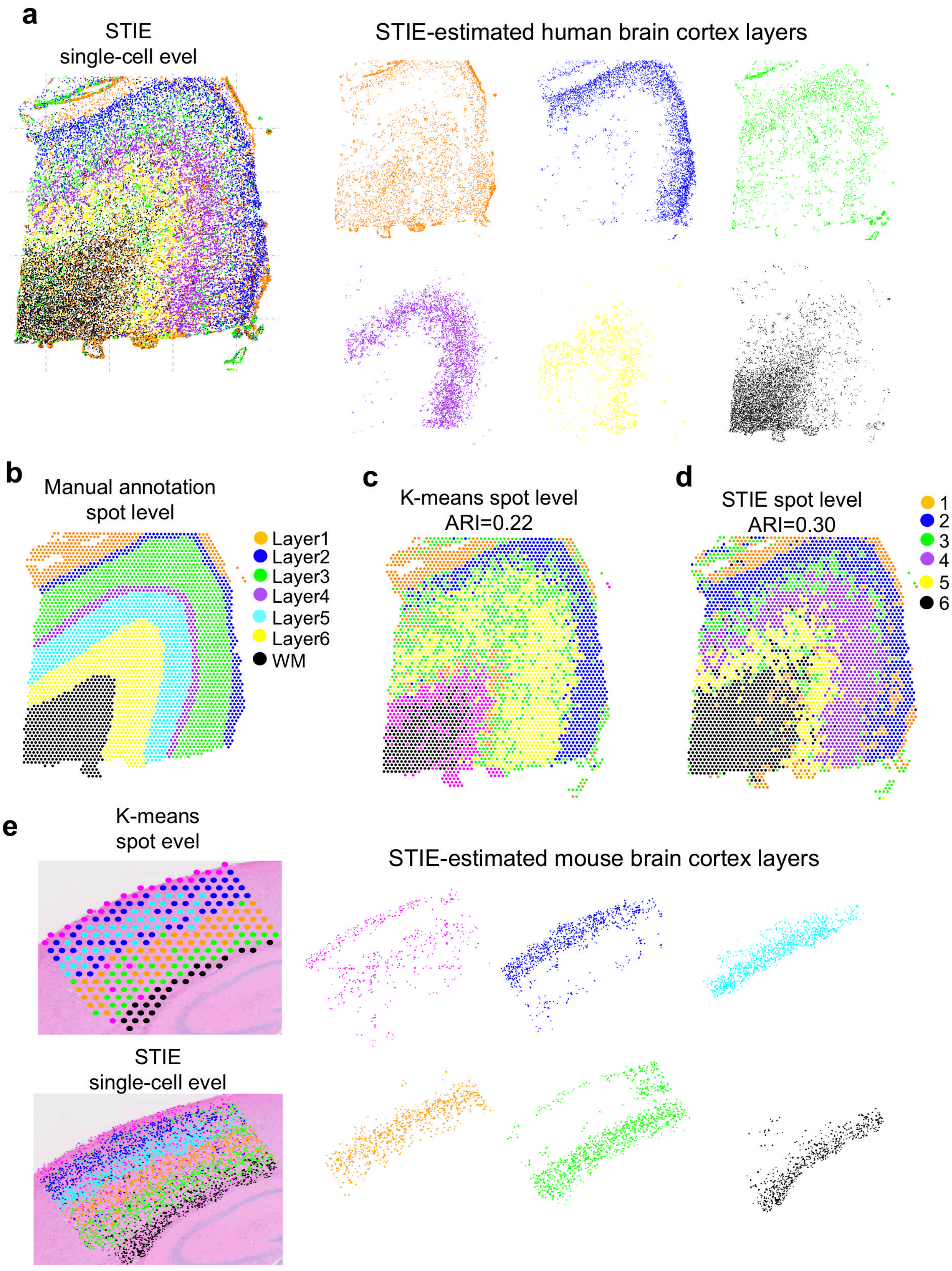

**Supplementary Figure 29.** STIE ("shape+size" as the morphological feature) on the human brain frozen section and mouse brain FFPE. **a-b.** The STIE clustering on the human brain frozen section at single-cell level (**a**) and spot level (**b**). **d.** The STIE clustering on mouse brain FFPE section at single-cell level. **c&e**. The concordance table of STIE cell typing at the single cell level between using "shape" and "shape+size" as morphological features on human brain frozen section (**c**) and mouse brain FFPE (**e**). **d.** The STIE clustering on mouse brain FFPE section at single-cell level. **f.** The STIE single-cell level clustering using SpaGCN spot-level clustering as initial values; (c) Downgrading from the single-cell level to the spot level (ARI=0.5 compared with manual annotation). Each color indicates one cluster.

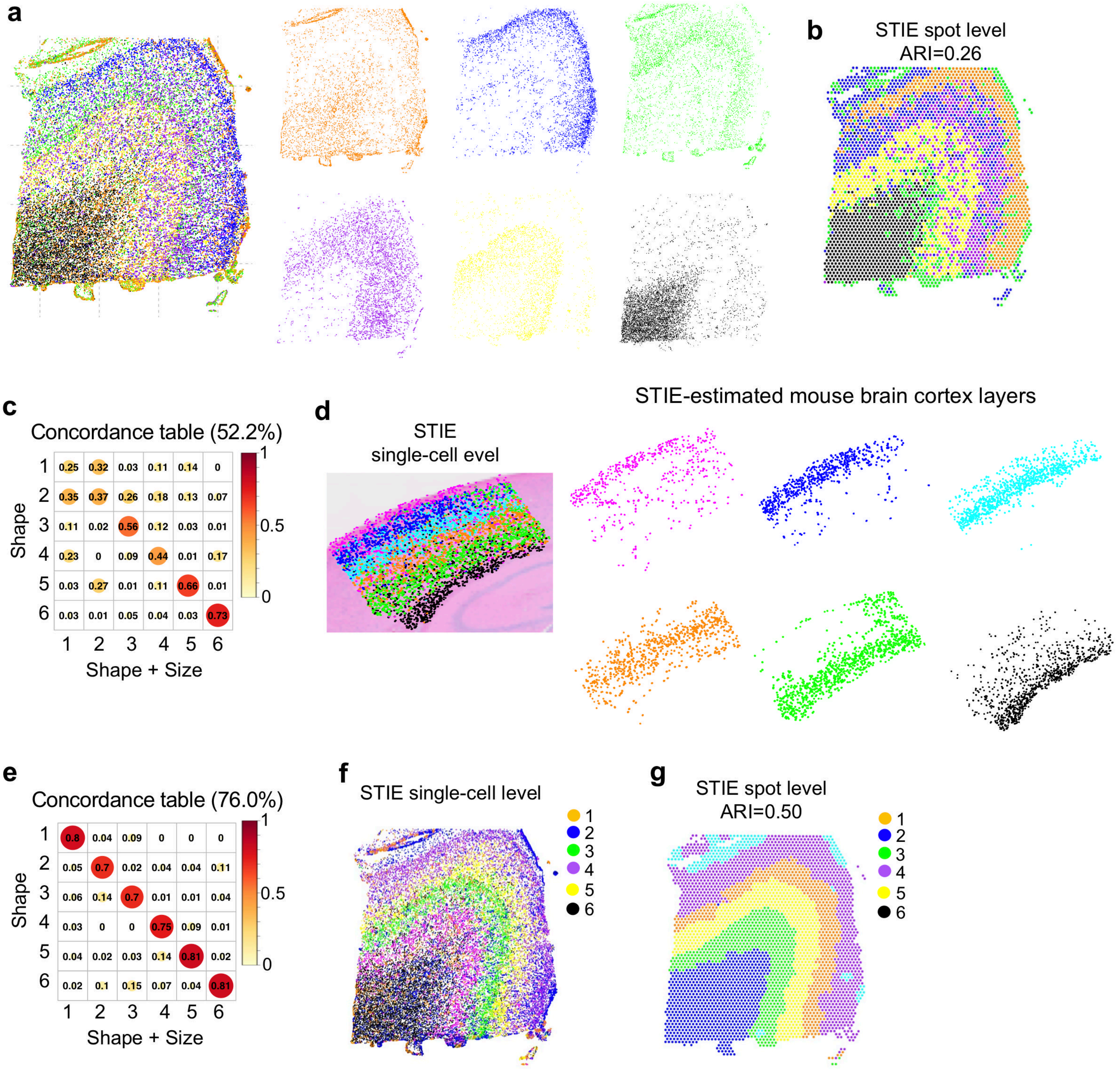

**Supplementary Figure 30.** Comparison of nuclear segmentation between the human brain frozen section (**a**) and mouse brain FFPE section (**b**). The nuclear segmentation on the frozen section may misclassify the nucleus and cytoplasm, while the nuclear segmentation on mouse brain FFPE identified the nuclear boundary more accurately.

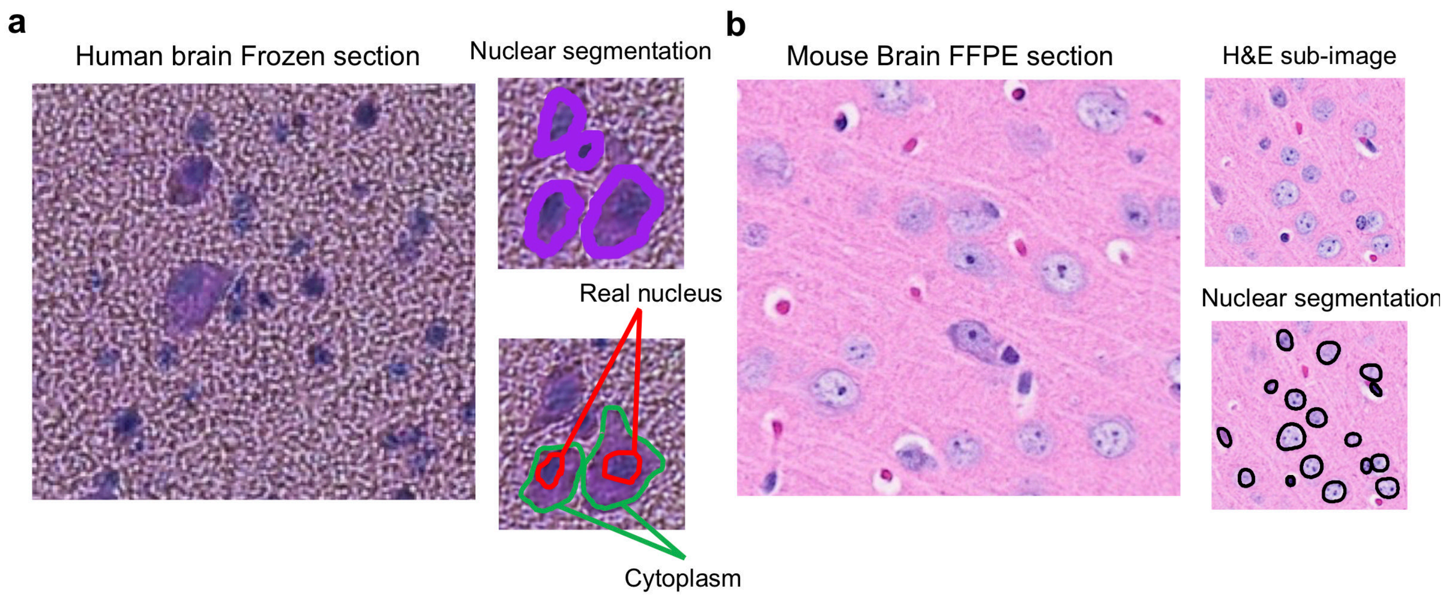

**Supplementary Figure 31.** Spot level K-means clustering on 10X Visium human breast cancer FFPE.

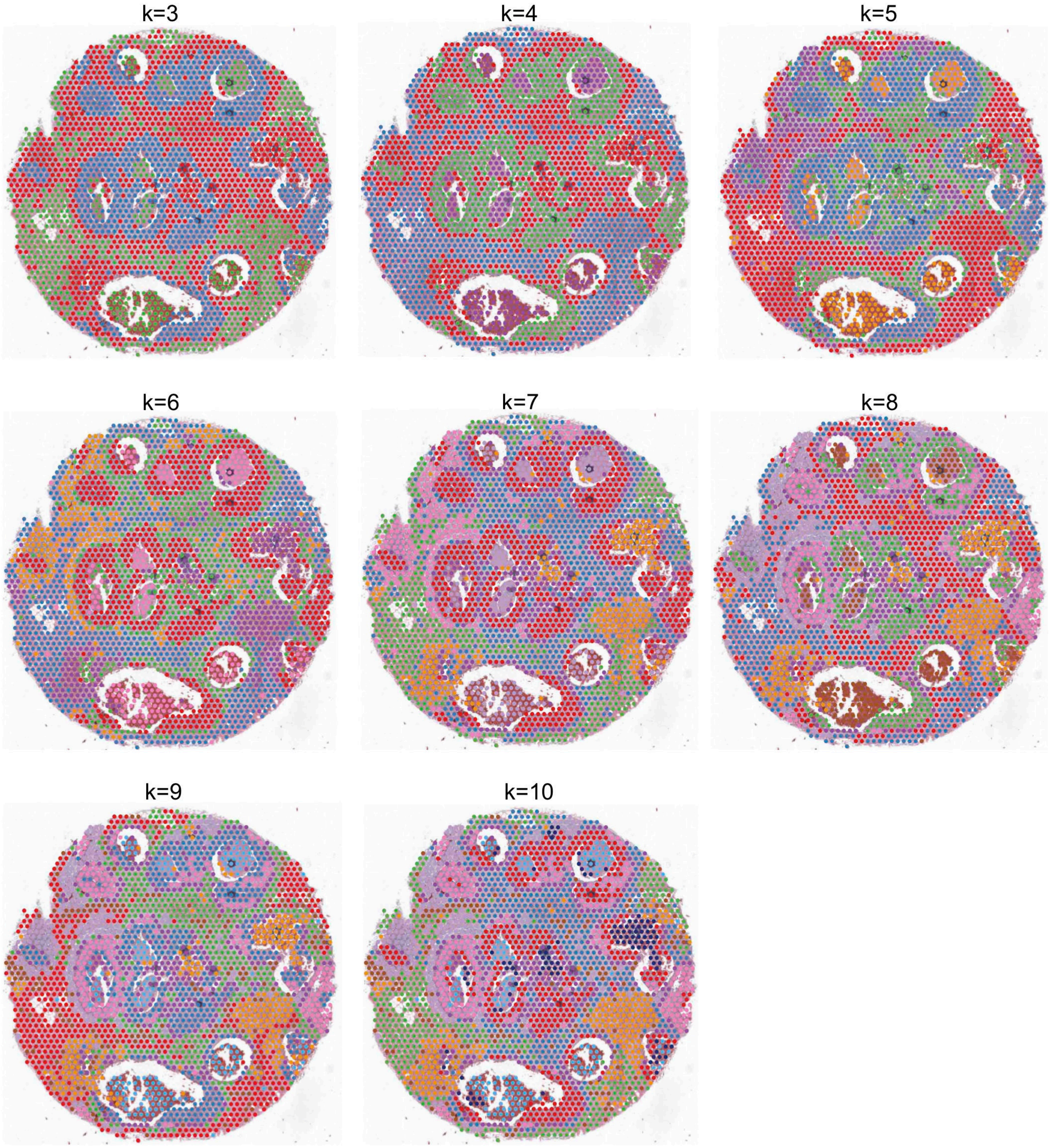

**Supplementary Figure 32.** Spot level SpaGCN clustering on 10X Visium human breast cancer FFPE.

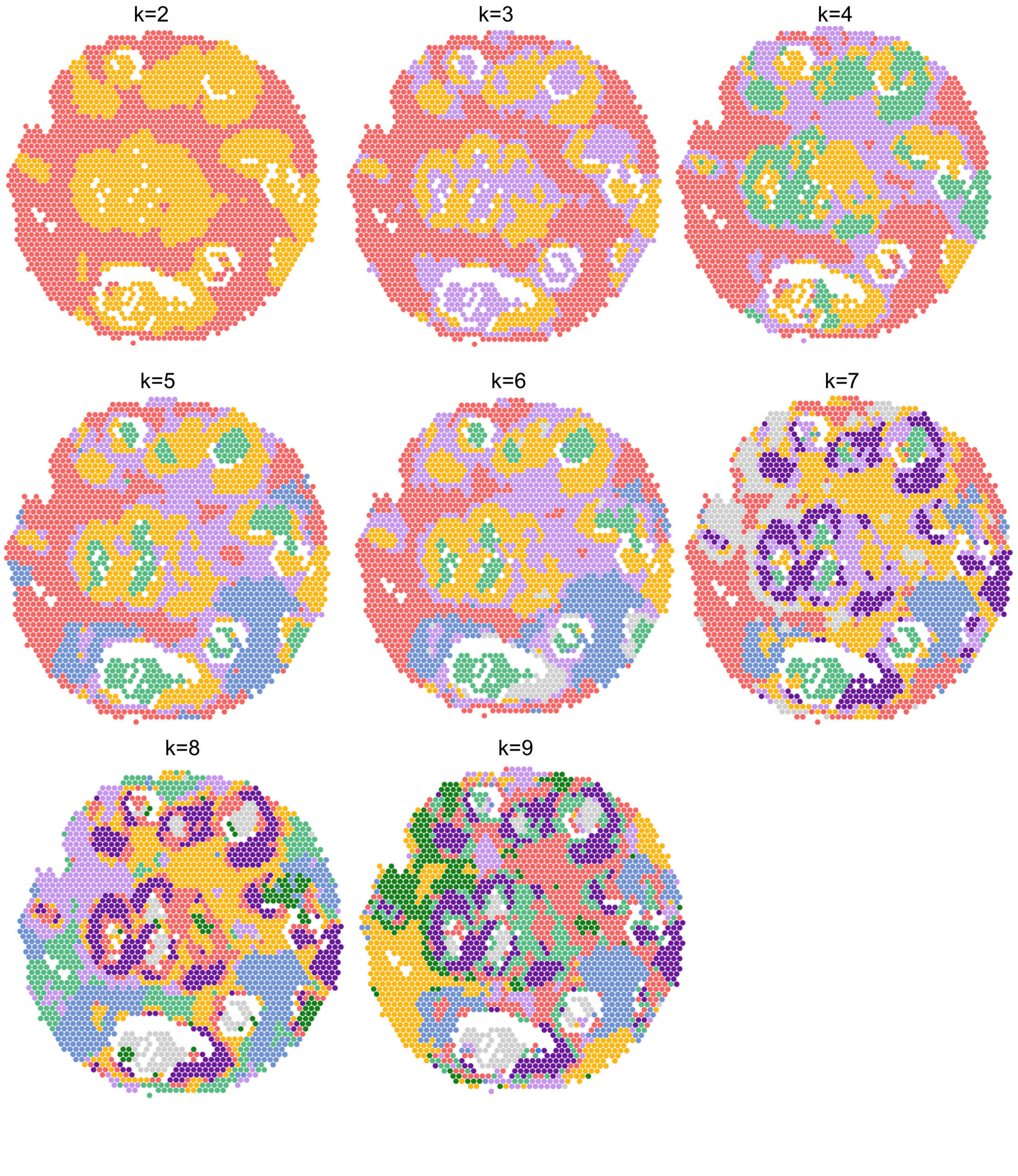

**Supplementary Figure 33.** Subspot level BayesSpace clustering on 10X Visium human breast cancer FFPE.

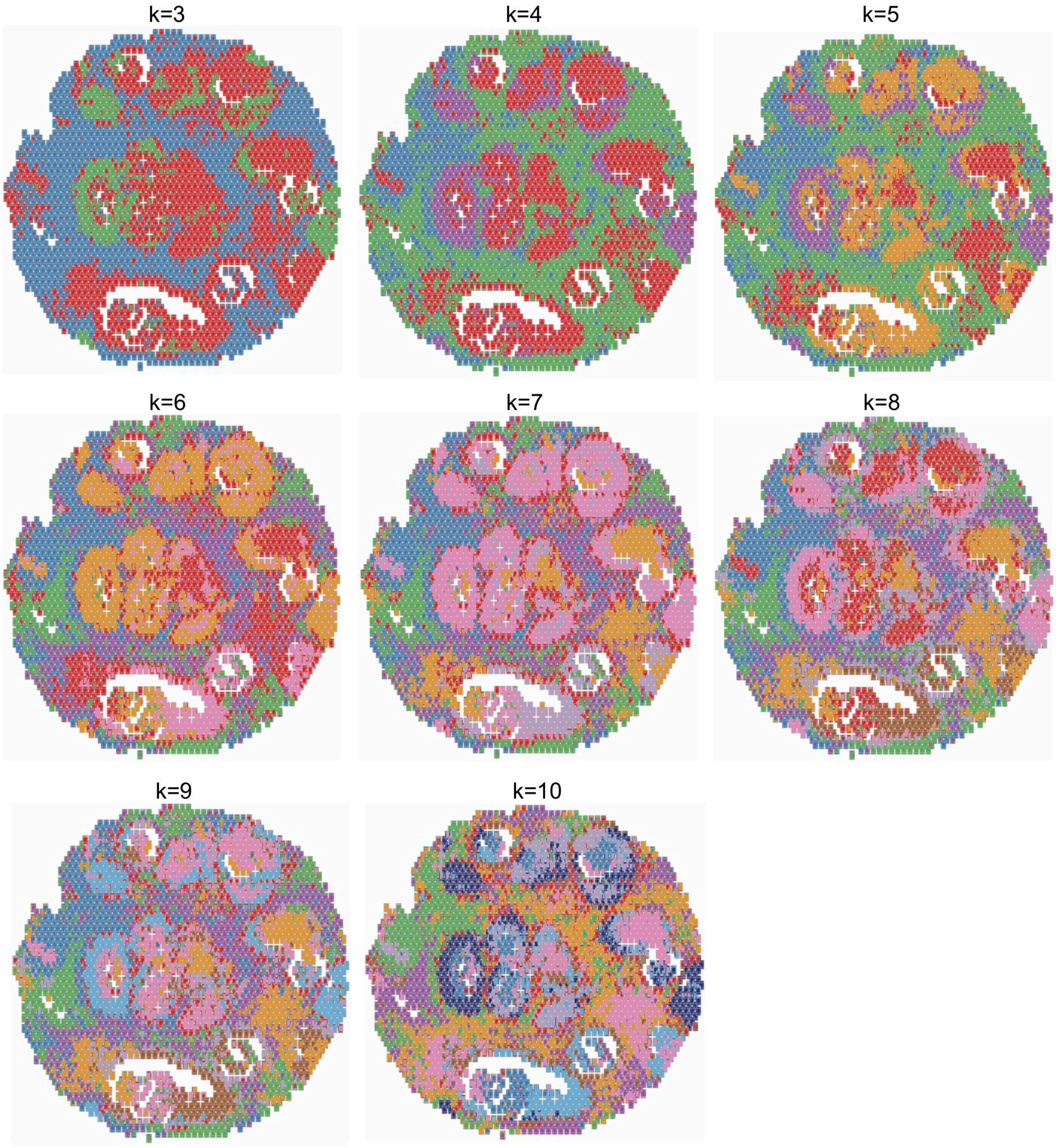

**Supplementary Figure 34.** Single-cell level STIE clustering on 10X Visium human breast cancer FFPE.

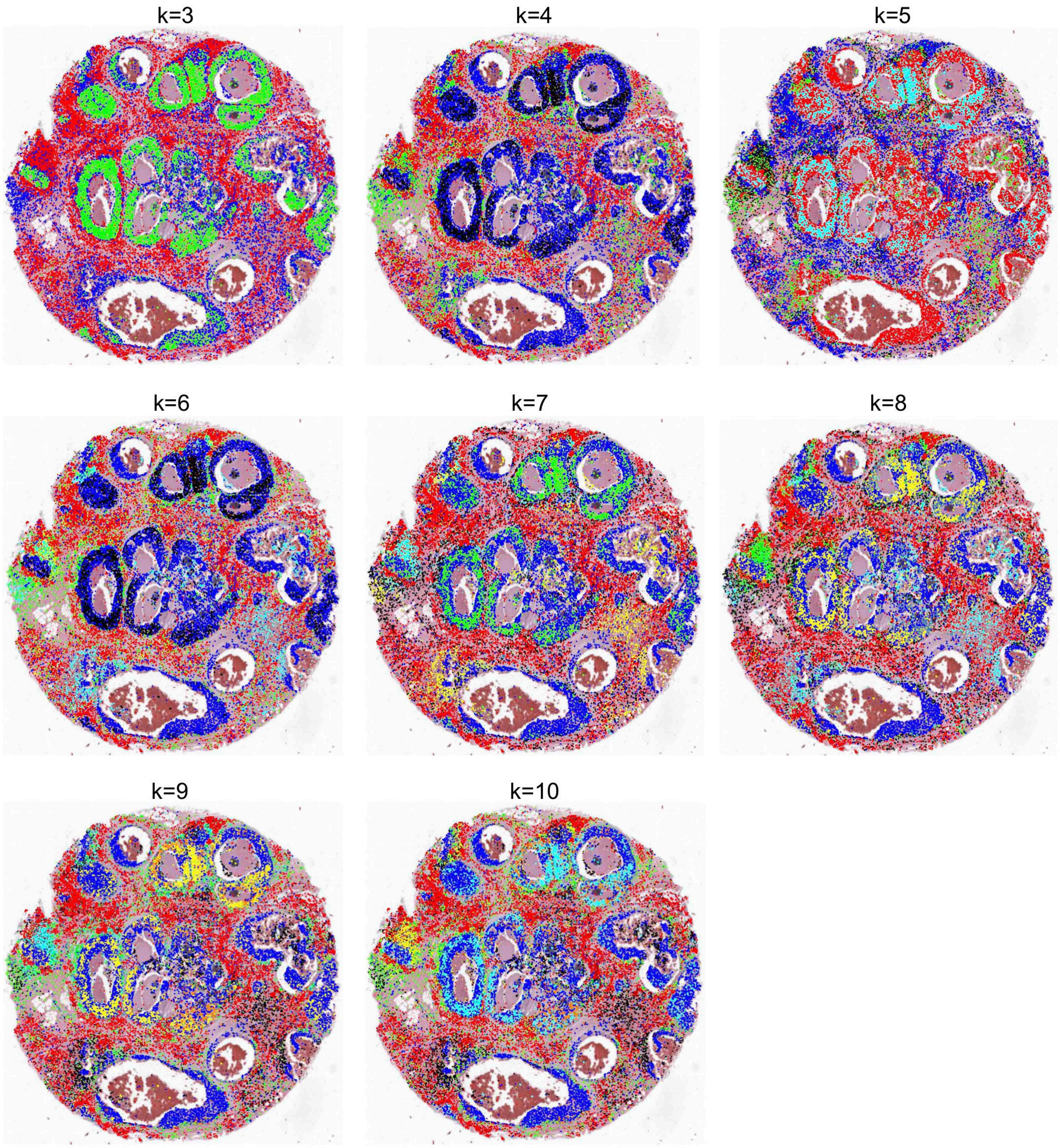

**Supplementary Figure 35.** Consistency of the tumor region between the clustering and the ground truth measured by Adjusted Rand Index (ARI) (**a**) and R2 (**b**).

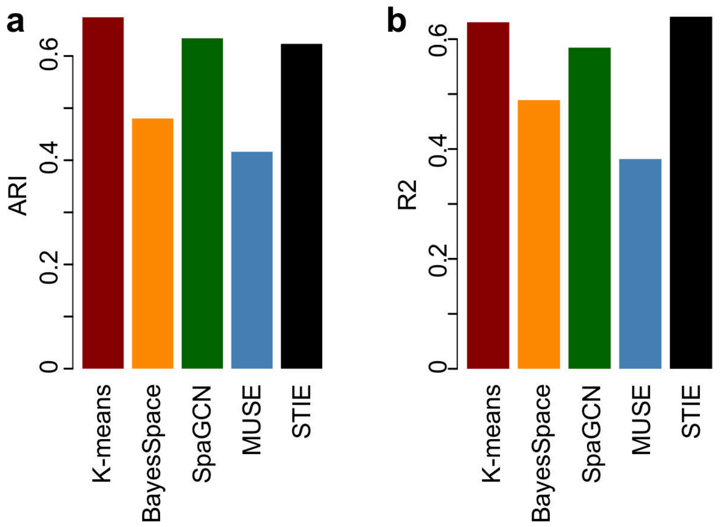

**Supplementary Figure 36. a,** The K-means and BayesSpace clustering on 10X Visium human breast cancer FFPE at k=7. **b,** The CAGEs by K-means and BayesSpcace at k=7, were deconvoluted based on the scRNA-seq derived transcriptomic signatures using NNLS.

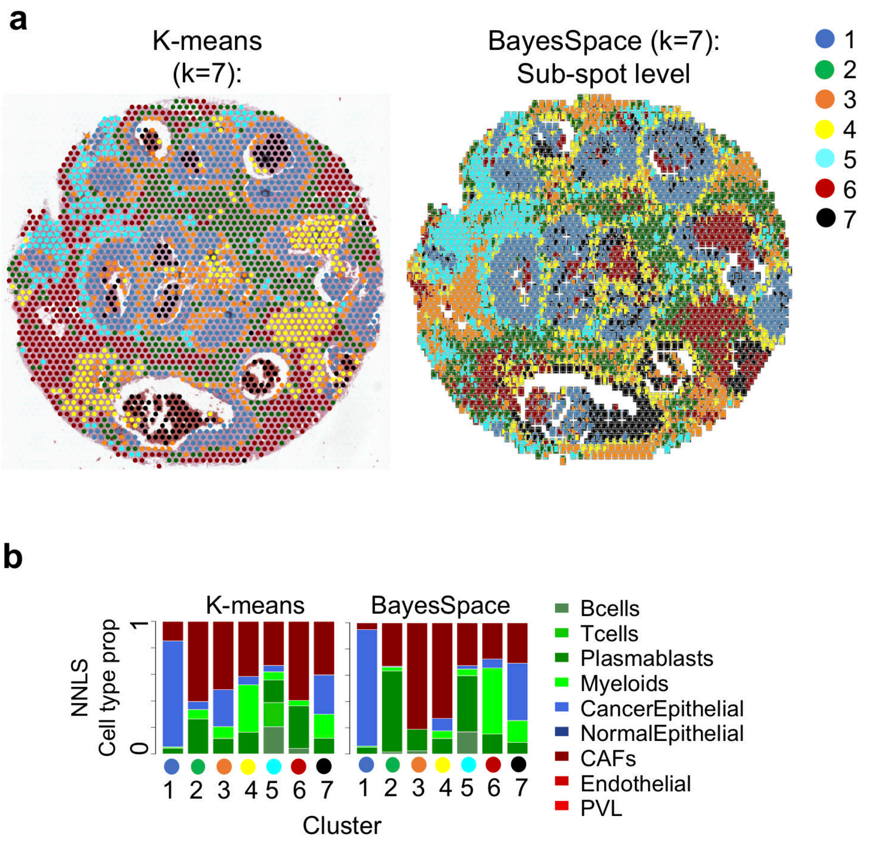

**Supplementary Figure 37.** The UMAP plot of human breast cancer scRNA-seq data from 26 primary tumors. **a,** The original cell typing of 10,060 single cells. **b,** Cells that are mapped to the six STIE clusters as well as the unmapped ones (black).

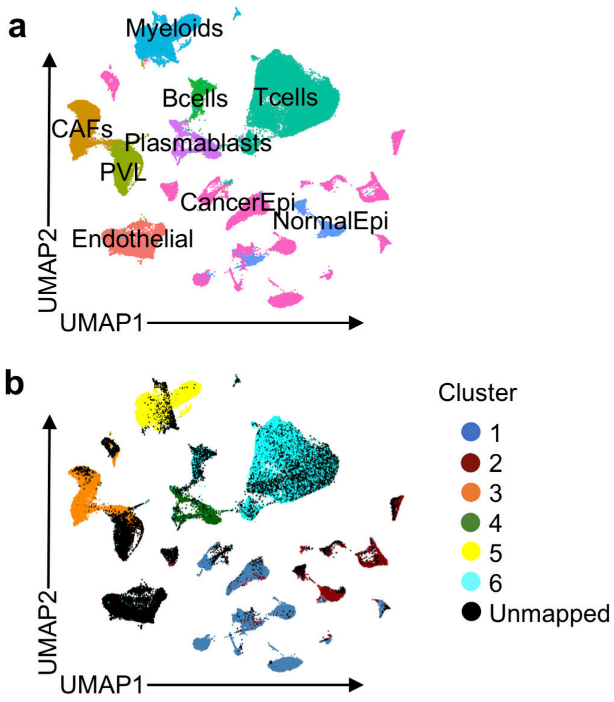

**Supplementary Figure 38.** The K-means and STIE clustering (at k=5) on the simulated high-resolution spot (5μm spot diameter) spatial transcriptomics data of mouse brain hippocampus. The CAGE by K-means and STIE were deconvoluted based on the scRNA-seq derived transcriptomic signatures using both NNLS and DWLS.

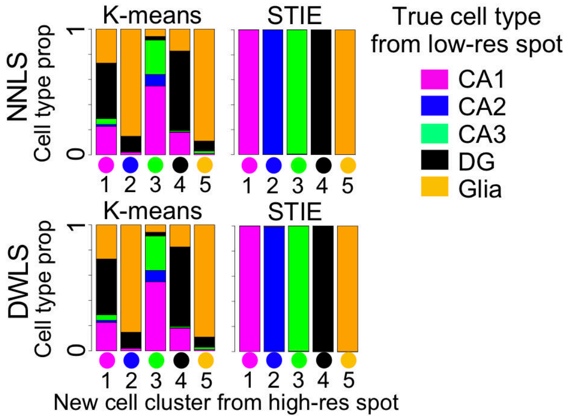

**Supplementary Figure 39. a,** The K-means and STIE clustering (at k=6) on the simulated high-resolution spot (5μm spot diameter) spatial transcriptomics data of human breast cancer. **b,** The CAGEs by K-means and STIE were deconvoluted based on the scRNA-seq derived transcriptomic signatures using both NNLS and DWLS.

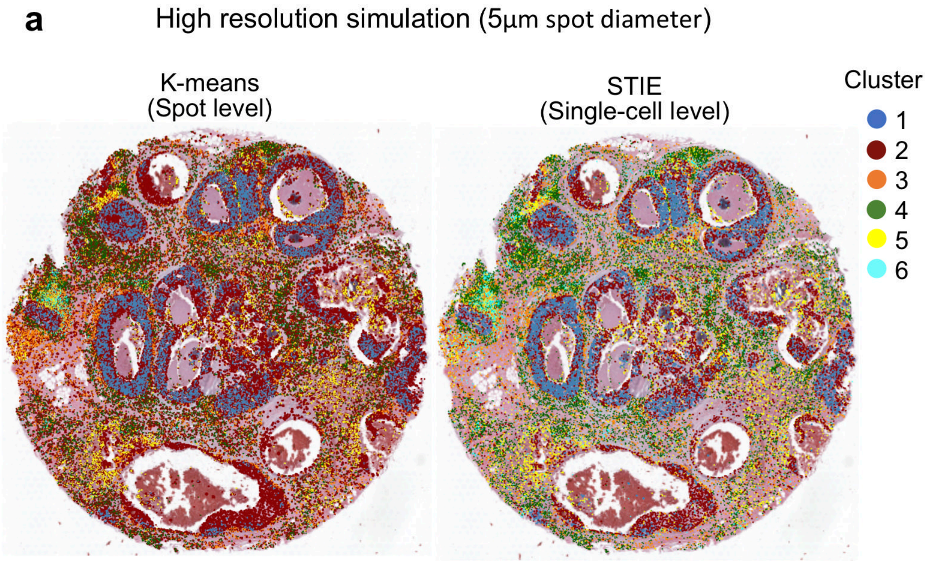


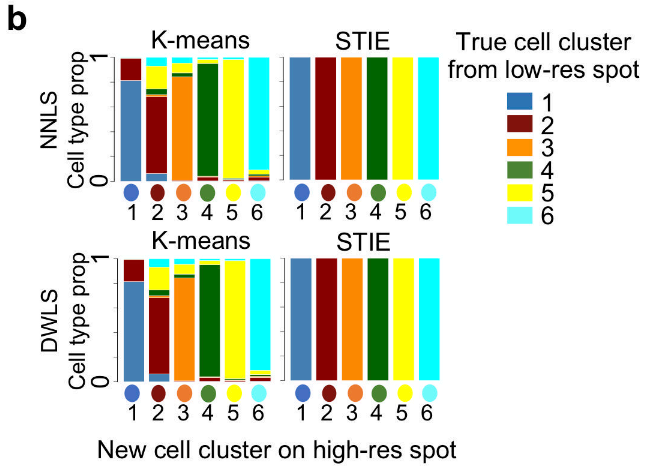

**Supplementary Figure 40.** Spot level MUSE clustering on the simulated high-resolution spot spatial transcriptomics data of mouse brain hippocampus (**a**) and human breast cancer (**b**).

**a** MUSE on mouse brain hippocampus high-res simulated spot (50 clusters)

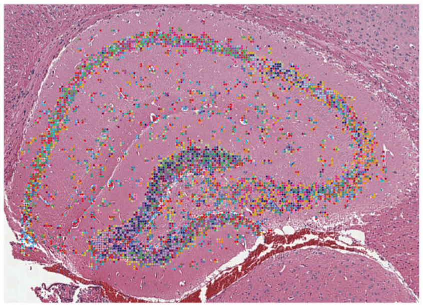

**b** MUSE on human breast cancer high-res simulated spot (159 clusters)

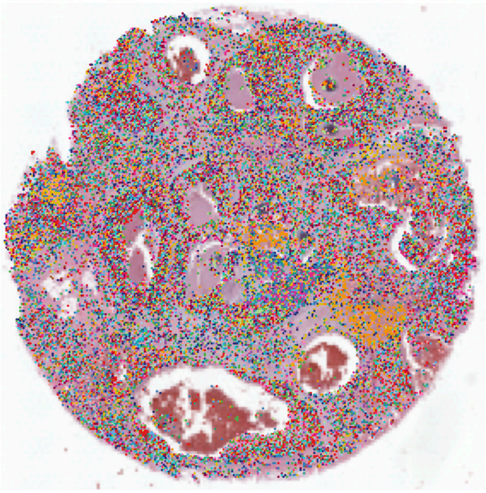

**Supplementary Figure 41**. Evaluation of RMSE of STIE clustering over different λ on the 10X Visium mouse brain cortex (**a**), 10X Visium C2 Chemistry CytAssist mouse brain cortex section 1 (**b**), and section 2 (**c**).

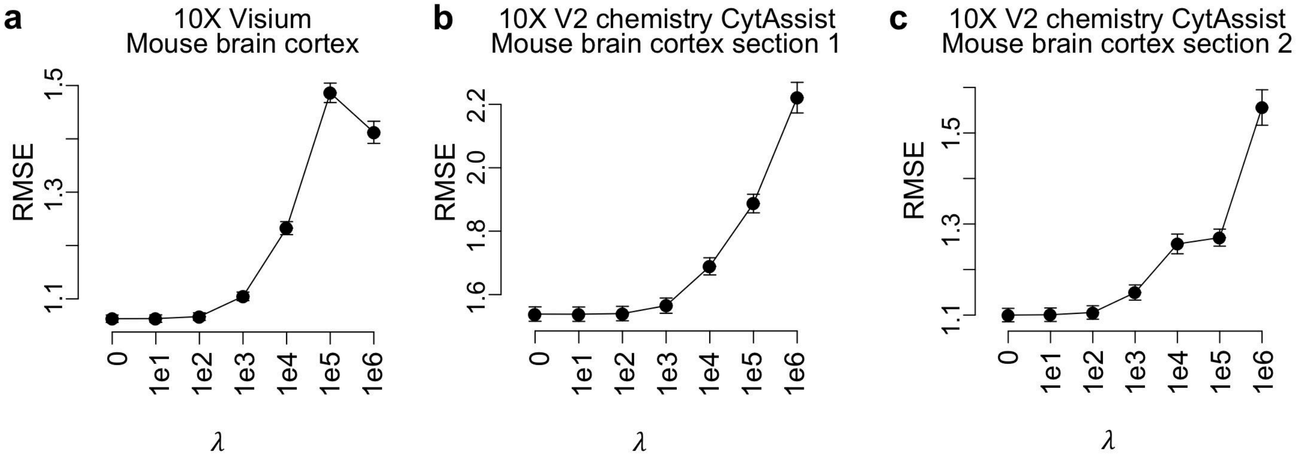

**Supplementary Figure 42.** The boxplot of human breast cancer cell-type colocalization detected by SPOTlight, DWLS, Stereoscope, RCTD, Tangram, BayesPrism, and STIE.

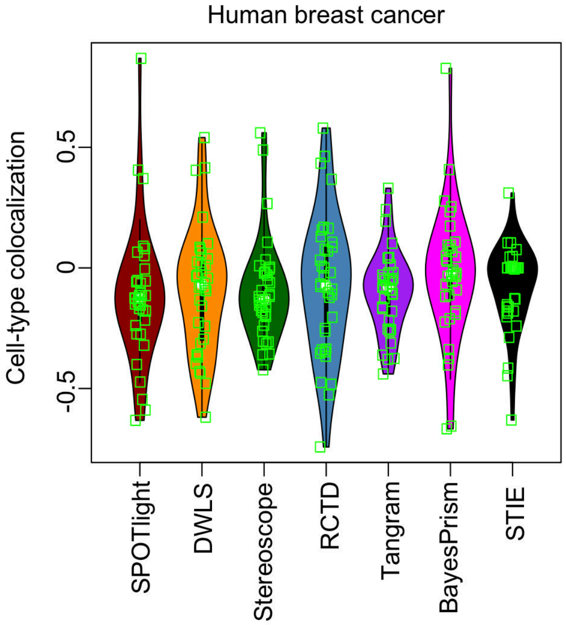

**Supplementary Figure 43.** Heatmap of the cell-cell interaction strength of pathways in sender cell types (left) and receiver cell types (right) at λ=1e4 for STIE on the human breast cancer. In sender and receiver heatmaps, the bar plots on top and right represent the total amount of cell-cell interaction within the cell type and pathway, respectively.

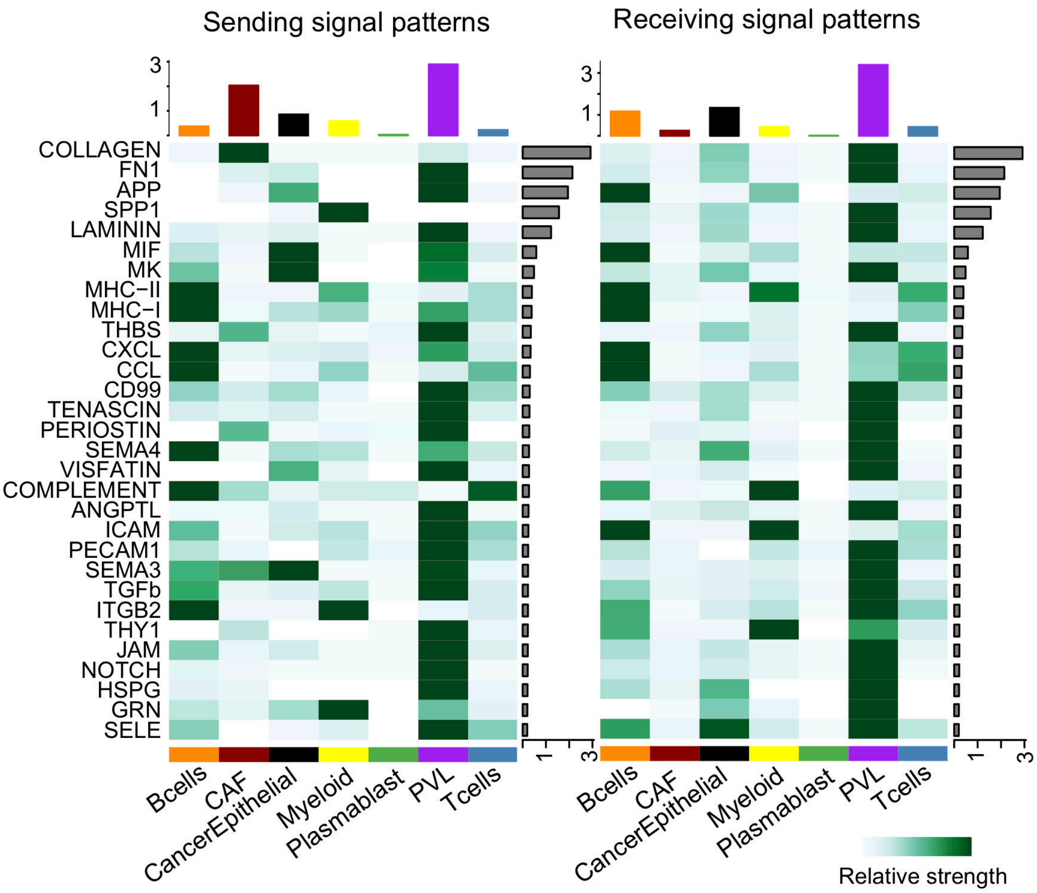

**Supplementary Figure 44.** The plot of Cophenetic and Silhouette values to select the number of latent sending (**a**) and receiving (**b**) patterns to drive the cell-cell communication at λ=1e3 for STIE on the human breast cancer. Both Cophenetic and Silhouette metrics measure the stability for a particular number of patterns based on a hierarchical clustering of the consensus matrix. The Cophenetic and Silhouette values begin to drop suddenly when the number of sending and receiving patterns are 3, suggesting the suitable number of patterns.

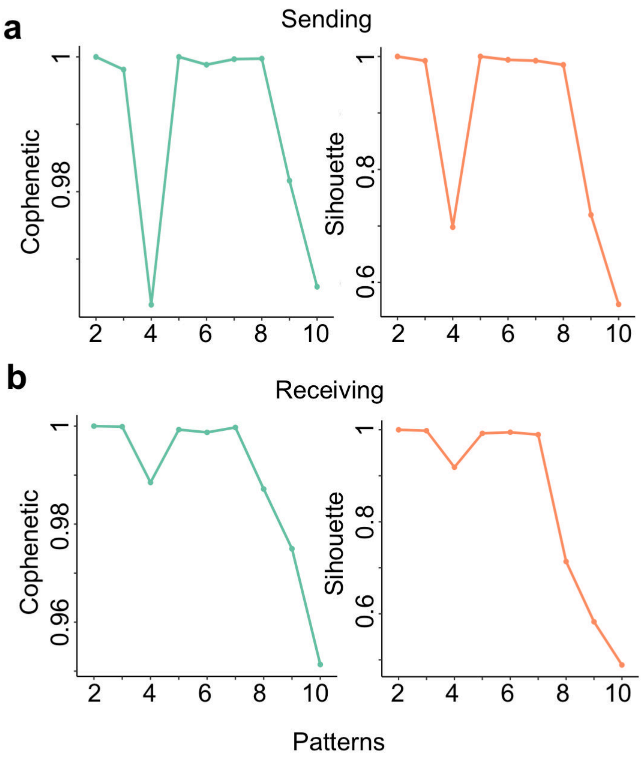

**Supplementary Figure 45.** The heatmap of contribution of pathways (up) and cell types (bottom) to the pattern of senders and receivers (at λ=1e3 for STIE on the human breast cancer). **a**, The sending patterns at k=3. **b**, The receiving patterns at k=3.

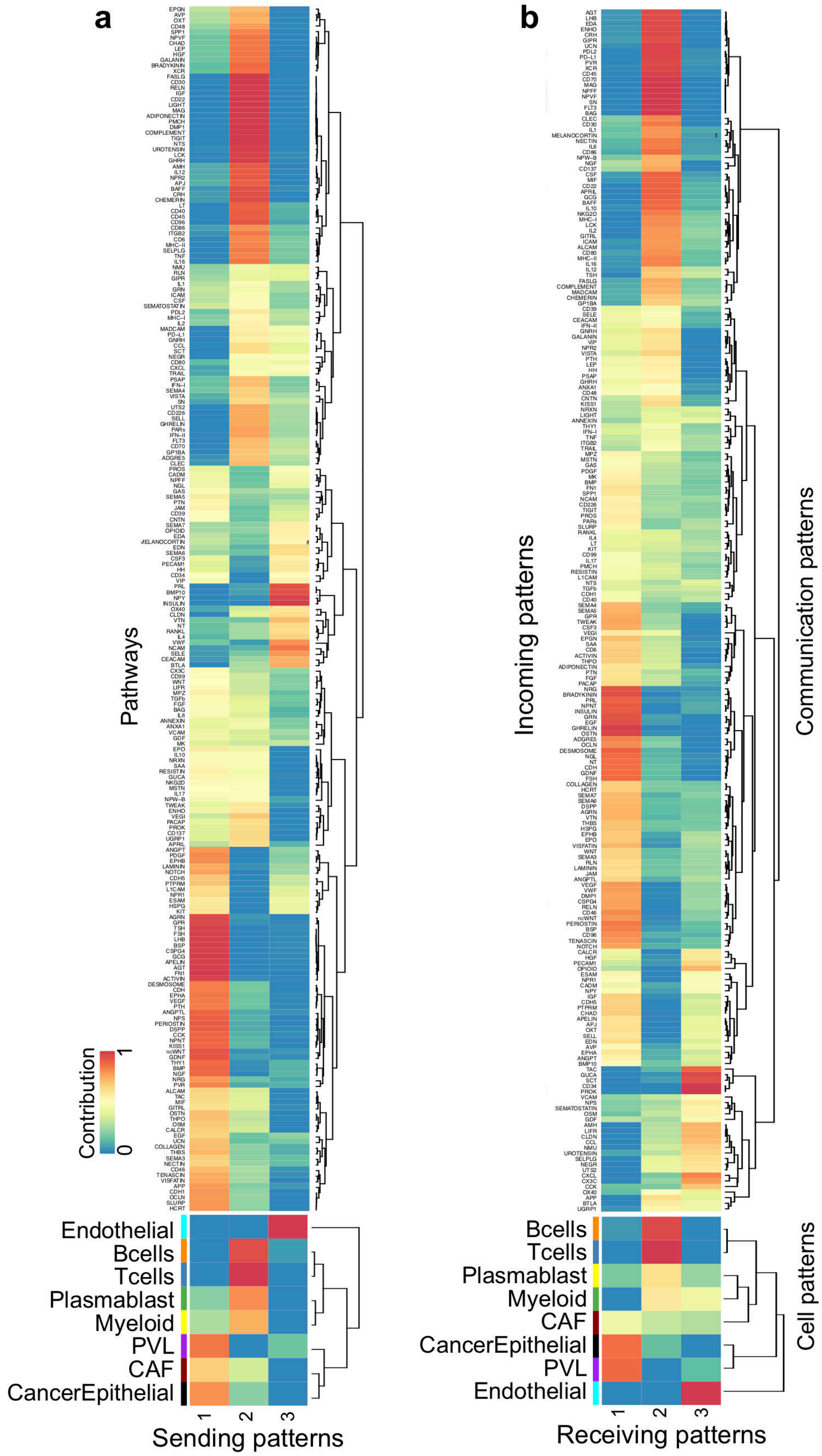

**Supplementary Figure 46.** The plot of Cophenetic and Silhouette values to select the number of latent sending (**a**) and receiving (**b**) patterns to drive the cell-cell communication at λ=1e4 for STIE on the human breast cancer. We chose 2 patterns for both sending and receiving signals for the following analyses.

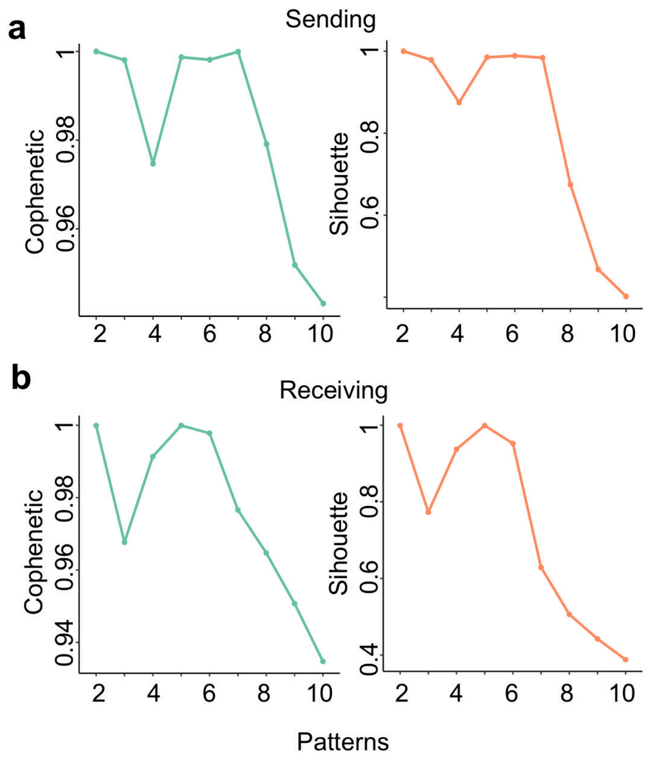

**Supplementary Figure 47.** The heatmap of contribution of pathways (up) and cell types (bottom) to the pattern of senders and receivers (at λ=1e4 for STIE on the human breast cancer). **a**, The sending patterns at k=2. **b**, The receiving patterns at k=2.

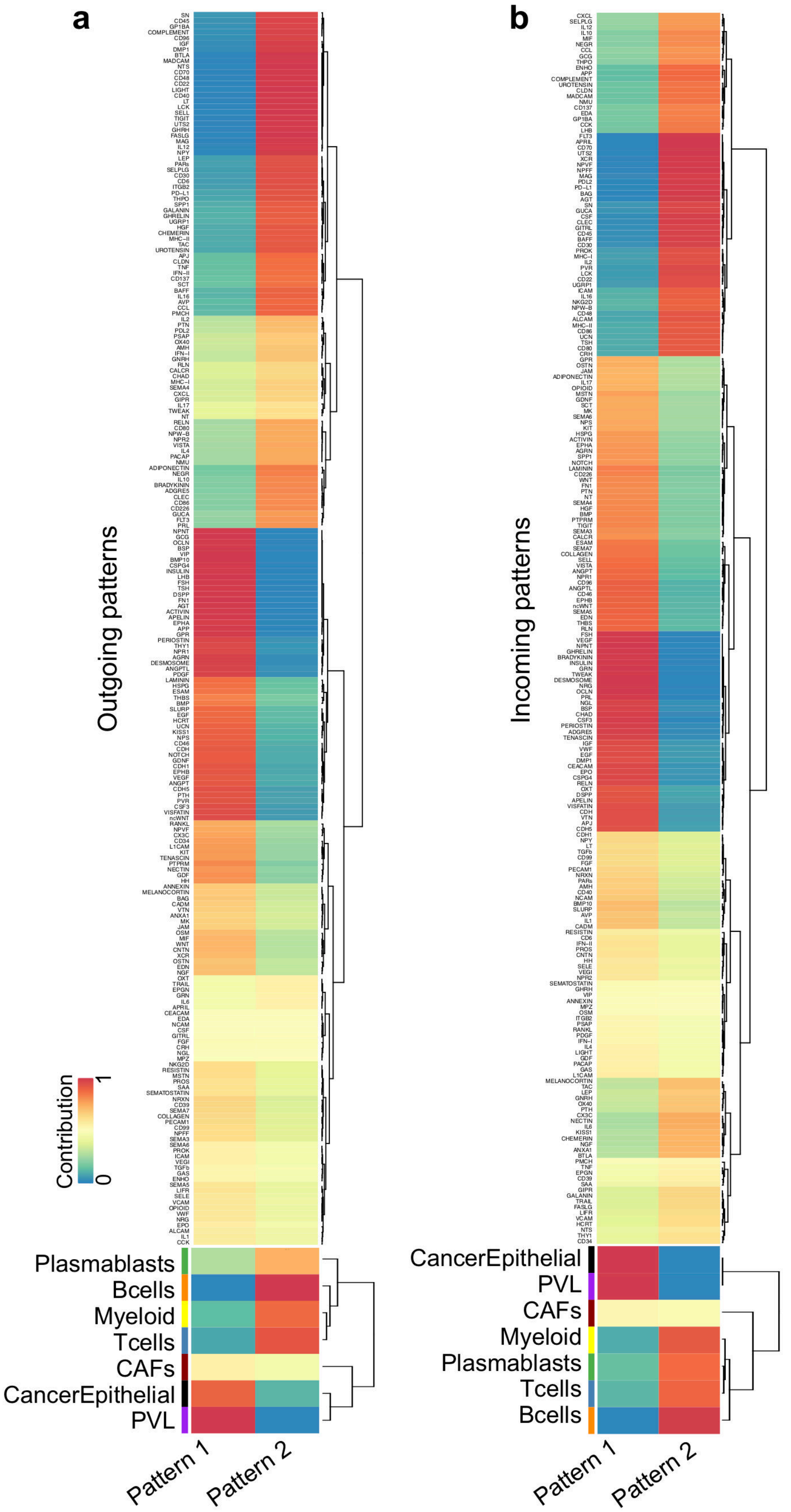

**Supplementary Figure 48.** The sender and receiver cell-cell interaction patterns across the tissue (at λ=1e4 for STIE on the human breast cancer). The color of the edge is indicated by the sender cell type and the receiver cell type in the sender and receiver patterns, respectively.

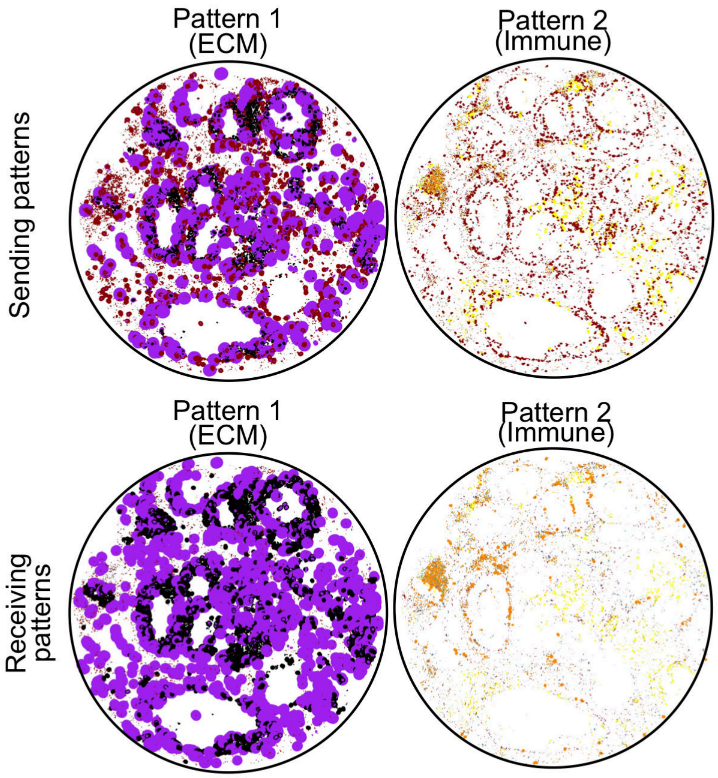

**Supplementary Figure 49.** The plot of Cophenetic and Silhouette values to select the number of latent sending (**a**) and receiving (**b**) patterns to drive the cell-cell communication (at λ=0 for STIE on the mouse brain hippocampus). Both metrics begin to drop suddenly when the number of sending and receiving patterns are 5, suggesting the suitable number of patterns .

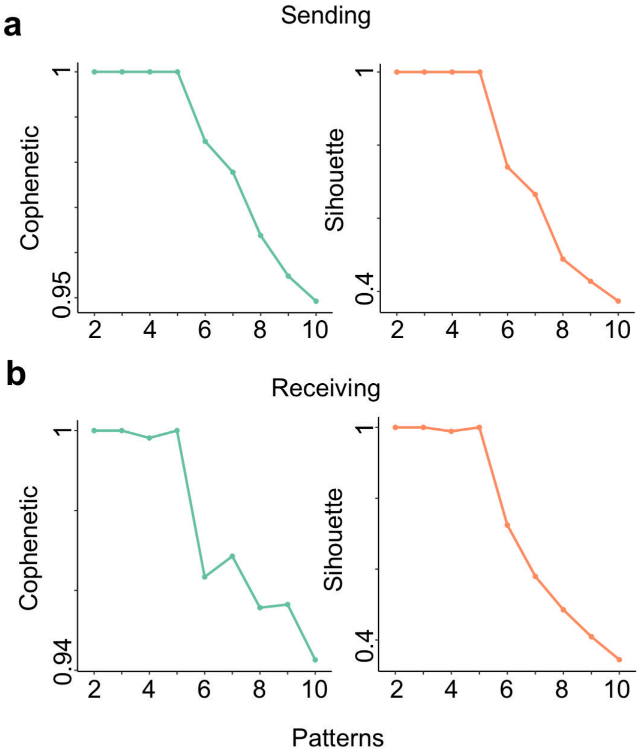

**Supplementary Figure 50.** The heatmap of contribution of pathways (up) and cell types (bottom) to the pattern of senders and receivers (at λ=0 for STIE on the mouse brain hippocampus). **a**, The sending patterns at k=5. **b**, The receiving patterns at k=5.

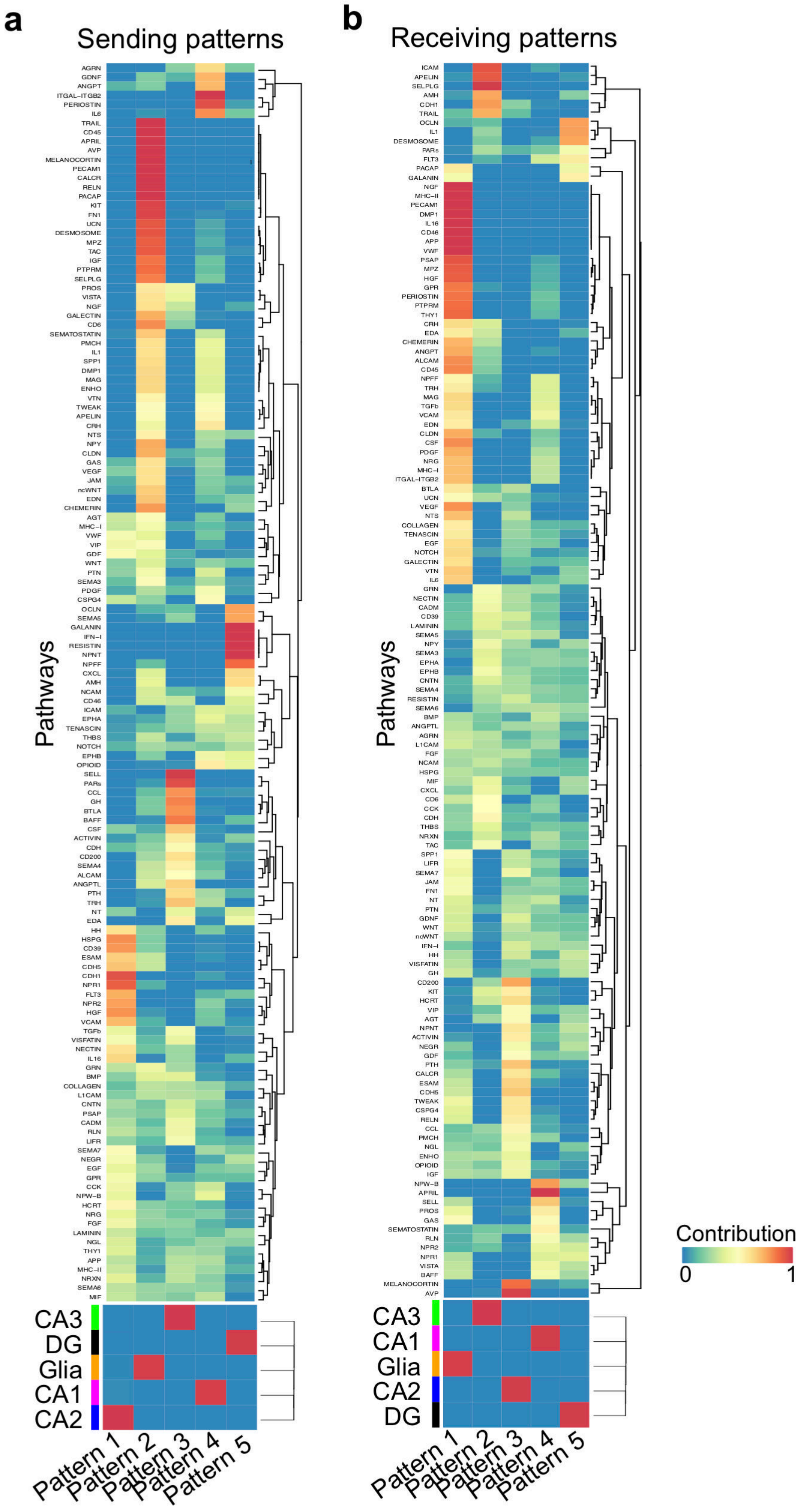

**Supplementary Figure 51.** The spatially resolved cell type interaction across the tissue (at λ=0 for STIE on the mouse brain hippocampus). **a**, The sending patterns at k=5. **b**, The receiving patterns at k=5.

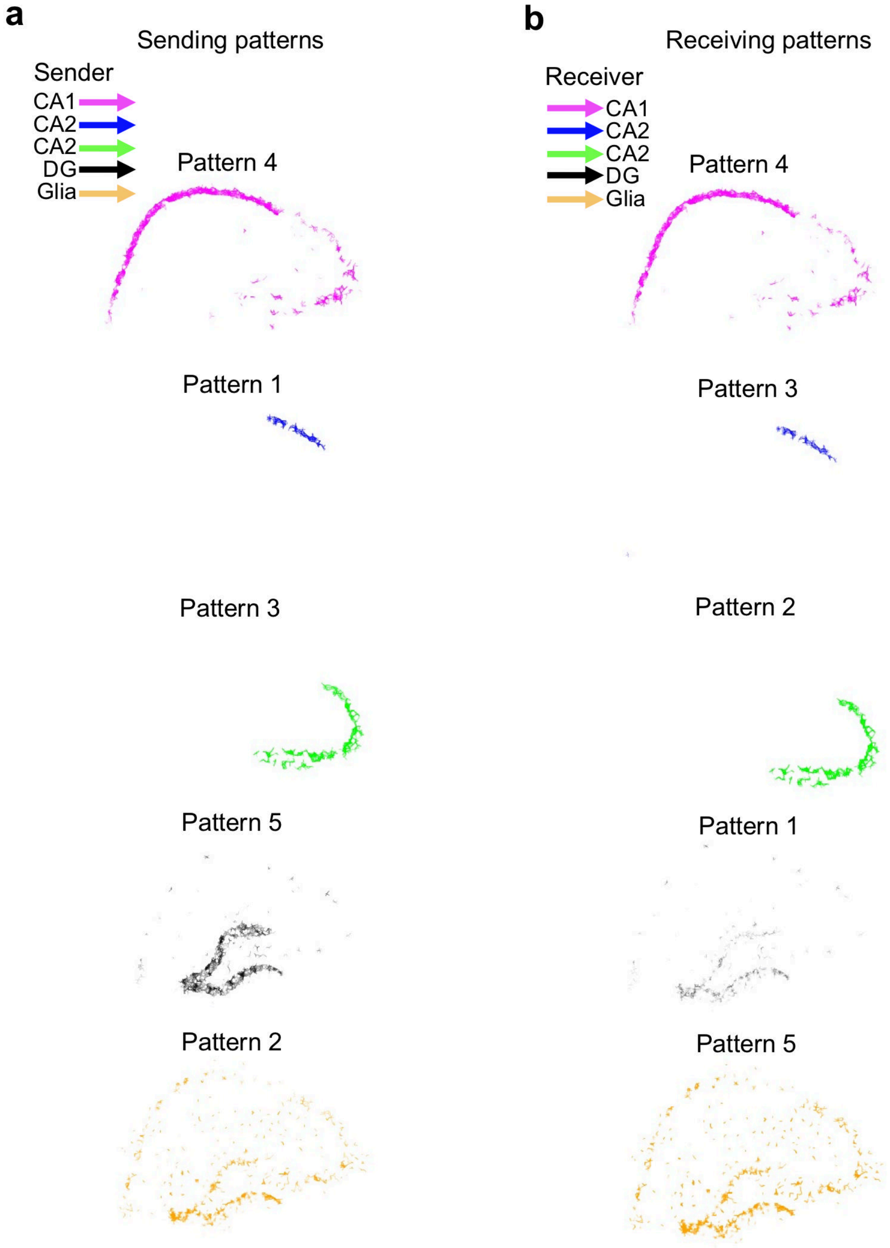
